# Phylogenetic estimation of diversity-dependent biogeographic rates using deep learning

**DOI:** 10.64898/2026.02.17.706216

**Authors:** Albert C. Soewongsono, Michael J. Landis

## Abstract

Ecological theory predicts that local species richness can influence biogeographic rates of speciation, extinction, and dispersal. For instance, increasing the number of competing species within a region may cause local speciation and dispersal rates to decrease but local extinction rates to increase, inducing a carrying capacity for local species richness. In this article, we introduce a fully generative, event-based phylogenetic diversification model, called DDGeoSSE, that allows diversity-dependent effects of local species richness to modulate biogeographic rates of diversification and range evolution. DDGeoSSE can accommodate and test a variety of alternative diversification scenarios that involve positive, negative, and neutral interactions among sympatric species for speciation, extinction, and dispersal. We derive mathematical and statistical properties of biogeographic outcomes generated by this model, such as the carrying capacity for a clade at equilibrium, which we validate through simulation. Because diversity-dependent phylogenetic models typically do not have tractable likelihood functions, we use deep learning with phyddle to perform parameter inference and model selection. Separately applying DDGeoSSE to Caribbean *Anolis* lizards and cloud forest-dwelling *Viburnum* plants, we find evidence that local species richness plays a significant role in shaping diversification dynamics for both clades.

## Introduction

Biologists have long puzzled over two interrelated problems concerning spatial patterns of biodiversity. On one hand, species richness is unevenly distributed in space, with some regions containing far more species than others (Gaston, 2000; Hillebrand, 2004; Mittelbach et al., 2007). On the other hand, species richness at both local and global scales does not increase indefinitely (Van Valkenburgh, 1999; Phillimore and Price, 2008; Pires et al., 2017), where ecological forces, such as competition, are predicted to limit unbounded growth (Macarthur and Wilson, 1967; van Valen, 1973). The exceptionally clear fossil record for marine invertebrates also documents that taxon richness varies among habitats (Bambach, 1977) and latitudes (Jablonski et al., 2006) over time. In addition, periods of higher richness tend to instigate faster extinction and slower origination rates whereas, inversely, faster origination and slower extinction rates accompany periods of lower richness (Alroy, 2008). Learning what dynamics govern the gain, loss, and maintenance of species diversity in space, particularly over long timescales, is therefore a question of central importance to ecologists and evolutionary biologists alike.

To study diversification, researchers have designed a variety of phylogenetic models that describe how clades accumulate species richness with time. Beginning with the pioneering work in the 1990s by Sean Nee and collaborators (Nee et al., 1992, 1994), myriad phylogenetic birth-death models have been developed over the last several decades (Stadler, 2013; Morlon et al., 2024; Title et al., 2026) that variously allow diversification rates to vary over time, across lineages, and in response to species age (Rabosky and Lovette, 2008; Hagen et al., 2015; Soewongsono et al., 2022), inherited traits (Maddison et al., 2007; FitzJohn, 2010; Beaulieu and O’Meara, 2016), and environmental conditions (Condamine et al., 2013; Mazet et al., 2023). One such model is the Geographic State-dependent Speciation-Extinction (GeoSSE) process (Goldberg et al., 2011), which models how speciation, extinction, and dispersal rates are shaped by the geographical range of each species as it evolves. Since then, researchers have further improved GeoSSE to allow unobserved (hidden) characters (Caetano et al., 2018), regional features (Landis et al., 2022), or both (Quintero et al., 2023), to modulate biogeographic rates.

Despite these advances, most existing diversification models allow species richness to increase exponentially with time, contradicting predictions made by the equilibrium theory of island biogeography (Macarthur and Wilson, 1967): that, all else being equal, local increases in species richness should tend to accelerate extinction while decelerating speciation and immigration, inducing an equilibrium level of species richness when all rates sum to zero. However, models of island biogeography are typically deterministic (no randomness of the timing and placement of biogeographic events), phenomenological (island area is used as a proxy to govern competition, rather than actual numbers of species), and non-phylogenetic (they do not use any explicit representation of shared common ancestry when considering historical dynamics).

Relatively few investigators have designed diversity-dependent phylogenetic models of biogeography for measuring carrying capacities (reviewed in Morlon et al., 2024; Title et al., 2026). Rabosky and Glor (2010) innovated a diversity-dependent extension to the Multistate Speciation-Extinction model (MuSSE; FitzJohn, 2012) that assumes each species occupy one island at any instant and models diveristy-dependence using a deterministic linear function of time to approximate the negative effect of species richness on diversification rates. The Dynamic Assembly of Islands through Speciation, Immigration and Extinction framework (DAISIE; Valente et al., 2015, 2017, 2020; Xie et al., 2023) models species assembly in island systems using diversity-dependent speciation and immigration rates that decrease linearly with the actual number of species present in the island system at a given time (Valente et al., 2015). DAISIE is ideally suited for mainland-island systems, as it does not model dispersal among islands, diversification dynamics within the mainland region, or the general existence of widespread species. DAISIE and the Rabosky-Glor model possess many desirable properties, such as the ability to forbid certain inter-island dispersal events (Rabosky and Glor, 2010) or to estimate carrying capacity parameters simultaneously from multiple clades (Valente et al., 2015, 2017). However, both frameworks disallow the full set of widespread species ranges, do not use the full set of biogeographic events allowed by GeoSSE, and assume all biogeographic processes (speciation, extinction, dispersal) are governed by a single carrying capacity parameter, which together hinder their ability to determine whether diversity has the same or different impacts on those three processes.

Building upon these previous advances, we designed a new Diversity-Dependent GeoSSE (DDGeoSSE) model that retains the flexibility of the GeoSSE framework while allowing local species richness to influence rates of evolution, either positively or negatively. Unlike existing models, our model explicitly accounts for the actual number of species present at any given time to separately modulate the rates for each process operating within (within-region speciation, extinction) or among (between-region speciation, dispersal) regions. In addition, rather than assuming the carrying capacity for a clade exists by treating it as a parameter in the model’s rate equations, the local level of equilibrium diversity (analogous to carrying capacity) emerges as an expectation from the data-generating DDGeoSSE process. As such, we derive theory to measure the equilibrium diversity under various diversity-dependent scenarios using alternative sets of DDGeoSSE rate parameters. DDGeoSSE is fully generative, meaning it can simulate phylogenetic trees and the evolution of species ranges under a variety of diversity-dependent scenarios. Thus, we can use the model to explore the relationships between phylogenetic and biogeographic patterns with diversity-dependent processes. Using simulations, we show that alternative DDGeoSSE scenarios produce distinct patterns of tree balance, branching times, and mean range sizes.

We were also interested in applying DDGeoSSE for inference tasks, such as estimating model parameters and performing model selection for biological datasets. However, as models become more complex, it becomes increasingly challenging to derive corresponding likelihood functions that are analytical and tractable (Xie et al., 2023), where DDGeoSSE is no exception. This presents a problem, as likelihood-based estimation (e.g., standard maximum likelihood and Bayesian inference) typically requires an explicit likelihood function under the given model. Fortunately, deep learning has shown promising results as an alternative tool for likelihood-free inference, particularly in phylodynamic applications (Voznica et al., 2022; Lambert et al., 2023; Thompson et al., 2024; Qin et al., 2026). Using simulated datasets and a deep learning pipeline implemented in the phylogenetics software phyddle (Landis and Thompson, 2025), we trained several neural networks to perform parameter estimation and model selection under different DDGeoSSE model variants. We then analyzed two empirical datasets with our trained networks: (1) the phylogeny of 158 *Anolis* lizard species occupying Caribbean island regions (Poe et al., 2017), and (2) the phylogeny of 38 species of cloud forest-dwelling *Viburnum* plants in the *Oreinotinus* clade (Donoghue et al., 2022). Both of these systems represent well-studied biogeographic radiations suitable for testing diversity-dependent diversification and dispersal dynamics.

## Methods

### Orientation

We introduce the DDGeoSSE model, which allows local species counts to influence regional rates of speciation, extinction, and dispersal. These diversity-dependent event rates can create distinct phylogenetic and biogeogeographic patterns, such as regional turnover events, as illustrated in Figure 1. For readers unfamiliar with GeoSSE, we provide a description of the model in the Supplement (Supp. Fig. S1). Here, we briefly explain how the DDGeoSSE model behaves, and then outline how the Methods section is structured.

**Figure 1:**
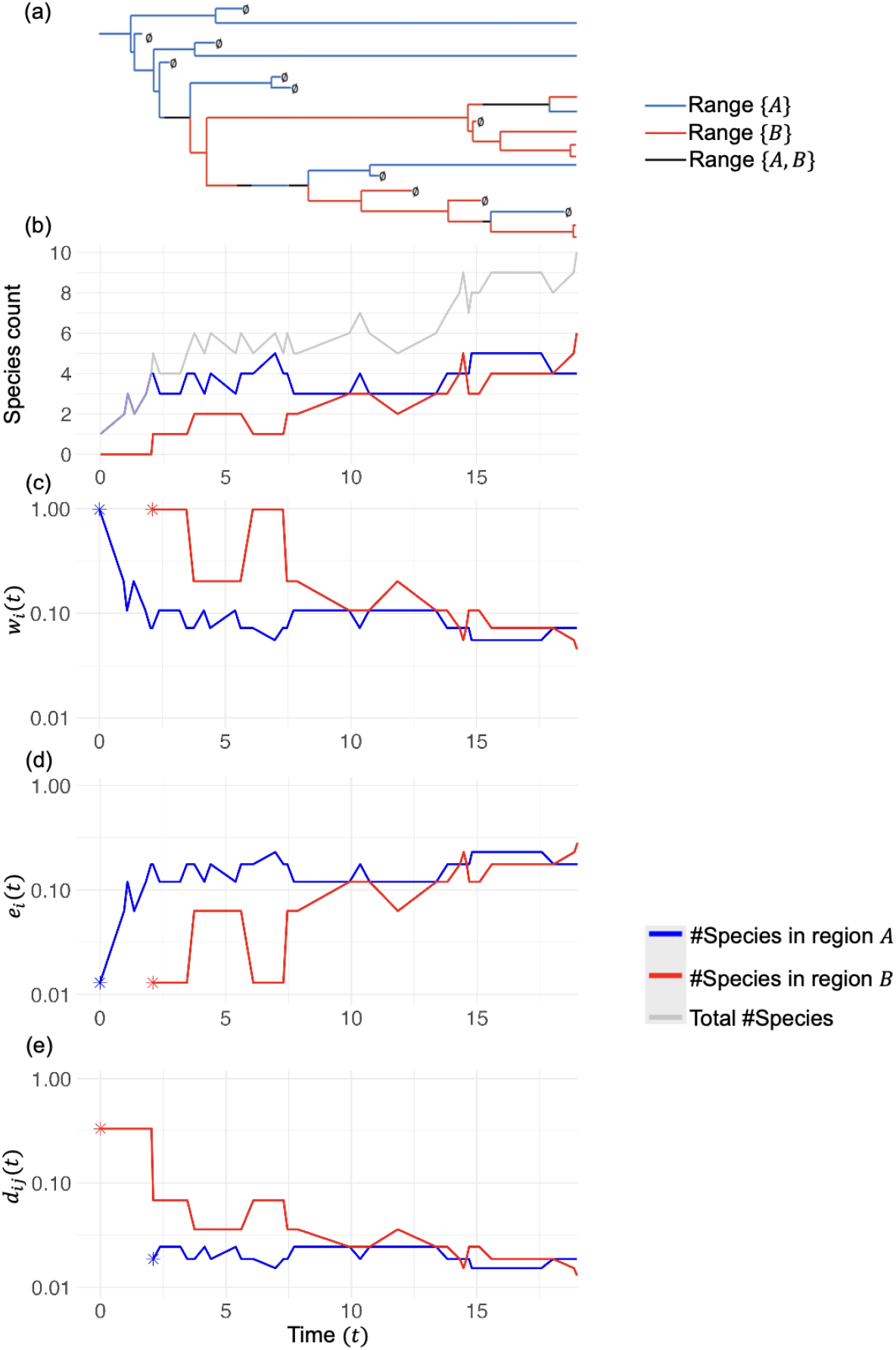
Cartoon of Logarithmic DDGeoSSE model dynamics. (a) Phylogenetic tree with extant taxa that evolved among regions *A* and *B*. Blue represents species that only occupy region *A*, red represents species that only occupy region *B*, black represents species that occupy both regions, and ∅ represents species extinction. (b) The number of species in region *A* (blue), in region *B* (red), and in either region (gray) over time. (c) Within-region speciation rates in region *A* (blue) and region *B* (red) over time. (d) Extinction rates in region *A* (blue) and region *B* (red) over time. (e) Dispersal rate from region *B* into *A* (blue) and from region *A* into *B* (red) over time. The blue and red start in panels (c)-(e) indicate the start times when regions A and B (respectively) contain 1+ species, corresponding to non-zero rates (in linear scale) for each process. Species richness in regions *A* and *B* both reach local equilibria as time progresses, with increasingly stable biogeographic rates.

To associate local species counts with the absolute rate for each biogeographic process, the DDGeoSSE model multiplies a base rate parameter, *ρ*_*p*_, with a diversitydependent rate factor, 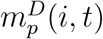, in region *i* for process *p* ∈ {*d, e, w, b*}, corresponding to dispersal, extinction, within-region speciation, and between-region speciation. As an example, for the process *p* = *e*, the extinction rate in region *i* at time *t* is:

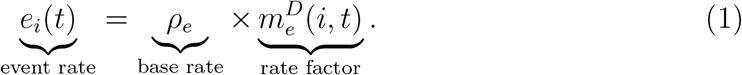

The rate factors for extinction and within-region speciation are defined in terms of single regions, whereas dispersal and between-region speciation are defined in terms of multiple regions. While the base rate parameter, *ρ*_*p*_, has a constant value, the rate factor, 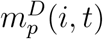, is governed by an estimated diversity-dependent effect parameter, *p*^*D*^, that controls the sign and strength for how species diversity in region(s) *i* at time *t* influences the rate of the process *p* (see Table 1). This flexibility allows biologists to test a wide range of hypotheses concerning diversity-dependent biogeography.

**Table 1:** DDGeoSSE relationships between five biogeographic event types, biogeographic rates, and diversity-dependent effect parameters as regional species richness grows. Negative diversity-dependent effect values decrease the relevant event rates (↓), and positive effect values increase the rates (↑). Zero effect values correspond to the case where species diversity has no influence on event rates (−). Diversitydependent effects that are usually associated with ecological competition are marked with asterices (∗): larger numbers of species in region *i* are predicted to increase (↑) local extinction rates (*e*_*i*_(*t*)) and decrease (↓) local within-region speciation (*w*_*i*_(*t*)) and inbound dispersal (*d*_*ji*_(*t*)) rates.

|  |  | Parameters |  | Effect on rate |  |  |
| --- | --- | --- | --- | --- | --- | --- |
| Process, $p$ | Region(s) | Effect | Rate | $p^D < 0$ | $p^D = 0$ | $p^D > 0$ |
| Extinction | $i$ | $e^D$ | $e_i(t)$ | ↓ | $-$ | $\uparrow^*$ |
| Within-region speciation | $i$ | $w^D$ | $w_i(t)$ | $\downarrow^*$ | $-$ | ↑ |
| Dispersal (outbound) | $i$ | $d^{D,src}$ | $d_{ij}(t)$ | ↓ | $-$ | ↑ |
| Dispersal (inbound) | $i$ | $d^{D,dest}$ | $d_{ji}(t)$ | $\downarrow^*$ | $-$ | ↑ |
| Between-region speciation | $i \in k, j \in \ell$<br>(ranges $k$ and $\ell$) | $b^D$ | $b_\ell^k(t)$ | ↓ | $-$ | ↑ |

After providing the formal probabilistic description of DDGeoSSE, below, we define the equilibrium conditions to attain a desired level of species richness per region, and derive several novel and relevant theoretical results. We use simulation and derived theory to validate our simulator. Next, we use simulations to explore how different DDGeoSSE modeling scenarios produce predictable changes in phylogenetic and biogeographic data patterns. For deep learning-based estimation tasks, we first describe how we simulated training and validation datasets, followed by a detailed description of the pipeline for parameter inference and model selection using phyddle (Landis and Thompson, 2025). Finally, we apply our DDGeoSSE-trained networks to two empirical datasets.

### Logarithmic DDGeoSSE model

Consider a GeoSSE model (Goldberg et al., 2011) defined on a set of *n* discrete regions in the set of regions, *R*. DDGeoSSE allows local species richness to alter the biogeographic rates any particular species experiences, where the rate for each process (within-region speciation, between-region speciation, extinction, and dispersal) changes as a function of the total number of species coexisting at a particular place and time and the diversity-dependent effect parameter. Unless otherwise stated, all following DDGeoSSE analyses use a logarithmic function to model how intensely species diversity influences rates, which we call the *Log-DDG* model variant. Biologically speaking, this means diversity-dependence acts on the orders of magnitude instead of on the raw counts of species richness. In the Supplement, we define an alternative formulation using a linear relationship, called *Standard-DDG*, but we did not use it in our analysis, as we found its behavior to be exceedingly sensitive to its parameter values. We stress that the functional relationship between species richness and the diversity-dependent effect for any process *p* is not restricted to only the logarithmic or linear relationships described here.

Next, we define the effect of five diversity-dependent parameters on the event rates for the four canonical GeoSSE event types: extinction, within-region speciation, dispersal, and between-region speciation (Table 1). The rate for each species to go extinct in region *i* at time *t, e*_*i*_(*t*), is:

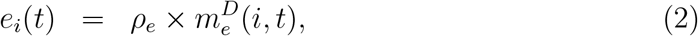

where

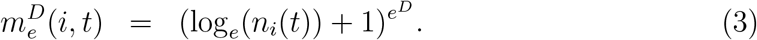

The extinction rate in region *i* are unaffected by diversity-dependent effects only if no other species are currently present (*n*_*i*_(*t*) = 1) or if the effect parameter has no strength (*e*^*D*^ = 0).

The absolute within-region speciation rate in region *i* at time *t* is:

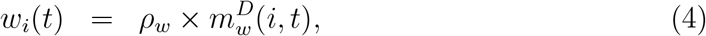

where

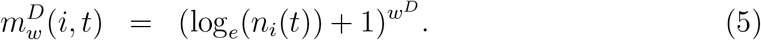

Similar to extinction, these regional speciation rates experience no diversity-dependent effects if no other species are present (*n*_*i*_(*t*) = 1) or if the diversity-dependent effect parameter is zero (*w*^*D*^ = 0).

The dispersal rate from region *i* to *j* at time *t, d*_*ij*_(*t*), is defined as:

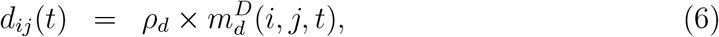

where

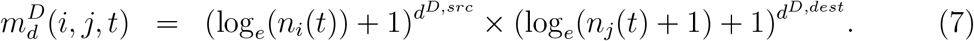

Both *d*^*D,src*^ and *d*^*D,dest*^ are the diversity-dependent effect parameters defined on the source region *i* and the destination region *j*, respectively. The +1 inside the logarithm ensures that the expression is defined when the destination region is empty (i.e., *n*_*j*_(*t*) = 0), whereas the additional +1 outside the logarithm ensures that the diversity-dependent multiplier in the destination region equals one when the region is empty. This also keeps the expression well-defined when *d*^*D,dest*^ ≠ 0. Two simple cases where dispersal is unaffected by diversity levels are when no other species in habit the source region and the destination region has no species, or when the source and destination effect parameters have values of zero.

The between-region speciation rate, 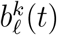, gives rise to two new species with ranges *k* ∈ *S* and ℒ ∈ *S*, respectively, from an ancestral species with range *m* ∈ *S* at time *t*, where *S* is the set of possible species ranges. We define the absolute rate as follows:

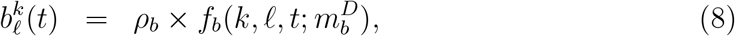

where

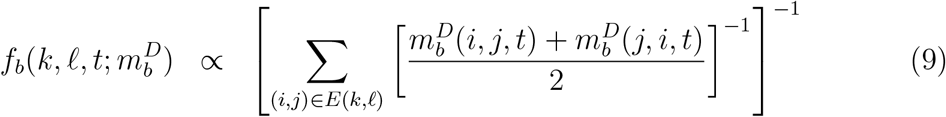

is the range split score function (Landis et al., 2022) that depends on time *t*, species ranges *k* and ℒ, and

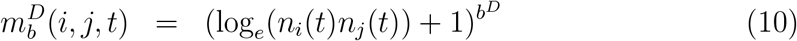

is the diversity-dependent rate factor for species in regions *i* and *j*. The range split score function, *f*_*b*_, computes the relative rates of between-region speciation for different range-splits. A higher range split score means that the particular range split pattern is more likely to occur. *E*(*k, l*) represents the minimal set of edges that must be separated to fully isolate all nodes in range *k* ∈ *S* from all nodes in range *l* ∈ *S*, and the diversity-dependent rate factor 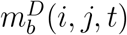 can be thought of as an edge weight that measures the connectivity between region *i* and *j* at time *t*. The edge weight accounts for the numbers of different species currently present in both regions at time *t*. Note that when 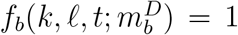, Eq. (9) reduces to the diversity-independent base between-region speciation rate, 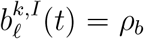 when its effect parameter is zero (*b*^*D*^ = 0) or when the species is alone in all regions it occupies (*n*_*i*_(*t*) = *n*_*j*_(*t*) = 1, ∀ *i, j* ∈ *E*(*k, l*)).

Regional rate variation for each process depends on the species diversity in the relevant region(s) (e.g., *n*_*i*_(*t*)). In the current formulation used for simulation and inference here, we assume that differences in local species diversity is the only factor inducing rate variation within each biogeographic event type. For example, if two regions have identical species diversity, then their extinction rates would be identical as well. However, the framework can be generalized to incorporate non diversity-dependent parameters that are unique to each region into the rate formulation (see discussion in the Supplement).

### Local equilibrium diversity

We give a mathematical definition of local equilibrium diversity for DDGeoSSE in Definition 1, below. Biologically, this equilibrium represents the stationary phase of the process, during which the mean number of local species remains unchanged over time. Importantly, local diversity is tied to each individual region, rather than being the global sum of diversity across all regions. In this section, we derive theoretical results to get this quantity under different formulations of the DDGeoSSE model (see also the Supplement for additional special cases). These results provide a theoretical benchmark for validating our implementation of the DDGeoSSE simulator. Specifically, we can simulate the process until it approaches equilibrium and assess whether simulated local species diversities converge toward the values predicted analytically. This use of equilibrium is specific to simulator validation; our deep-learning inference does not require simulated trees to reach equilibrium, nor do we assume that the empirical datasets are at equilibrium.

In addition, local equilibrium diversity in DDGeoSSE is not a parameter that controls evolutionary rates, but instead it is an emergent property (e.g., expected value) of the data-generating process. This formulation may help reduce bias in statistical inference since we do not target the equilibrium diversity as an estimated parameter (Etienne et al., 2016). We also note that our model does not assume that speciation, extinction, and dispersal are collectively governed by a single, shared diversity-dependence parameter; rather, each process may contribute different amounts to the aggregate level of equilibrium diversity within a region.

**Definition 1**. *Given a diversity-dependent GeoSSE model with range state space S and region state space R, we define the local equilibrium diversity in region i as:*

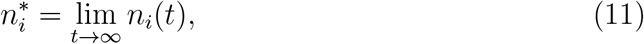

*where n*_*i*_(*t*) *is the solution of the following balance equation:*

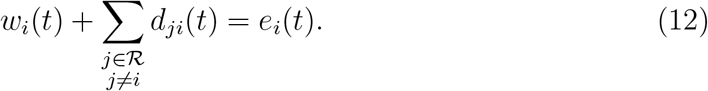

*The left-side of Eq*. (12) *can be thought of as the total incoming rate to region i, I*_*i*_(*t*), *and the right-side of the equation can thought of as the total outgoing rate from region i, O*_*i*_(*t*). *The balance equation can also be viewed as in Figure 2*. *That is*, 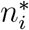 *is such that*

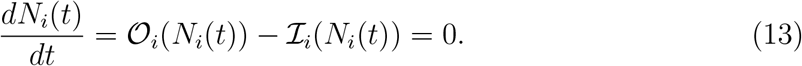

**Figure 2:**
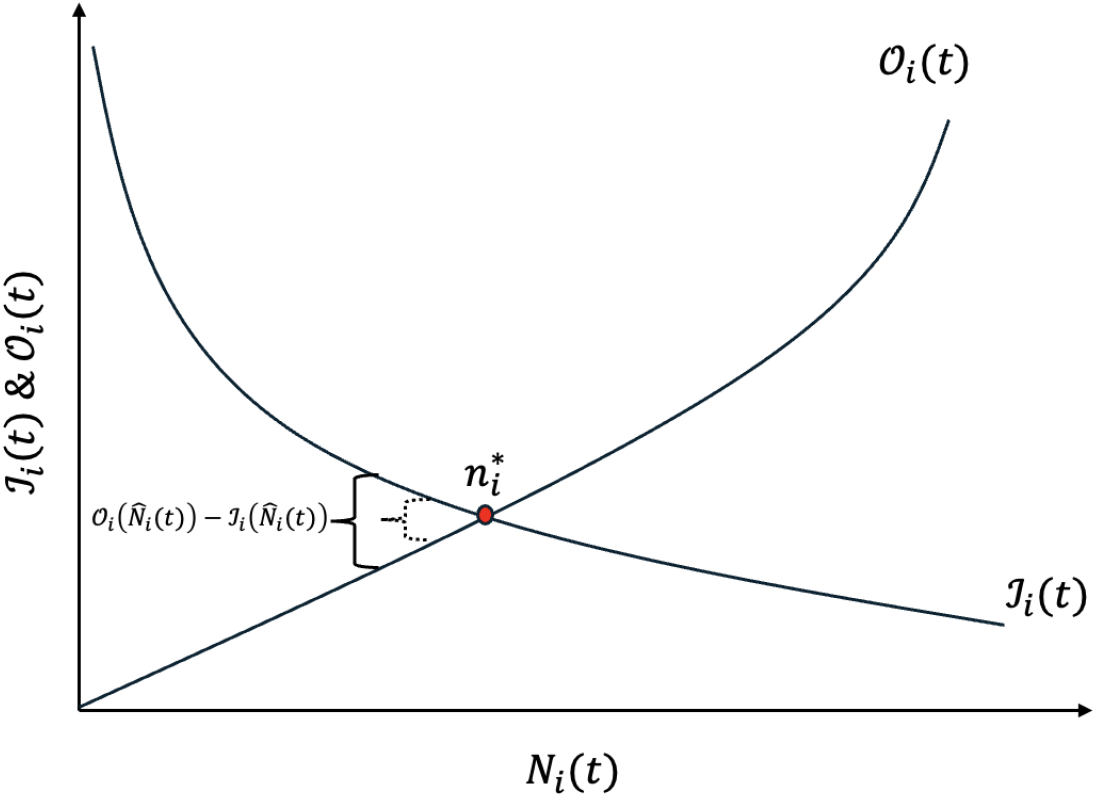
A visualization of the relationship between *O*_*i*_(*t*) and *I*_*i*_(*t*). The equilibrium solution 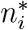 for region *i* is obtained when *O*_*i*_(*t*) = *I*_*i*_(*t*) (as shown by the curly brackets where the difference between the two quantities gets smaller and smaller after each infinitesimal time step Δ*t* until it reaches the intersection point). Note that this figure shares a similar resemblance to the theory of island biogeography from Macarthur and Wilson (1967).

Note that the balance equation is general, meaning that it can be used for any diversity-dependent model that accounts for the same evolutionary processes and local species counts (see also Chevin (2016) for a similar formulation of the balance equation in the BiSSE model (Maddison et al., 2007), and Soewongsono and Landis (2024) for a ClaSSE implementation (Goldberg and Igić, 2012)).

#### Lemma 1.

***(Existence of solution)***

*Given the balance equation described in Eq*. (12) *on an* |*R*|−*region system, where* |*R*| *denotes the number of discrete regions, and let n*_*i*_(*t*) *denote the number of local species diversity at time t* ∈ ℝ_≥0_, *defined on a closed interval* [*t*_*a*_, *t*_*b*_] ⊂ ℝ_≥0_ *where t*_*a*_ *< t*_*b*_. *Assume further that* 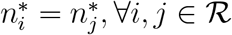. *Then, there exists* 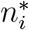 *that solves the balance equation if and only if f*_*i*_(*t*_*a*_)*f*_*i*_(*t*_*b*_) *<* 0 *where:*

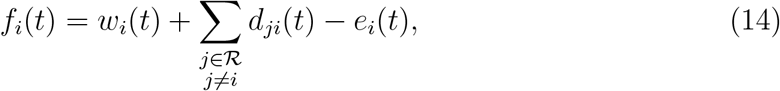

*and* 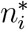 *denotes the local equilibrium diversity in an arbitrary region i*.

The proof of Lemma 1 follows directly from Bolzano’s theorem, since *w*_*i*_(*t*), *e*_*i*_(*t*) and *d*_*ji*_(*t*) are logarithmic functions under Log-DDG, which are continuous on the non-negative real numbers representing *N*_*i*_(*t*). From here on, the theoretical results derived assume only the case of symmetric equilibrium diversity (i.e., 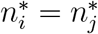).

However, under the current implementation of the Log-DDG model, where rate variation only depends on differences in the numbers of species among regions, one can find model parameters, ***θ*** = {*ρ*_*w*_, *ρ*_*e*_, *ρ*_*d*_, *w*^*D*^, *e*^*D*^, *d*^*D,src*^, *d*^*D,dest*^}, that produce asymmetric (deterministic) equilibrium diversity across all regions (i.e., 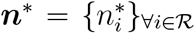, where 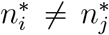). Scenarios inducing asymmetric equilibrium diversity suffer from exchangeability across region labels that can be resolved by adjusting the model to account for intrinsic region-specific characteristics. We provide an example modeling density-dependence (diversity divided by region size) in the Supplement.

Next, we derive the analytical solution for the local equilibrium diversity, 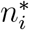, assuming that diversity-dependence acts on all processes relevant to the balance equation described in Definition 1, namely within-region speciation, species dispersal, and species extinction (we refer to this as the *full process*). However, derivations under the full process require some constraints to model parameters to obtain analytical solutions, because the exponent terms (i.e., the diversity-dependent effect parameters) in the balance equation are not always integers, and therefore the equation is not a polynomial. Moreover, considering only polynomial cases, no general closed-form solution exists for polynomials of degree five or higher (Abel–Ruffini theorem). Instead, a numerical solution for 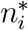 can be found under the unconstrained settings for the full process, such that the following is true:

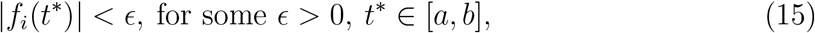

where *f*_*i*_(*t*^∗^) is defined in Lemma 1 and *ϵ* is chosen to be arbitrarily close to zero.

#### Lemma 2.

*Given the balance equation described in Eq*. (12) *and* |*R*| *denotes the number of discrete regions in the system, the local equilibrium diversity in region i under the full process with no diversity-dependent effect on dispersal in the source region*, (*d*^*D,src*^ = 0), *is given by:*

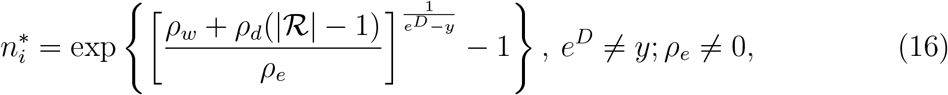

*assuming* 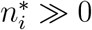 *and w*^*D*^ = *d*^*D,dest*^ = *y under Log-DDG model*.

**Proof:** Proof of Lemma 2 is given in the Supplement.

#### Lemma 3.

*Given the balance equation described in Eq*. (12) *and* |*R*| *denotes the number of discrete regions in the system, the local equilibrium diversity in region i under the full process with time-varying dispersal rate into region i is given by:*

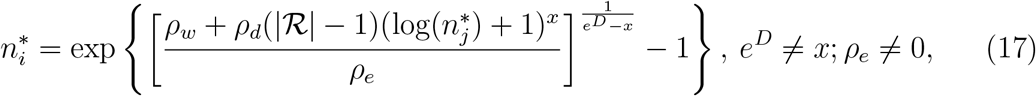

*assuming* 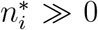 *and w*^*D*^ = *d*^*D,src*^ = *x under the Log-DDG model. Note* 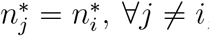, *since the model assumes equal local equilibrium diversity between i and j*.

**Proof:** Proof of Lemma 3 is given in the Supplement

We demonstrate Lemma 1 by using Eq. (15) to numerically solve for 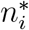 under a fully unconstrained Log-DDG model with parameters *ρ*_*w*_ = 15, *ρ*_*e*_ = 0.03, *ρ*_*d*_ = 0.05, *ρ*_*b*_ = 1, *w*^*D*^ = −0.5, *b*^*D*^ = −1, *d*^*D,dest*^ = −2, *d*^*D,src*^ = −1, *e*^*D*^ = 4, and assuming symmetric local equilibrium diversity across regions 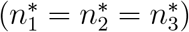. This yields 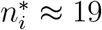 across all regions. Then, we show that simulations under these parameters obtain the desired equilibrium diversity level of 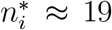 (Fig. 3(a)). Finally, we demonstrate both Lemma 2 and Lemma 3 by analytically finding sets of parameters that correspond to the same 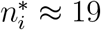. Specifically, we fix all parameters except one and use the corresponding lemma to solve for the remaining parameter such that 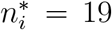. We then validate these results through simulation (Figs. 3(b)-(c), respectively).

**Figure 3:**
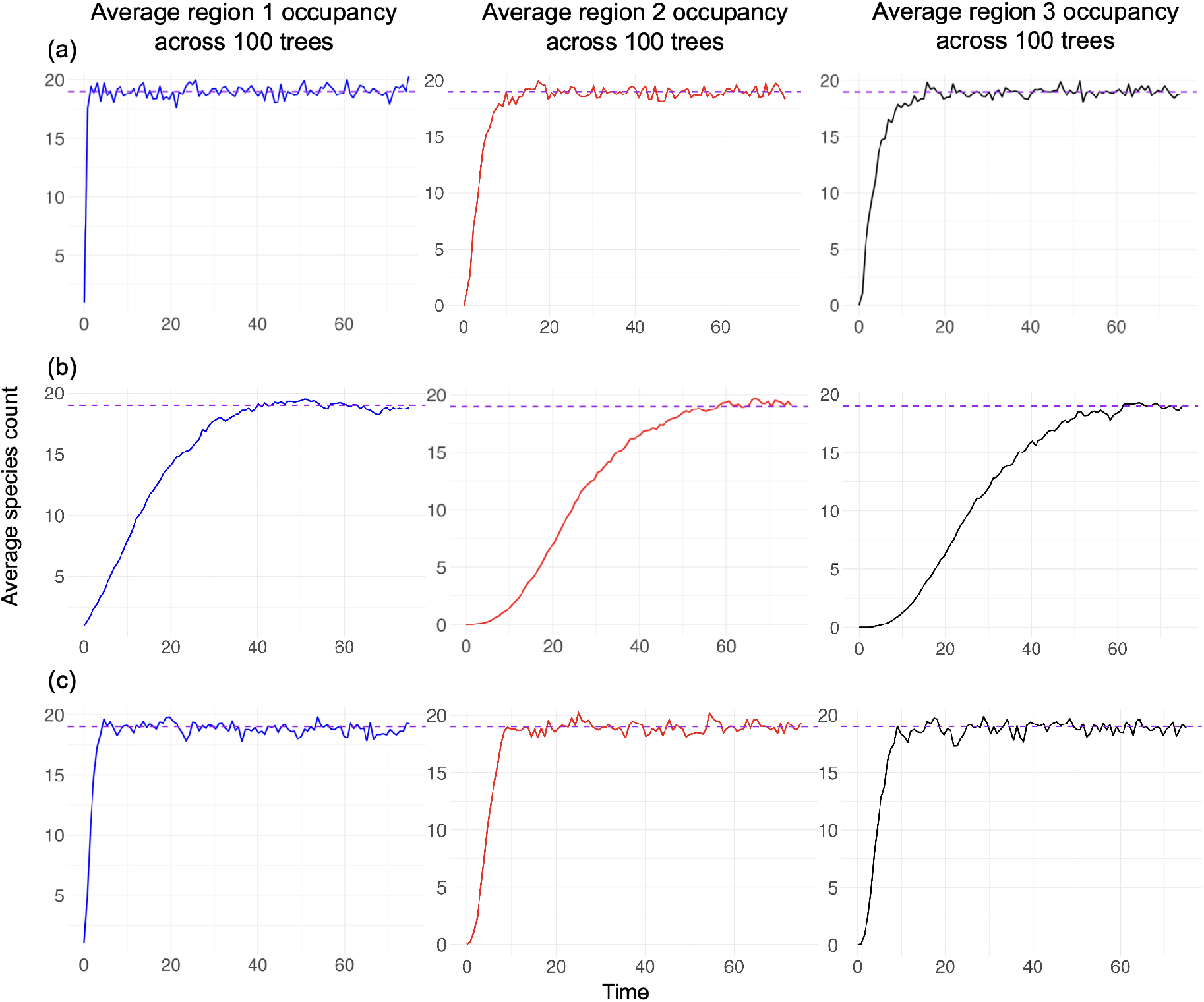
Mean trajectories of *n*_*i*_(*t*), for each of three regions and averaged across 100 replicates. Trajectories were simulated under (a) a fully unconstrained Log-DDG model according to Lemma 1 with parameter values equal to *ρ*_*w*_ = 15, *ρ*_*e*_ = 0.03, *ρ*_*d*_ = 0.05, *ρ*_*b*_ = 1, *w*^*D*^ = −0.5, *b*^*D*^ = −1, *d*^*D,dest*^ = −2, *d*^*D,src*^ = −1, and *e*^*D*^ = 4, (b) a constrained Log-DDG model according to Lemma 2 with parameter values equal to *ρ*_*w*_ = 0.610047, *ρ*_*e*_ = 0.01, *ρ*_*d*_ = 0.01, *ρ*_*b*_ = 1, *w*^*D*^ = *d*^*D,dest*^ = −1, *d*^*D,src*^ = 0, *b*^*D*^ = 0, and *e*^*D*^ = 2, and (c) a constrained Log-DDG model according to Lemma 3 with parameter values equal to *ρ*_*w*_ = 3.730315, *ρ*_*e*_ = 0.03, *ρ*_*d*_ = 0.04, *ρ*_*b*_ = 1, *w*^*D*^ = *d*^*D,dest*^ = *d*^*D,src*^ = −0.5, *b*^*D*^ = 0, and *e*^*D*^ = 3. In all scenarios, each region is expected to have local equilibrium diversity, 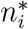, of around 19 extant species. 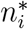 is solved numerically in the fully unconstrained model using Eq. (15). For each tree simulation, we initialize a tree with root species only in region 1.

### Tree statistics for trees simulated under DDGeoSSE model

We explore several tree statistics under DDGeoSSE with diversity-dependence on within-region speciation, extinction, and incoming dispersal (i.e., *w*^*D*^, *e*^*D*^ and *d*^*D,dest*^ ≠ 0) using a 3-region system. We examine variation in mean range size (number of occupied regions) and mean number of extant taxa across the simulations. We also summarize extant tree balance using the size-corrected Colless index (Heard, 1992) and average ladder length (Kendall et al., 2018), and branch lengths using the *γ* statistic (Pybus and Harvey, 2000), as implemented in `treestats` (Janzen and Etienne, 2024).

To simulate data, we construct a 3-dimensional mesh grid of *w*^*D*^, *e*^*D*^ and *d*^*D,dest*^. The values for both *w*^*D*^ and *d*^*D,dest*^ parameters are defined on the interval [−2, 0]. This range of values covers both diversity-independent scenario (*w*^*D*^, *d*^*D,dest*^ = 0) and scenarios where the rates decrease as the local number of species increases (*w*^*D*^, *d*^*D,dest*^ *<* 0) for both events. Meanwhile, the values for *e*^*D*^ are defined on the interval [0, 2], which covers both diversity-independent scenario (*e*^*D*^ = 0) and scenarios where rates increase as the local number of species increases (*e*^*D*^ *>* 0). For each parameter combination in the grid, we simulate 50 extant-only trees of height 100, and compute the mean for each statistic across each tree set. All simulations use event base rates of *ρ*_*w*_ = *ρ*_*d*_ = 0.05 and *ρ*_*e*_ = *ρ*_*b*_ = 0.02. We ignore the case where *w*^*D*^ = *e*^*D*^ = *d*^*D,dest*^ = 0 and exclude invalid trees (trees with only 1 or 2 extant taxa). In total, we simulate 49, 950 extant-only trees across all combinations. To assess the effect weak diversity-dependent effects on tree statistics, we simulate and analyze a second, smaller dataset where *w*^*D*^ and *d*^*D,dest*^ are drawn from [−0.1, 0] interval, and [0, 0.1] for *e*^*D*^. This dataset contains 6, 200 simulated examples.

### Simulated training and validation datasets

We simulate 50, 000 valid training examples under each DDGeoSSE submodel defined in Table 2, while always using five regions and with no diversity-dependent effect on outgoing dispersal (*d*^*D,src*^ = 0) and between-region speciation (*b*^*D*^ = 0). For each replicate, we record the extant-only phylogenetic tree, tip states, and parameter values. Phylogenetic trees are recorded as the Newick strings, while the tip state files record information about species range at tips with one-hot encoding, by marking species as present (1) or absent (0) in each region. The parameter files record the true parameter values from the model for that particular replicate and its model type as described in Table 2. We simulate for *t*_sim_ units of evolutionary time, where *t*_sim_ ~ Unif(70, 100), or until the number of extant taxa reaches *n*_max_, where *n*_max_ ~ Unif(100, 300), whichever occurs first. Trees containing fewer than 10 extant taxa at termination are rejected, along with their model parameters, and resampled. Thus, *n*_max_ represents a randomly sampled stopping threshold rather than a restriction requiring trees to contain between 100 and 300 extant taxa. This procedure ensures that our training examples have varying tree heights and numbers of extant taxa.

**Table 2:**
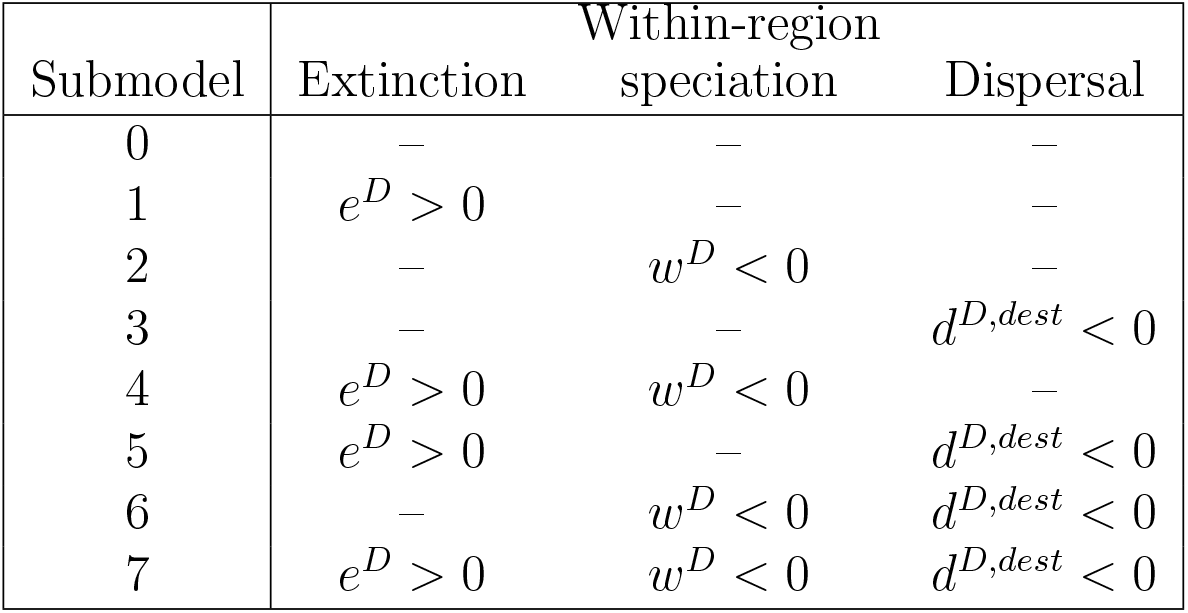
Settings for eight DDGeoSSE submodels without any diversity-dependent effects (0) and with one more more diversity-dependent effects (1-7). Entries marked with a dash (–) correspond to processes with no diversity-dependent effects.

The DDGeoSSE model we analyze assumes each region targets the same level of equilibrium diversity 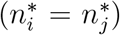, and no other effects alter event rates among regions besides species diversity. For simulation under the null model (submodel 0), we assume equal rates between regions/ranges for each event process. In each submodel, we draw the base rates (*ρ*_*w*_, *ρ*_*e*_, *ρ*_*d*_, *ρ*_*b*_) from the interval of [10^−4^, 1], with 25,000 examples drawn from the uniform distribution, and 25,000 examples drawn from the loguniform distribution. Through experimentation, we found that this mixture improves the equal representation of values that span different orders of magnitude. We draw the values of the diversity-dependent effect parameters (*w*^*D*^, *e*^*D*^, *d*^*D,dest*^) from a uniform distribution on [10^−4^, 2] for *e*^*D*^, and on [−2, −10^−4^] for *w*^*D*^ and *d*^*D,dest*^. Note that we exclude zero from these ranges because a value of zero represents the absence of the corresponding diversity-dependent effect; cases in which each effect is absent are represented separately by setting the corresponding parameter to zero in the relevant submodels (Table 2). These parameter ranges represent scenarios where extinction rates increase, and where within-region speciation and dispersal decrease, with increasing local species diversity. We summarize the details on simulation settings for each submodel in Table 3.

**Table 3:** Table showing simulation conditions and priors distributions used to simulate training datasets for each DDGeoSSE submodel.

| Submodel | Simulation condition | Min #taxa | Max #taxa | Max time | Base rates<br>($\rho_w, \rho_e, \rho_d, \rho_b$) | $w^D$ | $e^D$ | $d^{D,dest}$ | $d^{D,src}, b^D$ |
| --- | --- | --- | --- | --- | --- | --- | --- | --- | --- |
| 0 | time & taxa count | 10 | U(100,300) | U(70,100) | {U($10^{-4}$, 1), LogU($10^{-4}$, 1)} | 0 | 0 | 0 | 0 |
| 1 | | | | | | 0 | U($10^{-4}$, 2) | 0 | 0 |
| 2 | | | | | | U(-2, -$10^{-4}$) | 0 | 0 | 0 |
| 3 | | | | | | 0 | 0 | U(-2, -$10^{-4}$) | 0 |
| 4 | | | | | | U(-2, -$10^{-4}$) | U($10^{-4}$, 2) | 0 | 0 |
| 5 | | | | | | 0 | U($10^{-4}$, 2) | U(-2, -$10^{-4}$) | 0 |
| 6 | | | | | | U(-2, -$10^{-4}$) | 0 | U(-2, -$10^{-4}$) | 0 |
| 7 | | | | | | U(-2, -$10^{-4}$) | U($10^{-4}$, 2) | U(-2, -$10^{-4}$) | 0 |

### Deep learning method for parameter inference and model selection

We use phyddle (Landis and Thompson, 2025), a pipeline-based software for performing phylogenetic modeling tasks on trees using likelihood-free deep learning approaches, for our analyses. The parameter estimation pipeline consists of five main stages:

- **Simulate (S)**. In this stage, we simulate 50,000 training examples under each submodel described in Table 2, where each tree in our training set has between 10 to 300 extant tips. We chose this range because: (1) larger trees provide greater statistical power for inference, and (2) the empirical trees later used in this study fall within this range of simulated tree sizes. Simulated trees were simulated with origin ages between [70, 100] units of time. These bounds are slightly greater than the root age of the *Anolis* tree used in this study, as empirical trees lack an ‘origin’ species (subtending the root node), whereas our simulations start with a single ancestral species. We assume complete sampling of extant taxa when generating these training examples, consistent with the near-complete taxon sampling of the empirical phylogenies analyzed in this study. We use our custom script implemented in Julia (Bezanson et al., 2017) to simulate these datasets (see Supp. Fig. S31 on runtime benchmarking). Users wishing to generate training datasets under alternative DDGeoSSE settings (e.g., different parameter ranges, number of regions, or other model configurations) can modify this simulation script accordingly to produce the required inputs for `phyddle`, and subsequently use the general `phyddle` pipeline for data processing, neural-network training, and validation. Detailed instructions on the required inputs and workflow are provided in the phyddle documentation and tutorials (Landis and Thompson, 2025). We validate our DDGeoSSE simulator by comparing tree statistics from trees simulated under a standard GeoSSE model in Diversitree (FitzJohn, 2012) and our simulator (Supp. Figs. S32 and S33). We further validate the simulator by assessing whether local equilibrium diversity in datasets simulated under DDGeoSSE agrees with the derived theoretical expectations (Fig. 3).
- **Format (F)**. In this stage, we encode our raw simulated data (extant-only trees, tip states, and model parameters) into three types of tensors, namely a phylogenetic data tensor, label tensor, and auxiliary data tensor. A tensor is a multidimensional array used to represent data in deep learning. The phylogenetic data (i.e., topology and branch lengths) and the character matrix data (i.e., tip states) are encoded as phylogenetic data tensor using compact diversity vector (CDV) (Lambert et al., 2023) encoding with one presence-absence state per region associated with each tip (+S) (Thompson et al., 2024). The label tensor encodes the target labels (e.g., model parameters for numerical prediction) to be estimated by the neural network, and auxiliary data tensor contains a panel of summary statistics extracted from the inputted phylogeny and character data matrix for a given dataset (Supp. Table S1). Together, the phylogenetic and auxiliary tensors are used as inputs to train the neural network to predict the corresponding labels.
- **Training (T)**. We used the standard phyddle neural network architecture for our prediction tasks on parameter estimation. For each dataset simulated under each submodel, we reserve 5% of our simulated examples for our test dataset, 5% of examples for our validation dataset, 20% of examples for a calibration dataset, and train our network separately for each model with the 70% remaining examples. We train using stochastic gradient descent with batch sizes of 2,048 examples and using the weight decay option (weight decay = 0.001). Weight decay is a regularization technique that penalizes large neural-network weights during training, thereby constraining model complexity and potentially reducing overfitting, which may improve predictive performance on data not used during training (e.g., empirical data). The weight decay value controls the strength of this regularization, with larger values imposing a stronger penalty on large network weights. In total, we have 16 different trained networks across eight submodels, described in Table 2, for parameter estimation. For this study, we estimate the true parameter values along with their 80% conformalized prediction intervals (CPIs). This means, on average, the true parameter value for a new dataset should fall within CPI 80% of the time, assuming the new data are generated by the same model as used for training. For each epoch, we compute the loss score (*L*_*combined*_), defined as follows:

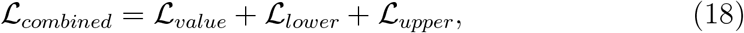

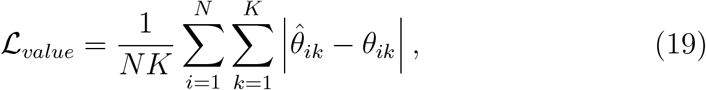

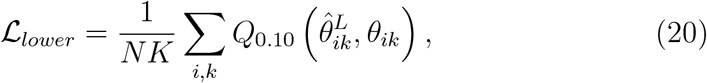

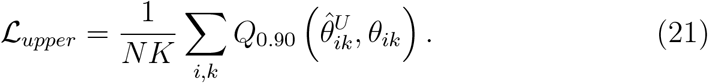

*N* denotes the number of samples in the validation dataset, and *K* denotes the number of parameters in the model. *L*_*value*_ is the mean absolute error (MAE) between the predicted 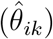 and true (*θ*_*ik*_) parameter values. 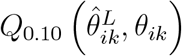 and 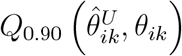 denote the 10th-quantile and 90th-quantile (pinball) losses evaluating the accuracy of the predicted lower 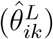 and upper 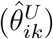 bounds of parameter *k* in dataset *i* relative to the true quantile values. Both quantiles are computed using the following formula:

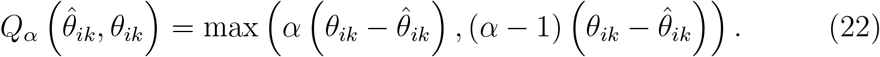

The CPIs are then adjusted through conformalization to ensure the the desired 80% coverage level is obtained (Romano et al., 2019). The loss score for the training data is then minimized to improve network performance, and evaluated for the validation data to measure the general predictive accuracy of the network. The epoch with the lowest validation loss score after 200 epochs of training was used as the trained network. Note that the validation loss is primarily useful for relative comparisons under a fixed model and training setup, as its magnitude depends on parameter scaling and the specific loss function used. Therefore, it should not be used to compare between networks trained using datasets from different models. It is also not to be interpreted as a measure for model adequacy. The latter instead is assessed using model comparison pipeline, which we also describe later in this section.
- **Estimate (E)**. In this stage, we predict the base rate and diversity effect parameters for the test dataset with our trained network from each submodel. We later use the same networks to generate predictions for our empirical datasets.
- **Plot (P)**. Lastly, we visualize the output from phyddle analyses to assess performance of our trained network and the accuracy of our predictions. In this step, phyddle generates PCA plots and histograms of variables for the training and empirical datasets, which used to inspect the plots to identify empirical examples that are outliers (e.g., under model misspecification; Supp. Figs. S18-S24).

Next, we predict parameter values for our empirical datasets using the trained network from each submodel described in Table 2 by running the **F, E**, and **P** steps. To evaluate trained network performance, we plot the point estimates and the 80% CPIs plots (Romano et al., 2019) for the test dataset. We use the slope and intercept of the true parameter value relative to its prediction, and the attained CPI coverage levels, to summarize network performance.

We also used phyddle to predict the best-fitting model among the eight DDGeoSSE submodels (Table 2). We first combined the 50,000 datasets simulated under each submodel to obtain a dataset of size 400,000. Through experiments, we found that using one-hot encoding instead of integer encoding for model types improved network performance (results not shown). That is, instead of using one variable with integers from 0 to 7 to represent the 2^3^ distinct model-type combinations, we used three binary variables to encode presence-absence for each diversity-dependent effect. We converted this combined dataset to tensor data by running the **F** step in phyddle, and then separately trained three networks by running the **T** step on the same dataset to predict whether a particular diversity-dependent feature – *w*^*D*^, *e*^*D*^, or *d*^*D,dest*^ – was non-zero (‘on’). We then trained 7 additional sets of networks for each of the three parameters to test the correctness of the first set of networks, for 24 (= 3*×* 8) networks in total. For each network, the loss score for the categorical model type was computed using the cross entropy loss function implemented in PyTorch library using *torch.nn.CrossEntropyLoss*() function. Similar to the networks trained for parameter estimation, a lower loss score also means better average predictive performance. Given the complexity of the model, we found that using this method (i.e., training for a single categorical prediction and using multiple independent networks) gave a more accurate and consistent prediction results. In other words, we used process-level classifiers to assess whether diversity-dependence is present in each rate component. We then associate the prediction with a submodel whose properties are consistent with the process-level predictions (Table 2). For example, we select Submodel 6 if the networks support diversity-dependence for within-region speciation and dispersal, but not extinction. For empirical analyses, our model selection predictions used the mean prediction scores across the eight networks for each parameter. Lastly, by running the **P** step, we obtained a figure illustrating the best fitting model for each empirical data.

Our phyddle configuration files and simulator scripts are provided with the Supplementary Information. The Supplement also displays the phyddle network architecture used for the parameter estimation (Supp. Fig. S34) and model selection analyses (Supp. Fig. S35).

### Empirical data

We used our DDGeoSSE-trained neural networks to analyze Caribbean *Anolis* lizards and cloud forest-dwelling *Viburnum* plants, allowing us to infer the extent to which either clade was governed by diversity-dependent rates of dispersal, speciation, and/or extinction.

*Anolis* is a adaptive radiation of ~380 neotropical lizard species, with more than half of its species inhabiting the Caribbean islands. Convergent evolution has repeatedly produced similar communities of anole ecomorphotypes within and among different islands (Losos et al., 1998), where ecological interactions among anole species are thought to shape regional patterns of species richness (Losos and Schluter, 2000). Later work by Rabosky and Glor (2010), using their modified MuSSE model, found that biogeographical speciation rates in each island slowed over time, attributing the cause to diversity-dependent deceleration. To test whether regional diversity influences *Anolis* biogeography, we reduced and re-analyzed the dataset of Poe et al. (2017) to obtain a dataset of 158 island species with ranges associated with five Caribbean regions: the Lesser Antilles, Puerto Rico, Cuba and the Caymans, Hispaniola, and Jamaica. The crown age of the phylogeny representing this taxon-subset is ≈64.9 Ma. Supplementary Figure S25 displays the phylogeny and biogeography we used.

The second group, *Oreinotinus*, is a clade of ~40 neotropical plant species, nested within the *Viburnum* radiation (Landis et al., 2021). Earlier work by Donoghue et al. (2022) showed that *Oreinotinus* likely dispersed along a series of montane cloud forests, from North America into South America, repeatedly evolving the same set of leaf ecomorphotypes in each region. Their work found almost no evidence for backdispersal into previously colonized regions, suggesting inbound dispersal rates might decrease with regional diversity. To test this hypothesis, we re-analyzed the *Oreinotonus* time tree and biogeographical ranges published by Donoghue et al. (2022), which contained 41 species spanning 12 regions, but modified it so that it was compatible with our neural networks trained for 5-region systems. We removed the outgroup species (*V. dentatum*) from Eastern North America and two Caribbean species (*V. alpinum* and *V. villosum*), leaving 38 species in our tree. We also combined the 10 remaining regions into 5 regions, using adjacency and shared history as criteria for merging: (1) merging Eastern, Central, and Western Mexico, (2) merging Oaxaca, Chiapas, and Central America, (3) keeping Costa Rica and Panama together, (4) merging Western, Central, and Eastern Colombia and Venezuela, (5) and merging Southern Colombia, Northern and Southern Ecuador, Northern and Southern Peru, Bolivia, and Northern Argentina. The original tree is relatively young (≈10.7 Ma) and does not fit with tree ages from the training dataset, so we rescaled the branch lengths of the tree to be 7 times longer before the analysis, and reverted to the original scale after the analysis. Supplementary Figure S26 displays the phylogeny and biogeography we used.

### Hardware used

All phyddle analyses were performed on a Dell Precision 7920 server with 2× Intel CPUs (112 cores total), 2× Nvidia RTX A4000 GPUs, and 128 GB of system RAM.

## Results

### Tree and range statistics under DDGeoSSE

Speciation, extinction, and dispersal each produce distinguishable patterns of summary statistics as diversity-dependence strength increases (Figure 4; Supp. Fig. S4). Increasing the strength of any diversity-dependent effect (*w*^*D*^ *<* 0, *d*^*D,dst*^ *<* 0, *e*^*D*^ *>* 0) reduces the mean number of extant taxa in simulated phylogenetic data, regardless of the process. For speciation (Figure 4(a)), as *w*^*D*^ becomes more negative, the mean number of occupied regions (range sizes) grows, topologies become less balanced at global scales (Colless index) yet more balanced at local scales (ladder index), and divergence times cluster near the root of the tree (*γ <* 0). For extinction (Figure 4(b)), as *e*^*D*^ increases, range sizes shrink, topologies become dramatically imbalanced at global yet more balanced at local scales, and divergence times remain clustered near the tips of the tree (*γ >* 0). Lastly, for dispersal (Figure 4(c)), as the effect of *d*^*D,dest*^ strengthens, ranges shrink, imbalance increases slightly at global scales, but remains consistently low at local scales, and divergence times tend to fall closer towards the tips. Particularly when diversity-dependent effects are strong (*w*^*D*^ = −2, *e*^*D*^ = 2, *d*^*D,dst*^ = −2), no pair of processes tend to generate datasets with similar combined sets of topology, branch length, and range size summary statistics. These results indicate the three processes may each produce unique signatures in real data that could be extracted to inform an inference method.

**Figure 4:**
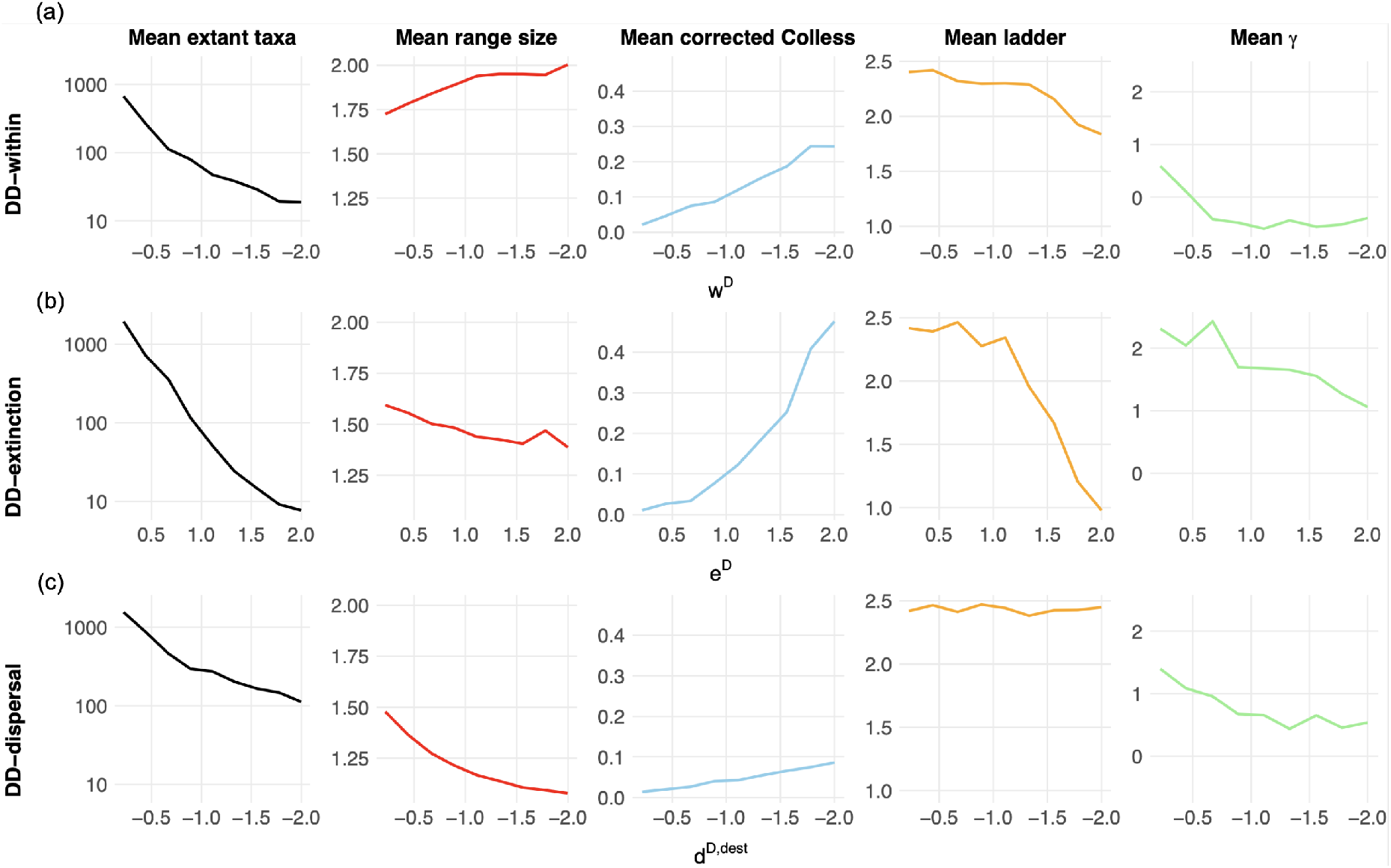
Summary statistics for simulations with diversity-dependent effects on (a) within-region speciation (“DD-within”), (b) extinction (“DD-extinction”), and (c) dispersal (“DD-dispersal”). The *x*-axis represents the values of each diversitydependent effect (top: *w*^*D*^; middle: *e*^*D*^; bottom: *d*^*D,dest*^) and the *y*-axis represent the mean values of the summary statistics. A value of zero is not included on the *x*-axis because these simulations represent submodels in which the corresponding diversitydependent effect is present; zero instead represents the absence of that effect, as represented by the corresponding alternative submodels in Table 2. The summary statistics include the mean number of extant taxa (log-scale), the mean range size (the average number of regions occupied per species), the mean corrected Colless index (tree topology; larger values indicate more imbalanced trees and smaller values indicate more balanced trees at global phylogenetic scales), the mean ladder index (tree topology; larger values indicate longer average ladder structures and smaller values indicate shorter average ladder structures at local phylogenetic scales), and the mean *γ* statistic (distribution of branching times; positive *γ* values indicate branching divergence times clustered near the present and negative *γ* values indicate branching divergence times clustered near the root).

### Parameter estimation and model selection for DDGeoSSE

Simulation results indicate varying degrees of accuracy for parameter estimation under submodel 7, where speciation, extinction, and incoming dispersal are all diversity-dependent (Table 2). All parameters attained the targeted coverage level of 80%, indicating that the prediction intervals are calibrated correctly (Fig. 5). We also observe varying levels of accuracy for parameter estimation among the other models (Table 2; Supp. Figs. S7−S13).

**Figure 5:**
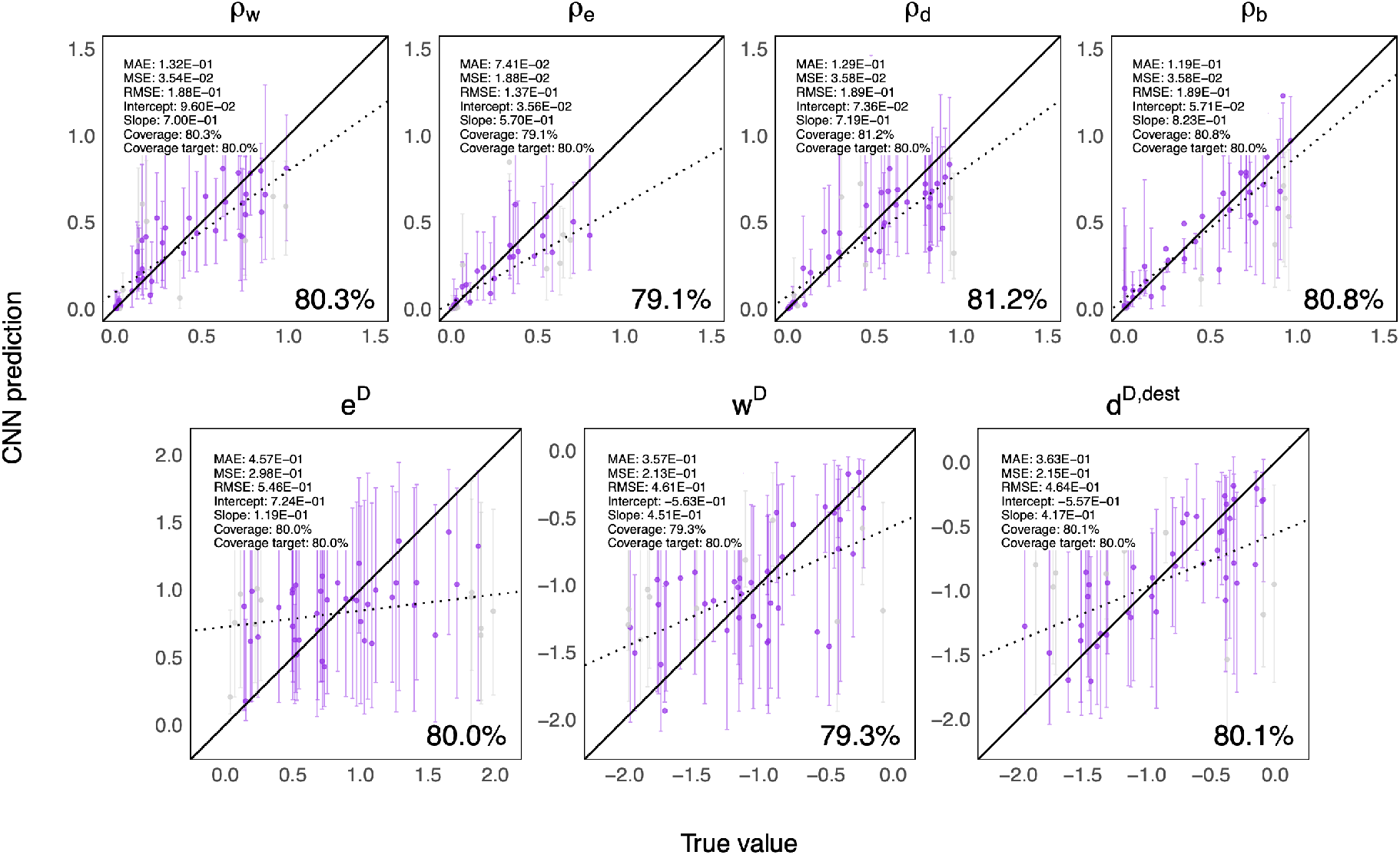
Parameter estimation accuracy against the test dataset simulated under DDGeoSSE submodel 7. The *x*-axis shows the true parameter values, and the *y*-axis shows the estimated values from CNN. All test data were used for the regression. Of these, point estimates (markers) and 80% CPIs (bars) are shown for 50 examples. Estimates are colored if their CPI contains the true value, and left if not. The number shown on the bottom right of each panel indicates the percentage of test cases where the true parameter value falls inside the 80% CPI.

While the estimates for all base rate parameters (*ρ*_*w*_, *ρ*_*e*_, *ρ*_*d*_, *ρ*_*b*_) are precise (tighter intervals), the estimates for the three diversity-dependent effects (*e*^*D*^, *w*^*D*^, and *d*^*D,dest*^) are less certain (wider intervals). In particular, the *e*^*D*^ estimates are the least precise, as extinction-associated parameters are often the most difficult to estimate accurately (Goldberg et al., 2011; Swiston and Landis, 2025). This is consistent throughout all DDGeoSSE submodels with diversity-dependent extinction (Supp. Figs. S8, S11, and S12). We caution readers when interpreting the estimate of this extinction parameter from data.

Next, using three sets of eight independently trained networks, we evaluated the ability of the networks to detect diversity dependence in within-region speciation, extinction, and dispersal. Roughly speaking, positive cases of diversity-dependence for within-region speciation were correctly identified 81% − 87% of the time, and negative cases were identified 73%−81% of the time. Similarly, the networks detected 78% − 86% of positive cases and 76% − 83% of negative cases for extinction, and 83% − 87% of positive cases and 74% − 80% of negative cases for inbound dispersal (Supp. Figs. S15 and S16). In addition, we examined how easily the true existence of one type of diversity-dependence was confused as that of another (Fig. 6 and Supp. Fig. S17). Overall, the networks show stronger support for diversity-dependence in processes where it is truly present (74% − 89%) and lower support when it is absent (9% − 38%), indicating that the process-level signals are generally distinguishable across submodels.

**Figure 6:**
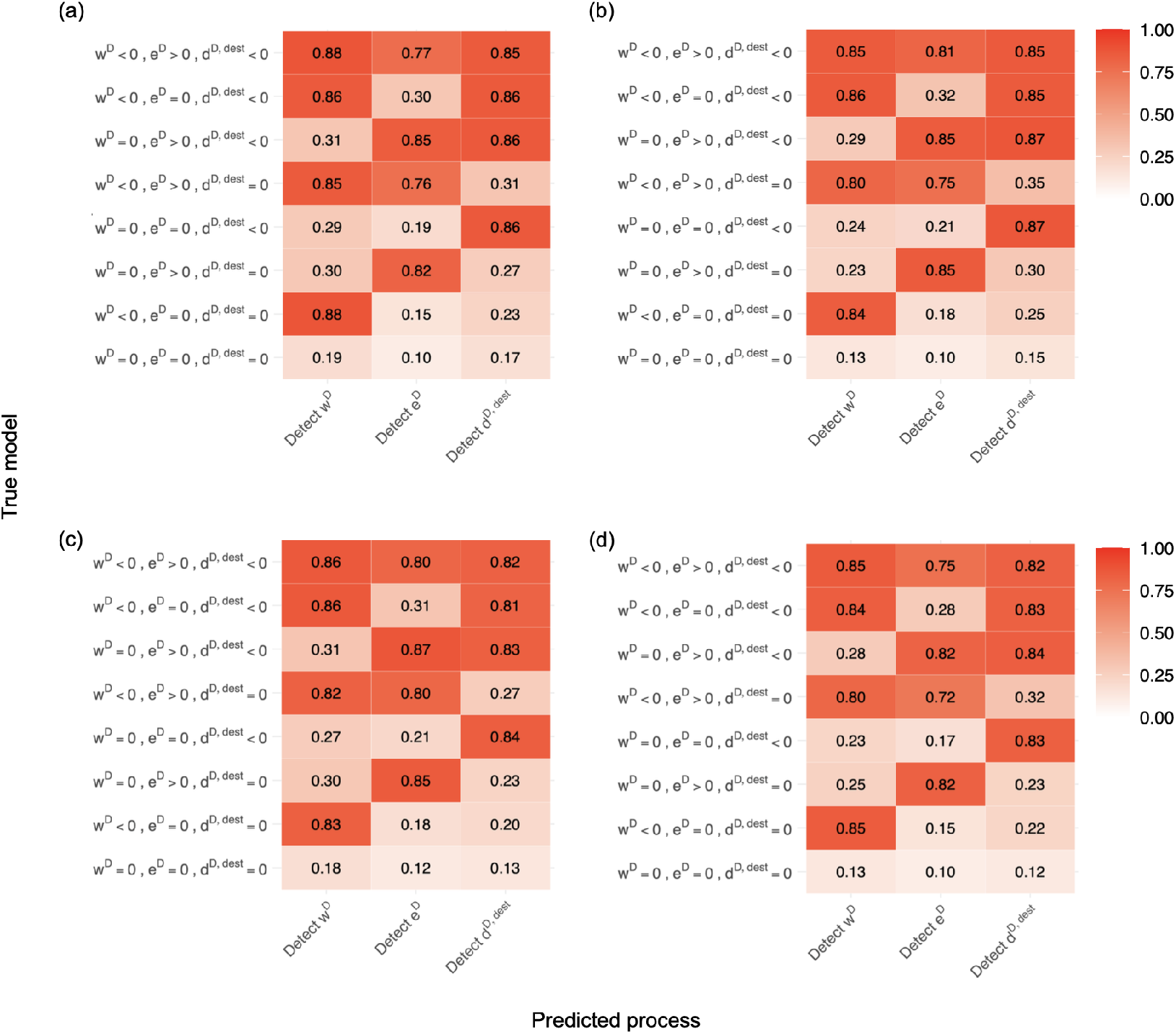
Process-level classification performance across test datasets under different submodels, which are categorized based on their properties of presence or absence of diversity-dependence for within-region speciation (*w*^*D*^), extinction (*e*^*D*^), and incoming dispersal (*d*^*D,dest*^). Rows correspond to the true generating submodel, while columns show the proportion of datasets for which the corresponding process-level classifier detects diversity dependence on within-region speciation, extinction, or incoming dispersal, using a predicted marginal probability threshold of 0.5. Panels (a)-(d) correspond to 4 sets of independent networks (networks 1-4) trained without using non-zero weight decay option in phyddle. Higher values indicate stronger support for diversity dependence in the corresponding process. Overall, the classifiers generally show high support for processes that are truly diversity dependent and lower support for processes that are absent.

### Model selection: Anolis tree

We found that the *Anolis* dataset is best supported by submodel 7, which has diversity-dependent effects on within-region speciation, extinction, and incoming dispersal, with slightly lower support for dispersal (Table 5). Parameter estimates under the best-fitting Submodel 7 were consistent across independently trained networks (Table 4; Supp. Table S2). Using the the number of present-day species in each region, the predicted net diversification rates are generally faster in smaller islands and slower in larger islands (Fig. 7 and Supp. Fig. S14). Overall, these results support the scenario where higher local species diversity reduces within-region speciation and immigration rates (*w*^*D*^, *d*^*D,dest*^ *<* 0), and increases extinction rates (*e*^*D*^ *>* 0).

**Table 4:** DDGeoSSE estimates for Caribbean *Anolis* lizards and Neotropical *Oreinotinus* plants using phyddle. Rows 1-7 show point estimates and 80% CPIs (in brackets) for the best-fitting model. Rows 1-4 show base rates for within-region speciation (*ρ*_*w*_), extinction (*ρ*_*e*_), dispersal (*ρ*_*d*_), and between-region speciation (*ρ*_*b*_). Rows 5-7 show diversity-dependent effect parameters for within-region speciation (*w*^*D*^), extinction (*e*^*D*^), and incoming dispersal (*d*^*D,dest*^). Note that the base rates for *Viburnum* have been converted to correspond to the original phylogeny (branch lengths are measured in unit time). The last column shows the loss score calculated from each model’s corresponding validation dataset.

|  | Estimate | *Anolis* |  | *Viburnum* |  |
| --- | --- | --- | --- | --- | --- |
| Parameter estimation for best-fitting model | Base within-region speciation ($\rho_w$) | 0.2700 | [0.1465, 0.5207] | 1.1382 | [0.4711, 1.7549] |
| | Base extinction ($\rho_e$) | 0.0002 | [0.0001, 0.0012] | 0.0021 | [0.00105, 0.0175] |
| | Base dispersal ($\rho_d$) | 0.0010 | [0.0004, 0.0020] | 0.0070 | [0.00056, 0.0161] |
| | Base between-region speciation ($\rho_b$) | 0.1036 | [0.0016, 0.4191] | 0.0028 | [0.0007, 0.2345] |
| | DD effect on within-region speciation ($w^D$) | -1.64 | [-1.86, -0.88] | -1.52 | [-1.73, -1.09] |
| | DD effect on extinction ($e^D$) | 0.71 | [0.18, 1.66] | — | — |
| | DD effect on incoming dispersal ($d^{D,dest}$) | -1.59 | [-1.93, -0.23] | -1.13 | [-1.98, -0.26] |
| Validation loss score |  | 0.609 |  | 0.478 |  |

**Table 5:** DDGeoSSE estimates for Caribbean *Anolis* lizards and Neotropical *Oreinotinus* plants using phyddle. Networks 1-4 are trained without using non-zero weight decay option, while networks 5-8 are trained using weight decay of 0.001. Model selection support is reported as the marginal probability of the presence of diversity - dependent effects on within-region speciation ℙ (*w*^*D*^ ≠ 0)), extinction ℙ (*e*^*D*^ ≠ 0)), and incoming dispersal ℙ (*d*^*D,dest*^≠ 0)) using 24 trained networks (8 for each evolutionary process). The last three columns show the loss scores from the validation dataset from each network for each process. The last row shows the average probabilities and validation loss scores across all networks.

| Options | Network | *Anolis* |  |  | *Viburnum* |  |  | Validation Loss Score |  |  |
| --- | --- | --- | --- | --- | --- | --- | --- | --- | --- | --- |
| | | $P(w^D \neq 0)$ | $P(e^D \neq 0)$ | $P(d^D \neq 0)$ | $P(w^D \neq 0)$ | $P(e^D \neq 0)$ | $P(d^D \neq 0)$ | $P(w^D \neq 0)$ | $P(e^D \neq 0)$ | $P(d^D \neq 0)$ |
| Without weight_decay | Network 1 | 1.00 | 1.00 | 0.87 | 0.98 | 0.49 | 0.86 | 0.42 | 0.39 | 0.42 |
|  | Network 2 | 1.00 | 1.00 | 0.94 | 0.98 | 0.43 | 0.49 | 0.40 | 0.38 | 0.42 |
|  | Network 3 | 0.98 | 0.98 | 0.80 | 0.93 | 0.25 | 0.97 | 0.43 | 0.38 | 0.40 |
|  | Network 4 | 1.00 | 0.98 | 0.85 | 0.79 | 0.31 | 0.71 | 0.41 | 0.38 | 0.42 |
| With weight_decay = 0.001 | Network 5 | 0.97 | 0.83 | 0.97 | 0.97 | 0.29 | 0.75 | 0.38 | 0.37 | 0.37 |
|  | Network 6 | 0.99 | 0.95 | 0.80 | 0.89 | 0.43 | 0.61 | 0.39 | 0.35 | 0.39 |
|  | Network 7 | 0.97 | 0.94 | 0.97 | 0.89 | 0.51 | 0.55 | 0.39 | 0.36 | 0.37 |
|  | Network 8 | 0.96 | 0.79 | 0.94 | 0.91 | 0.42 | 0.67 | 0.38 | 0.36 | 0.37 |
| Mean value |  | **0.98** | **0.93** | **0.89** | **0.92** | **0.39** | **0.70** | **0.40** | **0.37** | **0.39** |

**Figure 7:**
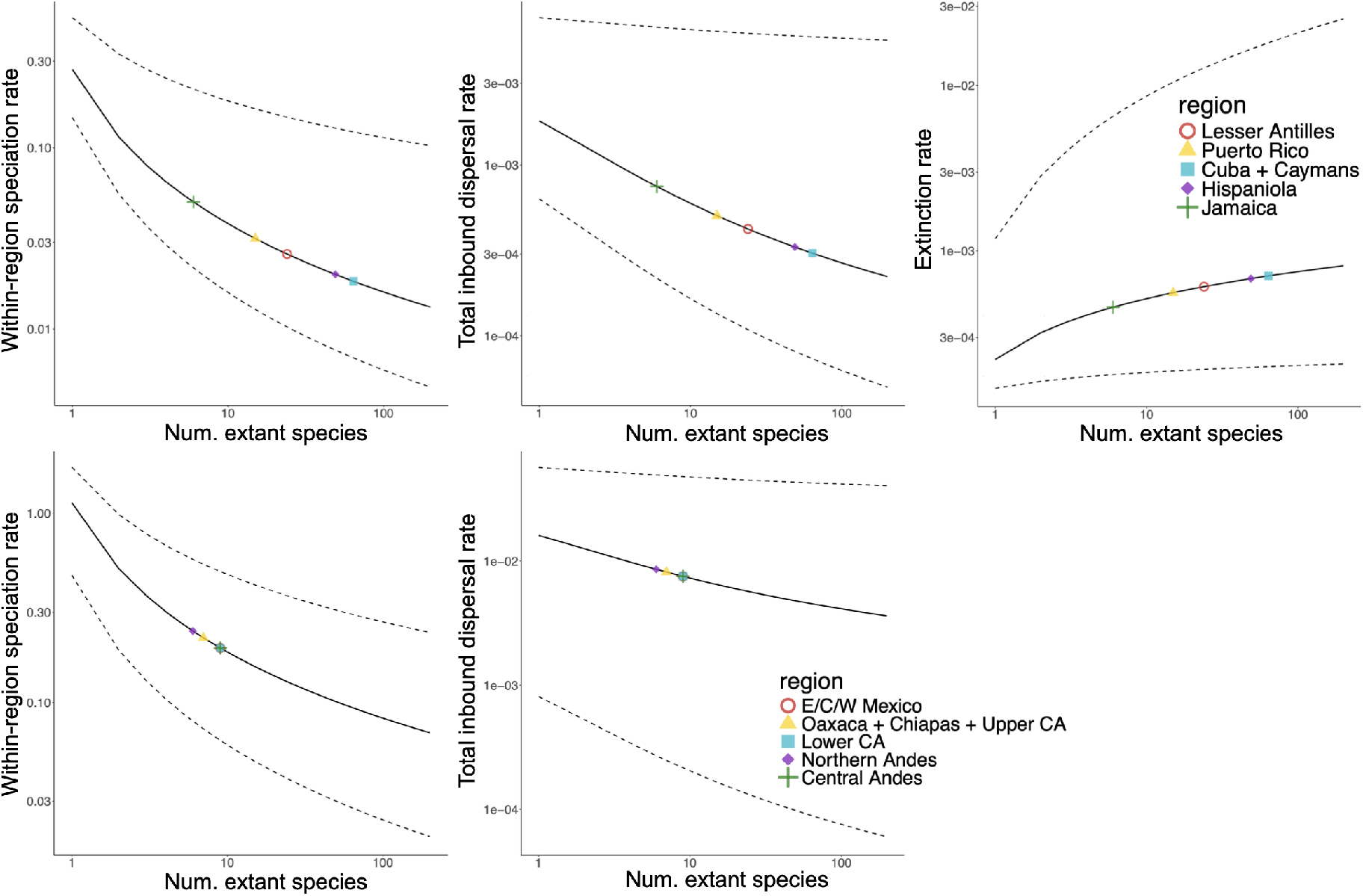
DDGeoSSE rates for *Anolis* (top panel) and *Oreinotinus* (bottom panel) using their best-fitting models, submodel 7 for *Anolis* and submodel 6 for *Oreinotinus*, using the networks trained with weight decay disabled. The *x*-axis shows the number of extant species and the *y*-axis shows the region-specific rates for within-region speciation (left column), total inbound dispersal (middle column), and extinction (right column), using a log-log plot. The total inbound dispersal rate for region *i* is computed using the formula: ∑_*j*_ *d*_*ji*_. The colored symbols represent the regional event rates for the extant species in each region, using the estimates from Table 4. The extinction rate for *Oreinotinus* is not plotted since it is equal to its base rate for all regions under submodel 6.

### Model selection: Viburnum tree

We found that the *Oreinotinus* (*Viburnum*) dataset is supported by submodel 6, which models diversity-dependent effects on within-region speciation and incoming dispersal, but not extinction, with weaker support for diversity-dependent dispersal than for within-region speciation (Table 5). Parameter estimates under the bestfitting submodel 6 were consistent across independently trained networks (Table 4; Supp. Table S2). Owing to the fact that all regions possess 8, 9, or 10 extant species, we found that present-day rates of within-region speciation and incoming dispersal were similar across regions (Fig. 7 and Supp. Fig. S14). Our findings support the hypothesis where higher local species richness reduces both within-region speciation and dispersal rates (*w*^*D*^, *d*^*D,dest*^ *<* 0), but has no effect on extinction.

We note that the ‘treeness’ score for the *Viburnum* dataset was slightly outside the range of the simulated data (0.43 ∉ [0.64, 0.90]) (Supp. Fig. S30). Treeness measures the proportion of internal branch lengths of the tree relative to the overall tree height. That said, the trained network predicts decreased within-region speciation rates as local species diversity increases, which aligns with the *Viburnum* tree having a smaller (more starlike) treeness value.

## Discussion

In this article, we introduce a fully generative, event-based phylogenetic diversification model, called DDGeoSSE. DDGeoSSE is intended to be flexible, and can accommodate a range of evolutionary scenarios in which local species richness influences diversification dynamics through different biogeographic processes. These include scenarios where high regional species diversity can have positive, negative, or neutral feedback on shifts in evolutionary rates for extinction, within-region speciation, between-region speciation, inbound dispersal, and/or outbound dispersal. This framework enables users to simulate species phylogenies and test hypotheses under varying diversity-dependent scenarios.

Our model differs from the few other existing birth-death models of biogeography that for account species richness, such as the the Rabosky-Glor MuSSE model variant (Rabosky and Glor, 2010) and the DAISIE framework (Valente et al., 2015). Unlike the Rabosky-Glor model, diversity-dependent effects in DDGeoSSE are mechanistically modeled as functions of actual species richness rather than modeled deterministically as a linear decline. Unlike DAISIE, DDGeoSSE allows species to move between all region in the system, and does not require the assumption of a mainland region that is unaffected by diversity-dependent effects. Unlike both models, since DDGeoSSE is a generalization of GeoSSE, it models how biogeographic processes operate differently upon species that are widespread (in multiple regions) or are becoming widespread. For example, widespread species in GeoSSE have more opportunities for speciation, produce a variety of possible range inheritance patterns through cladogenesis, and are buffered from the immediate threat of extinction. To the best of our knowledge, DDGeoSSE is the first birth-death model to account for species diversity, historical biogeography, and state-dependent diversification rates with this level of generality of event types (Morlon et al., 2024; Title et al., 2026).

In addition, rather than using a single, shared carrying capacity parameter to model asymptotic species richness (i.e., targeting a pattern; Rabosky and Glor, 2010; Valente et al., 2015), DDGeoSSE instead assigns independent diversity-dependence effect parameters to dispersal, speciation, and extinction. Interactions through these diversity-dependent processes then induce the equilibrium diversity levels for each region. We present techniques to compute the equilibrium diversity level in a region 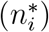 using balance equations, which finds the point at which the rate of species gain and loss in region *i* becomes equal. While the variant of DDGeoSSE used in this study assumes that all regions share the same level of equilibrium diversity 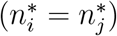, the modeling framework can be adapted to allow different regions to target different equilibrium diversity levels 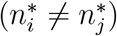. This can be achieved by incorporating regionspecific features, such as area size, as illustrated in Supplementary Figure S3. More generally, our framework can be extended to accommodate interactions between diversity-independent (i.e., abiotic) and diversity-dependent (i.e., biotic) factors as part of the data-generating process (see the discussion in the Supplement).

Simulating under alternative DDGeoSSE scenarios produces distinctive phylogenetic data patterns. Stronger effects of diversity-dependence always reduces species richness, either directly through declining speciation rates (*w*^*D*^ *<* 0) and increasing extinction rates (*e*^*D*^ *>* 0), or indirectly through reducing opportunities for speciation within smaller dispersal-limited range sizes (*d*^*D,dst*^ *<* 0; Fig. 4, first column). Stronger effects decrease mean range size (number of occupied regions per species) for extinction and dispersal, but increase range size through within-region speciation(Fig. 4, second column), as fewer within-region speciation events occur in regions at equilibrium diversity, resulting in fewer young, endemic species. Stronger values of *w*^*D*^ also increase topological imbalance at global, but not local scales. Global tree imbalance (corrected Colless statistic) always increases as diversity-dependent effects strengthens, with extinction being the most responsive, and dispersal the least (Fig. 4, third column). However, local tree imbalance (average ladder length) decreases as effect strength increases for extinction and, to a lesser degree for within-region speciation, but remains high and unchanged for dispersal (Fig. 4, fourth column). Imbalanced trees are also observed in other diversification models, such as age-dependent models where the speciation rate decreases with species age (Hagen et al., 2015; Soewongsono et al., 2022). In contrast with the diversity-dependent model, we do not observe an obvious pattern in tree shape when simulating under very weak to non-diversity-dependent model (Supp. Figs. S4). We conjecture that the contrasting signal in global versus local tree balance for within-region speciation and extinction is the consequence of finescale region-dependent turnover patterns, and the insensitivity tree balance to effects on dispersal is because dispersal events do not directly alter tree topology. Lastly, even though stronger diversity-dependent effects tend to decrease *γ*-statistic values, they remain positive for extinction and dispersal, indicating node ages skew towards the present and produce ‘tippier’ trees, whereas they become negative for speciation, meaning thy skew towards the root and produce starlike trees (Fig. 4, fifth column). In summary, DDGeoSSE produces characteristic patterns for various tree and range statistics, depending on which particular diversity-dependent effect is considered. These distinct patterns could be beneficial for model adequacy testing (see Schwery et al. (2023)), validating simulators that implement diversity-dependent models, and out-of-distribution checks for phylogenetic deep-learning studies (Landis and Thompson, 2025).

Our model uses a phylogenetic deep-learning approach implemented in `phyddle` (Landis and Thompson, 2025) to perform inference, such as for parameter estimation and process-level selection (i.e., presence or absence of the diversity-dependent effect for each process). Using a deep-learning approach bypasses the need to derive a tractable phylogenetic likelihood function, which may not exist for complex models, but is required for likelihood-based inference methods, such as maximum likelihood estimation (MLE) and standard Bayesian inference. The accuracy of different parameter estimates varies among DDGeoSSE submodels but, in general, base rate parameters (*ρ*_*d*_, *ρ*_*e*_, *ρ*_*w*_, *ρ*_*b*_) are estimated more accurately than the diversity-dependent effect parameters (*e*^*D*^, *w*^*D*^, *d*^*D,dest*^), with the diversity-dependent extinction effect parameter (*e*^*D*^) being the least accurate of all. Independently trained model selection networks, on the other hand, all perform quite well and consistent at correctly classifying which datasets did or did not evolve under different diversity-dependence effects, including extinction (Table 5). Thus, our networks appear to be reliable for distinguishing whether diversity-dependence acts on extinction (*e*^*D*^ ≠ 0), but not for estimating the precise value of the parameter. We do not believe that the difficulty estimating *e*^*D*^ is primarily caused by shortcomings with the deep learning approach, as changing network architectures and training strategies did not improve the accuracy (results not shown). We suspect it is simply that extinction-related parameters are intrinsically difficult to estimate across a range of biogeographical (Goldberg et al., 2011; Swiston and Landis, 2025) and diversity-dependent (Rabosky, 2009; Etienne et al., 2016) scenarios. Our sensitivity analysis further demonstrates that model misspecification can reduce inference accuracy. In particular, when networks trained under models that do not account for relative area size were applied to datasets simulated under models incorporating variation in relative area size, CPI coverage deteriorated for several parameters (Table S3). This deterioration is consistent with several of the misspecified datasets falling outside the training distribution, where reliable predictions from the trained networks should not be expected (Figs. S18-S24). We note that this analysis considers only one potential source of model misspecification. Other violations of the assumptions represented in the training simulations may similarly affect inference and remain to be systematically investigated. This consideration is particularly important for empirical applications, where the processes generating observed phylogenetic and biogeographic patterns are unlikely to conform exactly to the models used to generate the training data.

As a proof-of-concept, we applied DDGeoSSE to the empirical phylogenies and biogeographies of *Anolis* and *Viburnum*. For the *Anolis* system, our best-fitting model supports the hypothesis where higher local species diversity reduces withinregion speciation and incoming dispersal rates, and increases extinction rates. Poe et al. (2017) previously noted that *Anolis* species experienced a relatively even number of dispersal events throughout time, which is consistent with our estimates for diversity-dependent dispersal. Rabosky and Glor (2010), using their MuSSE model variant, found evidence of declining rates of diversification in Caribbean anoles (Rabosky and Glor, 2010), which we corroborate with our findings. However, we caution readers that the precise value of our *e*^*D*^ parameter in Table 4 may not be reliable, and is better interpreted merely as *e*^*D*^ *>* 0. Our model predicted that within-region speciation and inbound dispersal rates are highest in Jamaica and lowest in Cuba and the Cayman Islands, with intermediate values in Puerto Rico, the Lesser Antilles, and Hispaniola. Regional extinction rates follow the opposite pattern.

Applying DDGeoSSE to *Oreinotinus* (*Viburnum*), we recovered strong support for a scenario where higher levels of regional species richness limits within-region speciation, and moderate support for lower rates of back-dispersal into already-occupied regions, but we found no clear support for diversity-dependent extinction. We think the support for diversity-dependent dispersal is weaker than that for within-region speciation in part because the tree is relatively small and may not produce strong enough signal. We found species in Central Andes, Lower Central America, and Eastern/Central/Western Mexico, all share the lowest, and identical within-region speciation rate, followed by species in Oaxaca, Chiapas and upper Central America combined with species in Northern Andes having faster rates of within-regions speciation. Dispersal rates followed the same pattern, but with a weaker effect.

These findings align with a scenario where each region is initially seeded by one or two colonizing lineages that diversified early-on until niche saturation prevented the further accumulation of species richness. This is supported by the observation that species richness is higher in the longer-occupied areas and lower in more recently occupied regions (Donoghue et al., 2022), but the difference is too small to be compatible with the unbounded, exponential growth implied by a standard GeoSSE model. We predict that the negative effect of resident species richness on dispersal is consistent with an incumbency effect: i.e., resident species are locally adapted and less likely to be dislodged by newcomers. The long, terminal branches of the *Oreinotinus* phylogeny are also characteristic of scenarios with decreasing extinction and speciation rates. Moreover, recent study documented a likely case of incipient speciation for *Oreinotinus* in the recently occupied Central Andes, indicating that perhaps the clade has not yet equilibrium diversity in the southern parts of its distribution (Maya-Lastra et al., 2024).

New models of lineage diversification have been developed continuously since the introduction of the pure-birth (Yule) model (Yule, 1925). Each model has been proposed to test different hypotheses regarding how lineages diversify through time. Here, we introduced a novel biogeographic modeling framework to investigate how species and their geographic ranges evolve under the influence of diversity-dependent processes. Although the framework is sufficiently flexible to incorporate both diversity-independent and diversity-dependent factors in the data-generating process, the feasibility of inference under such models remains largely unexplored. In particular, it warrants further investigation whether the current inference infrastructure can provide accurate parameter estimates under these more complex scenarios. In the current study, inference is performed using phyddle, which relies primarily on convolutional neural network (CNN) architecture (Voznica et al., 2022; Lambert et al., 2023; Thompson et al., 2023) extract information from phylogenetic patterns.

Future work could investigate whether alternative architectures that more naturally exploit the graphical structure of phylogenetic trees, such as graph neural networks (GNNs) (Xu et al., 2018; Leroy et al., 2025; Du et al., 2026), provide improved performance and robustness in those scenarios. More broadly, the likelihood-free nature of the deep-learning framework offers opportunities to study increasingly complex diversification models for which tractable likelihood functions may be mathematically complex or computationally prohibitive. The trained networks provided in this study may be applied to empirical datasets that are adequately represented by the training distribution and are consistent with the assumptions used to generate the training data. For empirical datasets that fall outside the training distribution (see Figs. S18-S24) or require substantially different model settings, users should generate appropriate training datasets and train new networks following the phyddle workflow (Landis and Thompson, 2025). The likelihood-free nature of this framework also provides a straightforward way to accommodate incomplete taxon sampling. Although we assume complete sampling in the present study, users analyzing empirical clades with substantial known incomplete sampling can first generate trees using our simulator and then subsample extant taxa according to the corresponding sampling fraction before training the neural networks in phyddle, with the sampling fraction supplied as a known parameter. In this way, the trained networks can account for incomplete sampling in the empirical data. However, further work is needed to assess the extent to which incomplete sampling, or misspecification of the sampling fraction, biases parameter estimates, particularly on diversity-dependent effect parameters.

## Conclusion

We have described a novel birth-death process, called DDGeoSSE, that models how spatial rates of speciation, extinction, and dispersal depend on regional patterns of species richness. We derived several key mathematical results from DDGeoSSE and explored the statistical properties of tree shapes generated under our model using simulations. We demonstrated the accuracy of parameter estimation under our model by applying a deep learning approach for phylogenetic studies. By applying our model to two empirical systems, we demonstrated how DDGeoSSE can distinguish diversity-dependent effects on speciation, extinction, and dispersal, allowing explicit tests of process-specific hypotheses underlying biogeographic radiations.

## Data Availability

The dataset and all the relevant code used for this study are publicly available on https://github.com/alberts2/DDGeoSSE.

## Acknowledgements

We are indebted to two anonymous reviewers, Rick Ree, and Bob Thomson, whose insightful comments improved the quality of our study. We are also thankful to members of the Landis lab at Washington University in St. Louis for constructive feedback on an early version of the manuscript. Michael Donoghue has our gratitude for his thoughts regarding our analysis of *Oreinotinus* biogeography.

## Supplementary Materials

### GeoSSE events and state space

We describe the biogeographical model of species diversification, GeoSSE by Goldberg et al. (2011), which associates species’ biogeographical ranges with rates of evolution. The GeoSSE jointly models a branching process and the evolution of species ranges using both cladogenetic and anagenetic events.

Anagenetic events in GeoSSE include species dispersal and local extinction (also known as extirpation) events. A dispersal event leads to range expansion, where a species adds a new occupied region to its range. A local extinction event leads to range contraction, where a species becomes locally extinct in one region. Moreover, a species becomes globally extinct once it goes locally extinct in the last region in its range. Note that, like many other diversification models, GeoSSE also only allows one event occurring within an infinitesimal timestep. As a consequence of this assumption, a widespread species (i.e., species that occupies more than one region) cannot experience a complete extinction through a single event under the model.

Cladogenetic events in GeoSSE include within-region and between-region speciation events. Each within-region speciation event leads to the creation of a new species within any single region of the parental species range. Each between-region speciation causes a widespread parental species and its range to split, such that all regions in the parental range are distributed among the two new daughter species. Supplemental Figure S1 illustrates how we assign rates to these different events, and provides an example of a tree evolving under GeoSSE.

**Figure S1:**
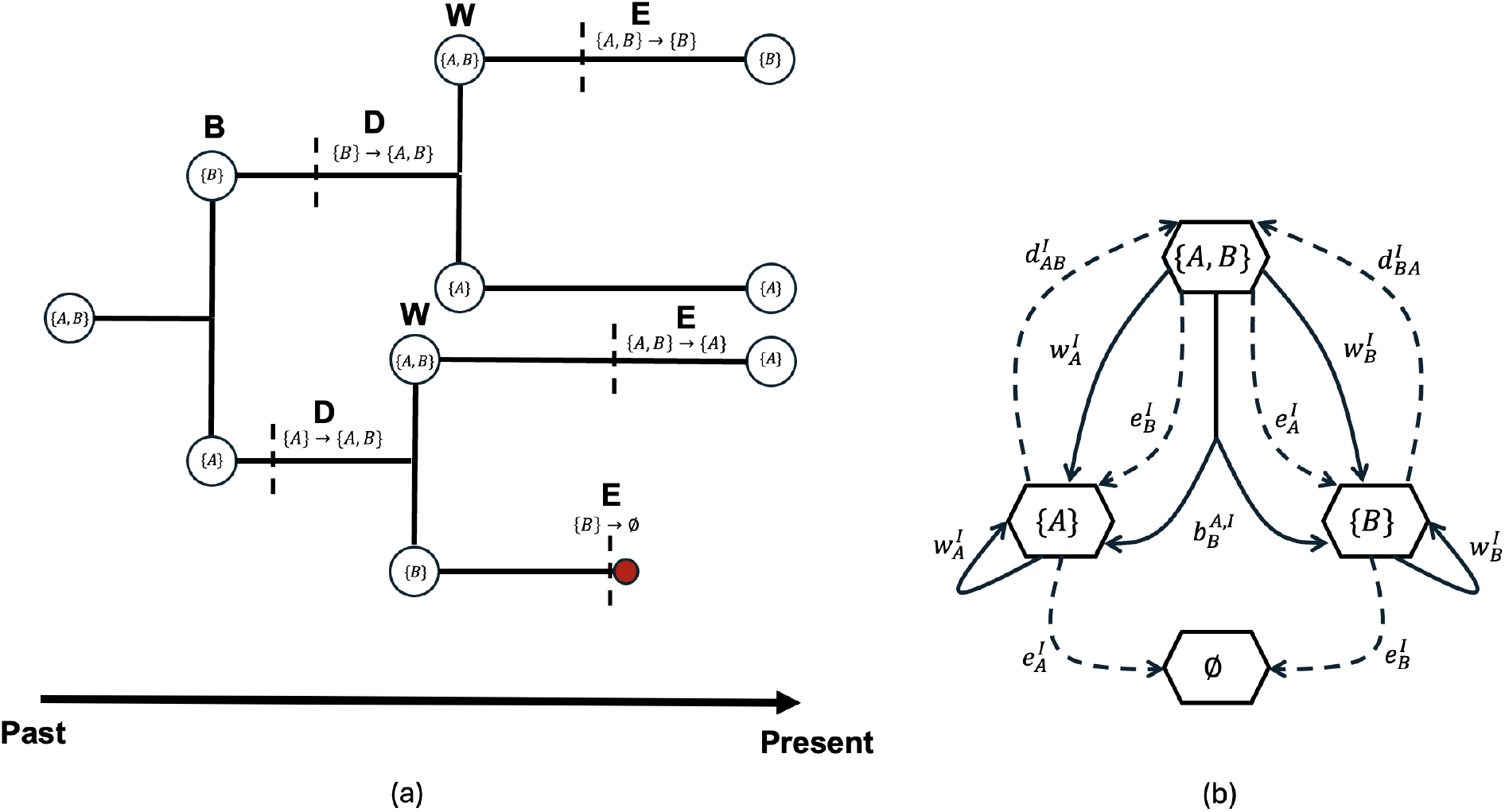
(a) Example phylogenetic tree showing all the history of four different GeoSSE event types in a two-region system (region *A* and *B*) of the model: within-region speciation event ***W*** associated with diversity-independent rates, 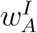 and 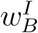, between-region speciation event ***B*** associated with diversity-independent rate, 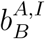, range dispersal event ***D*** associated with diversity-independent rates, 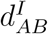 and 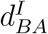, and extinction events (both local and global extinction of a species) ***E*** associated with diversity-independent rates, 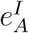 and 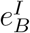. The red node represents a global extinction event of the corresponding species. (b) Transition diagram showing event types with their associated rates in a two-region GeoSSE system (GeoSSE model), following the structure in Goldberg et al. (2011). Solid arrows represent cladogenetic events and dashed arrows represent anagenetic events. Arrows going into the empty set represents a complete species extinction.

### Incorporating diversity-independent rate factors

Here, we describe a way to incorporate diversity-independent rate factors when computing DDGeoSSE event rates. These rate factors may be designed to vary with respect to time, species ranges, or regions. While diversity-dependence can be considered as biotic factor (e.g., species diversity), diversity-independence is more readily associated with abiotic factors, such as regional features (e.g., elevation, area size). Under the version of DDGeoSSE used throughout the main text, which lacks diversity-independent regional factors, asymmetric equilibrium diversity levels are exchangeable with respect to region labels. Incorporating diversity-independent regional factors could be used to mitigate this problem, as different intrinsic regionspecific characteristics would generally lead to different region-specific rates. That said, the absolute extinction rate in region *i* ∈ *R* can be formulated as follows:

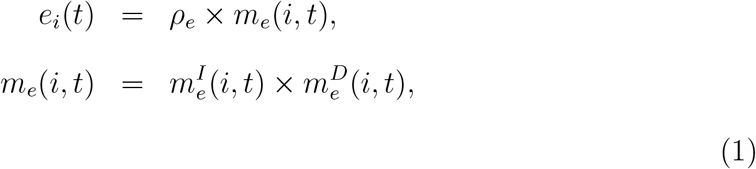

where *m*_*e*_(*i, t*) is the relative rate factor for extinction process in region *i* ∈ *R* at time *t*. It is a product of the diversity-independent rate factor 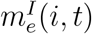 and diversitydependent rate factor 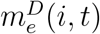, which is described in Eq. (3). Similarly, we can describe the absolute within-region speciation rate in region *i* ∈ *R* at time *t* as follows:

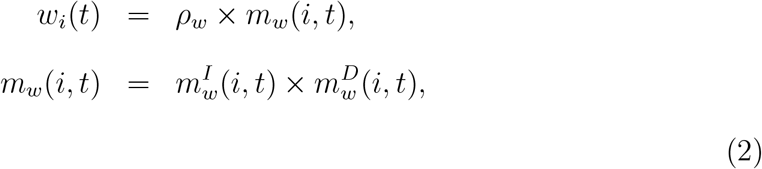

where *m*_*w*_(*i, t*) is the relative rate factor for within-region speciation process in region *i* ∈ *R* at time *t*. It is a product of the diversity-independent rate factor 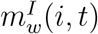 and diversity-dependent rate factor 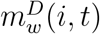, which is described in Eq. (5).

Next, we describe the absolute dispersal rate from region *i* to region *j* at time *t* as follows:

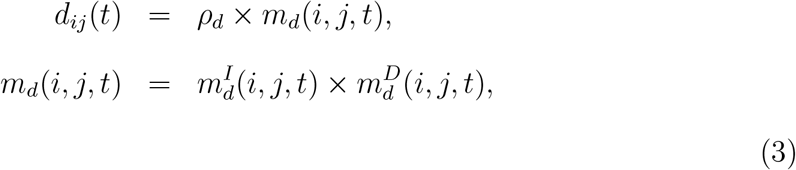

where *m*_*d*_(*i, j, t*) is the relative rate factor for dispersal process from region *i* to region *j* at time *t*. It is a product of the diversity independent rate factor 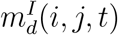 and diversity-dependent rate factor 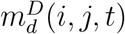, which is described in Eq. (7). Next, we describe the absolute between-region speciation rate, 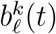, that gives rise to two new species with ranges *k* ∈ *S* and *l* ∈ *S*, respectively, from an ancestral species with range *m* ∈ *S* at time *t* as follows:

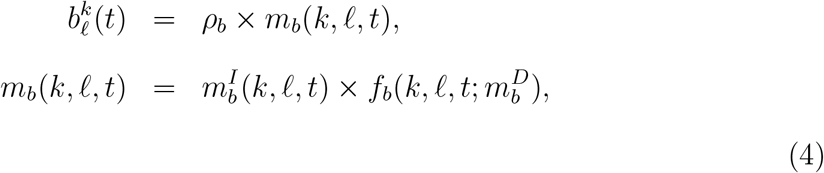

where *m*_*b*_(*k, l, t*) is the relative rate factor for between-region speciation process that gives rise to two new species with range *k* and *l*. It is a product of the diversityindependent rate factor 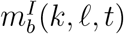 and diversity-dependent rate factor 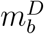 defined via the range split score function, 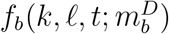, as described in Eq. (9). Note, the definition of 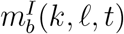 is arbitrary. One could define 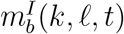 as 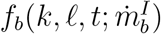 where 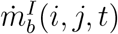 defines how all features shared between regions *i* and *j* influence the rate factors in the range split score function, *f*_*b*_.

### Model variant: Standard DDGeoSSE model

The Standard DDGeoSSE (Standard-DDG) model variant is similar to the Log-DDG model described in the main text, except diversity-dependence of Standard-DDG depends on the actual number of species in a region, rather than the actual natural log number of species. Regardless of which process (*p*) is considered, and unless otherwise stated, the variables in Equations 5 to 8 all share the similar constraints: species counts are positive (*n*_*i*_(*t*) *>* 0, *n*_*j*_(*t*) *>* 0), rates are positive (*p*_*i*_(*t*) *>* 0, *ρ*_*p*_ *>* 0), relative rate factors are positive 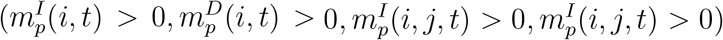, and diversity-dependent effect parameters are realvalued 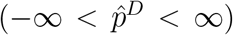. We define the absolute extinction rate for a species in region *i* at time *t* as follows:

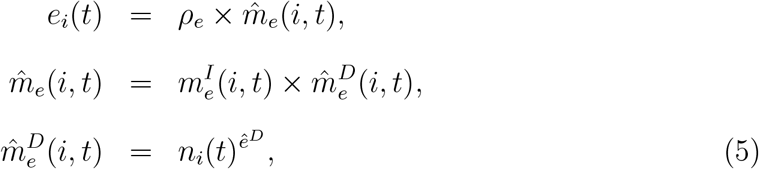

where the diversity-dependent effect parameter *ê*^*D*^ has the same interpretations as *e*^*D*^ in Table 1. Additionally, in the case of *ê*^*D*^ = 1, we have a linear-like behaviour in overall rate increases (resp. decreases) over time as the number of species in region *i* increases (resp. decreases).

Similarly, we define the absolute within-region speciation rate for a species in region *i* at time *t* as follows:

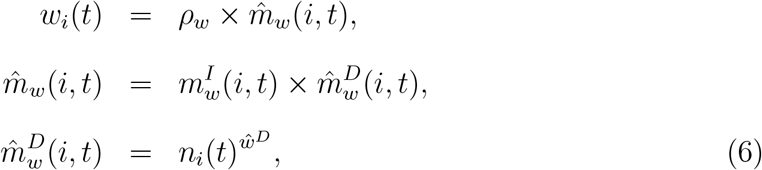

where the diversity-dependent effect parameter *ŵ*^*D*^ has the same interpretations as *w*^*D*^ in Table 1. Additionally, in the case of *ŵ*^*D*^ = 1, we have a linear-like behaviour in overall rate increases (resp. decreases) over time as the number of species in region *i* increases (resp. decreases).

Next, we define the absolute dispersal rate from region *i* to region *j* at time *t*, as follows:

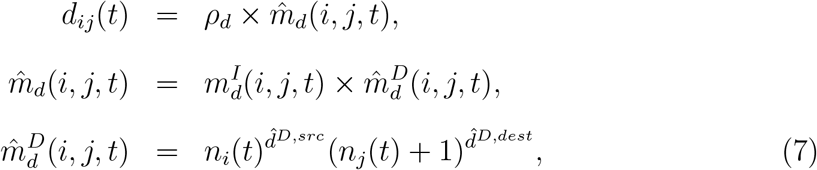

where the diversity-dependent effect parameters 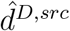 and 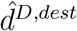 have the same interpretations as *d*^*D,src*^ and *d*^*D,dest*^, respectively, in Table 1. For dispersal, the destination region *j* may contain 0 species: *n*_*j*_(*t*) ≥ 0. Additionally, in the case of 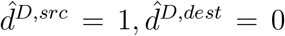 or 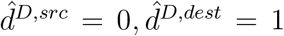, we have a linear-like behavior in overall rate increases (resp. decreases) over time as the numbers of species in regions *i* and *j* (both) increase (resp. decrease).

Finally, we define the absolute between-region speciation rate that gives rise to two new species with ranges *k* ∈ *S* and *l* ∈ *S*, respectively, from an ancestral species with range *m* ∈ *S* at time *t*. We define the overall rate as follows:

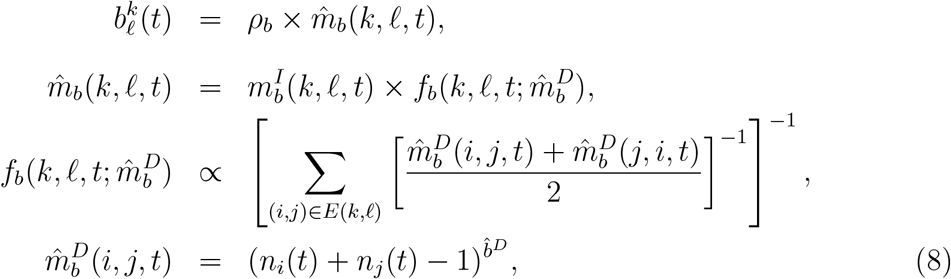

where the diversity-dependent effect parameter 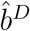 has the same interpretations as *b*^*D*^ in Table 1. Additionally, in the case of 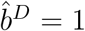, we have a linear-like behavior in absolute rate increases (resp. decreases) over time as the number of species in region *i* increases (resp. decreases).

### Local equilibrium diversity under DDG with only diversity-dependent within-region speciation and extinction (no dispersal)

We consider a scenario where only diversity-dependent within-region speciation and extinction, but with no dispersal whatsoever, drive the diversification process. We derive the solution to the local equilibrium diversity described in the main text under this particular scenario.

#### Lemma S1.

*Given the balance equation described in Eq*. (12) *in the main text, the local equilibrium diversity in region i under scenario with only within-region process and constant extinction rate is given by:*

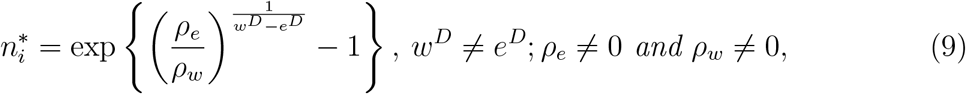

*under the Log-DDG model, and:*

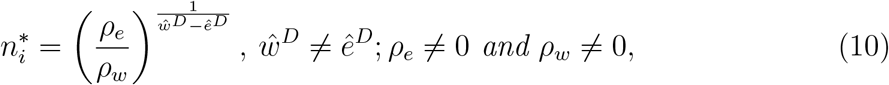

*under the Standard-DDG model. Moreover, if we have constant extinction rate, then both Eqs*. (9) *and* (10) *become:*

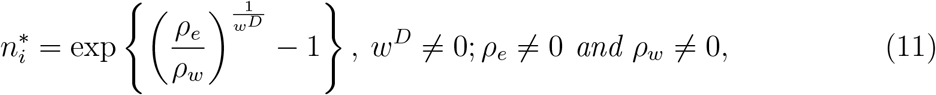

*and*

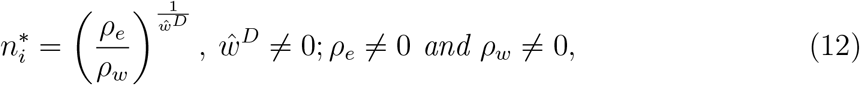

*respectively*.

*Proof*. We show the proof for the lemma under the Log-DDG model as the proof under the Standard-DDG model follows in a similar manner.

Following Eq. (12) in Definition 1 and by substituting Eqs. (3) and (5) we have:

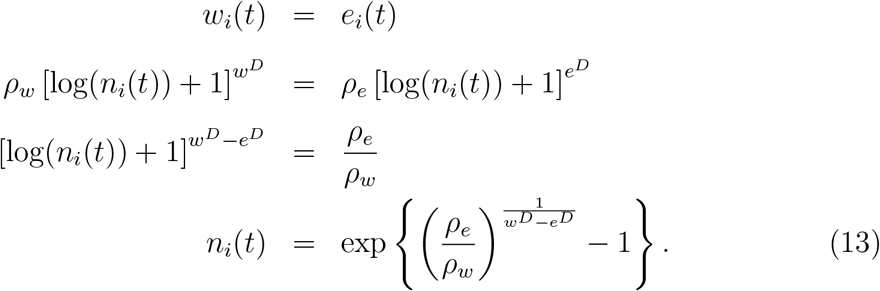

So,

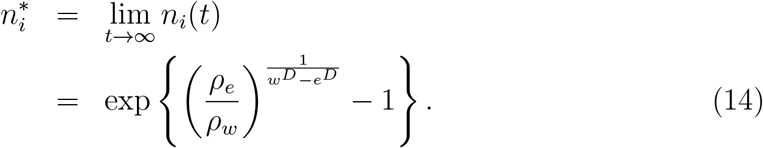

Then, we can set *e*^*D*^ = 0 in Eq. (14) to get a constant extinction rate. Clearly, *ρ*_*w*_ cannot be equal to 0. Moreover, given *ρ*_*w*_ ≠ 0, *ρ*_*e*_ also cannot be equal to 0 since we assume that extinction process exists.

## Local equilibrium diversity under DDG with diversity-dependent within-region speciation

We derive the solution to the local equilibrium diversity described in the main text under Log-DDG model with diversity-dependent within-region speciation and time-constant extinction and dispersal.

### Lemma S2.

*Given the balance equation described in Eq*. (12) *in the main text, the local equilibrium diversity in region i under the DDGeoSSE model with diversity-dependent within-region speciation is given by:*

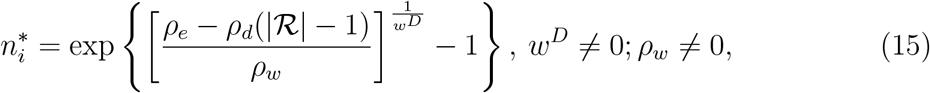

*assuming* 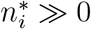 *under the Log-DDG model*.

*Proof*. Following Eq. (12) in Definition 1 and by substituting Eqs. (3) and (5) we have:

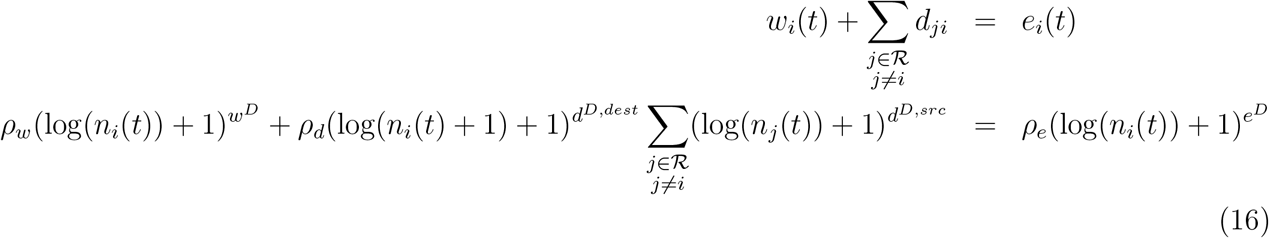

We solve the above equation at equilibrium diversity across locations (*t* → ∞). That is:

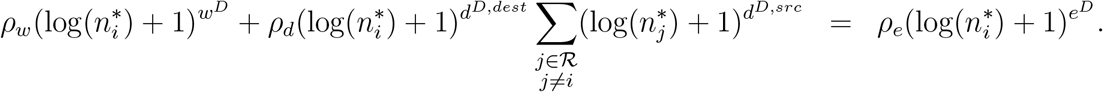

Then by assumption we have *d*^*D,src*^ = *d*^*D,dest*^ = *e*^*D*^ = 0. That is,

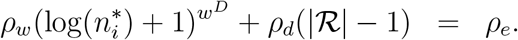

Thus,

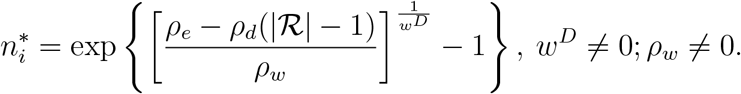

## Local equilibrium diversity under DDG with diversity-dependent extinction

We derive the solution to the local equilibrium diversity described in the main text under Log-DDG model with diversity-dependent extinction and time-constant within-region speciation and dispersal.

### Lemma S3.

*Given the balance equation described in Eq*. (12) *in the main text, the local equilibrium diversity in region i under DDGeoSSE model with diversitydependent extinction is given by:*

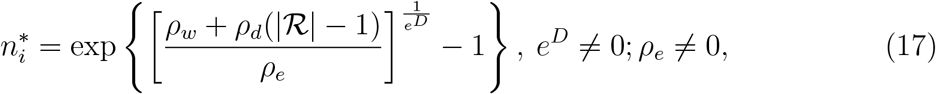

*assuming* 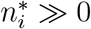 *under the Log-DDG model*.

*Proof*. Following the same steps as the proof for Lemma S2 and by substituting *d*^*D,src*^ = *d*^*D,dest*^ = *w*^*D*^ = 0 instead, it follows that:

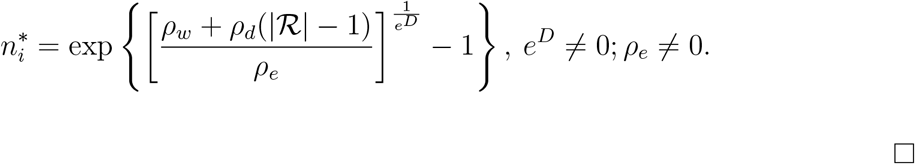

We demonstrate Lemma S3 using simulations, as shown in Figure S2

**Figure S2:**
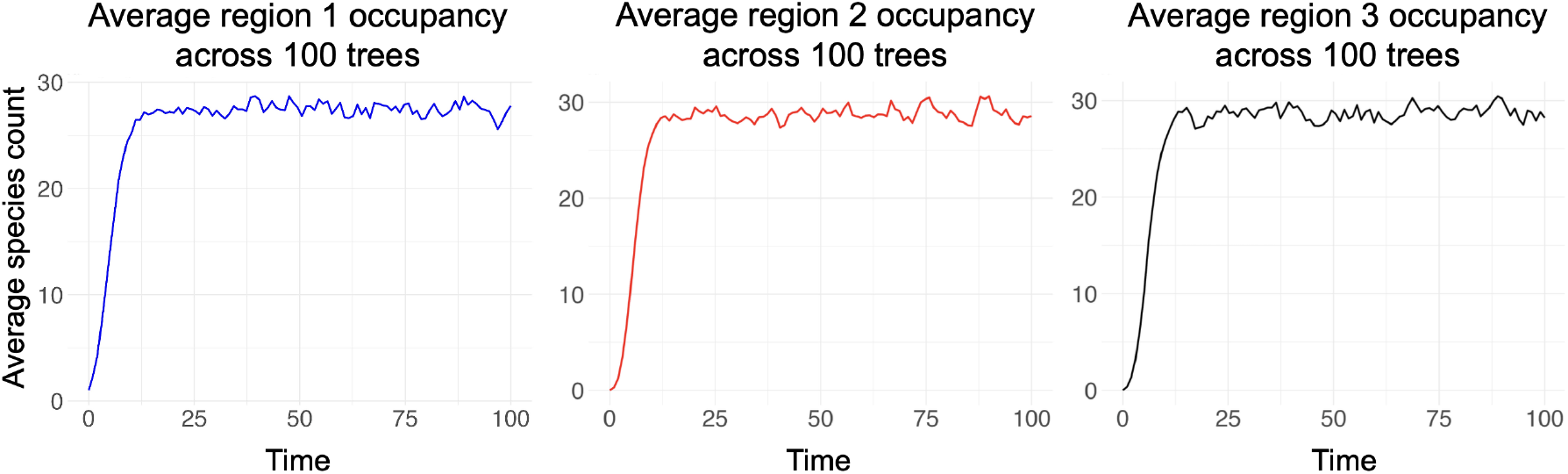
The trajectories of *n*_*i*_(*t*) in all three regions where 100 trees are simulated the Log-DDG model with 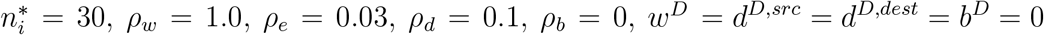, and *e*^*D*^ was chosen according to Lemma S3. For each tree simulation, we initialize a tree with root species in range {*A*}.

*Proof of Lemma 2*

*Proof*. Following Eq. (12) in Definition 1 and by substituting Eqs. (3) and (5) we have:

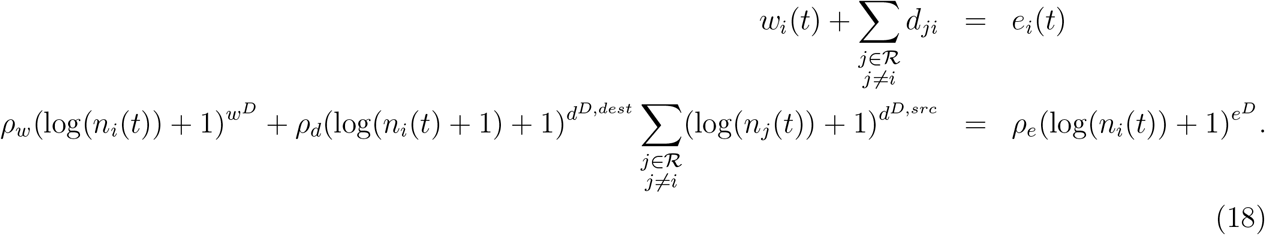

We solve the above equation at equilibrium diversity across locations (*t* → ∞). That is:

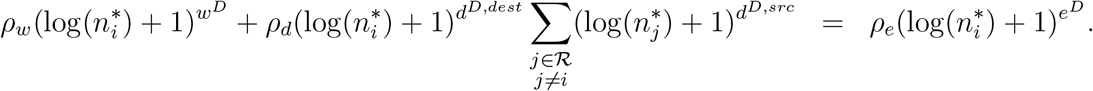

Assuming 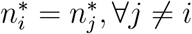, and *d*^*D,src*^ = 0, we have:

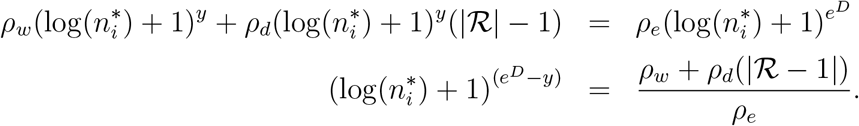

Thus,

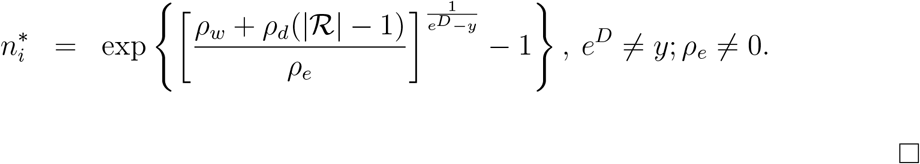

*Proof of Lemma 3*

*Proof*. Following Eq. (12) in Definition 1 and by substituting Eqs. (3) and (5) we have:

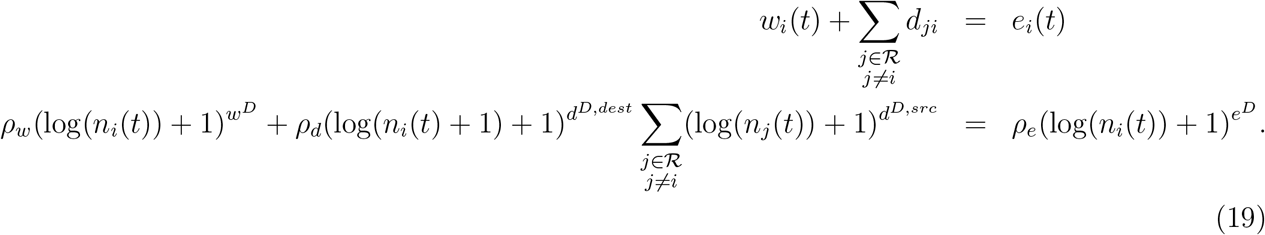

We solve the above equation at equilibrium diversity across locations (*t* → ∞). That is:

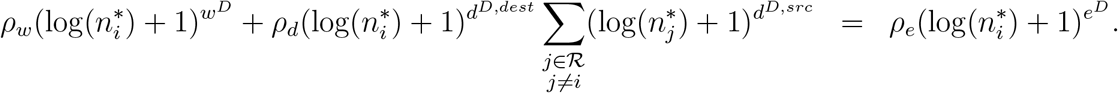

Assuming 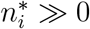 and *w*^*D*^ = *d*^*D,dest*^ = *d*^*D,src*^ = *x*, we have:

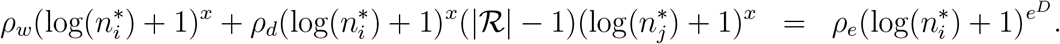

Note here from model assumptions, 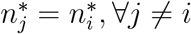. Then,

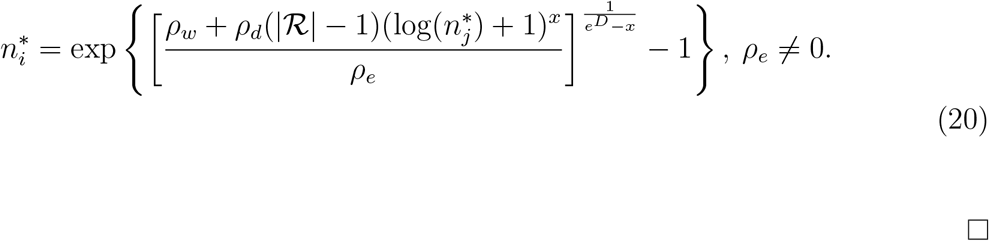

## Local equilibrium diversity under Standard-DDG with full process and without diversitydependent on outbound dispersal

### Lemma S4.

*Given the balance equation described in Eq*. (12) *in the main text and* |*R*| *denotes the number of discrete regions in the system, the local equilibrium diversity in region i under the full process with no diversity-dependent effect on dispersal in the source region*, (*d*^*D,src*^ = 0), *is given by:*

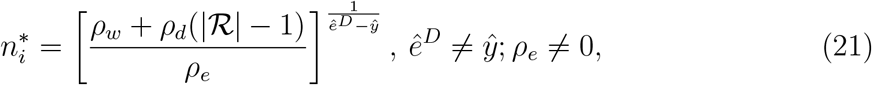

*assuming* 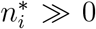 *and* 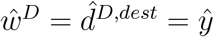 *under the Standard-DDG model*.

*Proof*. The proof under the Standard-DDG model follows in a similar manner to the proof under the Log-DDG model in Lemma 2.

*Local equilibrium diversity under Standard-DDG with full process and time-varying outbound dispersal*

### Lemma S5.

*Given the balance equation described in Eq*. (12) *in the main text and* |*R*| *denotes the number of discrete regions in the system, the local equilibrium diversity in region i under the full process with time-varying dispersal rate into region i is given by:*

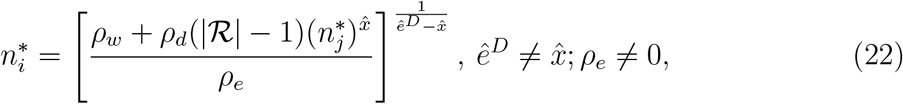

*assuming* 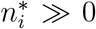 *and* 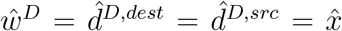 *under the Standard-DDG model. Note* 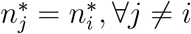 *since the model assumes equal local equilibrium diversity between i and j*.

*Proof*. The proof under the Standard-DDG model follows in a similar manner to the proof under the Log-DDG model in Lemma 3.

## Model variant: Log-DDG model with relative area size

Under the current implementation of the *Log-DDG* model, where rate variation only depends on differences in the numbers of species among regions, one can find model parameters, ***θ*** = {*ρ*_*w*_, *ρ*_*e*_, *ρ*_*d*_, *w*^*D*^, *e*^*D*^, *d*^*D,src*^, *d*^*D,dest*^}, that would produce asymmetric (deterministic) equilibrium diversity across all regions. To find those parameter values, we solve the system of balance equations simultaneously. Note that, under the assumption of equal equilibrium diversity across regions, only one balance equation needs to be solved. Specifically, we find ***θ*** such that:

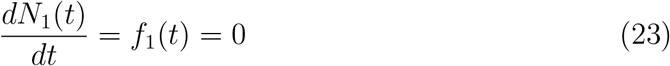

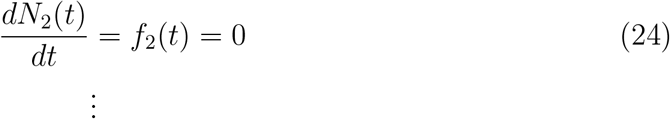

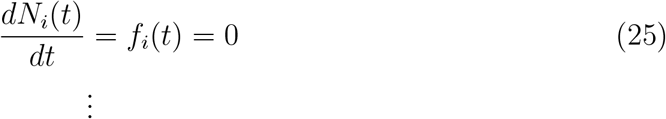

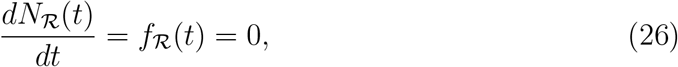

where *f*_*i*_(*t*) is the balance equation for region *i*, as defined in Eq. (14), evaluated at equilibrium 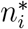, and |*R*| is the number of regions. However, the system of nonlinear equations above is underdetermined if the number of regions is fewer than the number of model parameters, i.e., |*R*| *<* |***θ***|. Thus, no unique solution for ***θ*** exists; multiple parameter combinations may produce the same set of local equilibrium diversities. Note that this does not imply model non-identifiability since phylogenetic patterns and tip character data may resolve the ambiguity.

Additionally, for a given set of model parameters ***θ***, one can solve for ***n***^∗^ by simultaneously solving the system of balance equations. The solution ***n***^∗^ may not be unique because the balance equations are nonlinear (logarithmic function). Since the parameters are not region-specific, asymmetric deterministic equilibria suffer from exchangeability with respect to region labels. That is, if an asymmetric equilibrium exists, then equivalent equilibria obtained by permuting region labels also exist. Equivalently, this corresponds to swapping the order of equations in the system of balance equations. For example, if 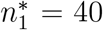 and 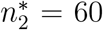 60 is a solution, then 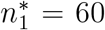 and 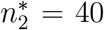 is also a solution. Moreover, if asymmetric equilibrium configurations are realized in the stochastic process (i.e., tree simulations), they would arise through stochastic variation rather than from region-specific intrinsic factors. In this case, the exchangeability of region labels implies that different replicates may approach different label-permuted versions of the same equilibrium. Consequently, averaging across replicates may obscure these asymmetric equilibria and produce an approximately symmetric mean equilibrium diversity pattern across regions.

Under a more general formulation of our model that accounts for additional sources of rate asymmetry (e.g., abiotic or other region-specific factors), the exchangeability of region labels associated with asymmetric equilibrium diversity can be alleviated. In this setting, a region exhibiting higher equilibrium diversity than other regions is no longer a consequence of stochastic variation, but can instead be attributed to its intrinsic region-specific characteristics. Consequently, asymmetric equilibrium diversity becomes uniquely associated with particular regions rather than with arbitrary permutations of region labels (see Supp. Fig. S3). For example, we derive an extension of the *Log-DDG* model that accounts for region-specific area sizes, *A*_*i*_. This extension uses relative species density, 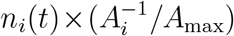, rather than raw species diversity, *n*_*i*_(*t*), to model rate variation among regions. In particular, the extinction rate of a species in region *i* at time *t*, denoted by, *e*_*i*_(*t*), is given by:

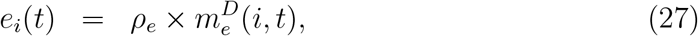

where

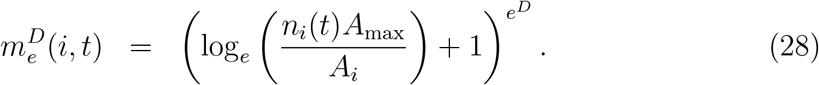

The absolute within-region speciation rate in region *i* at time *t* is:

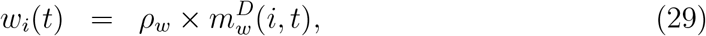

where

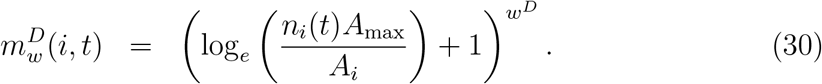

The dispersal rate from region *i* to *j* at time *t, d*_*ij*_(*t*), is defined as:

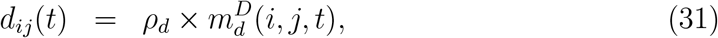

where

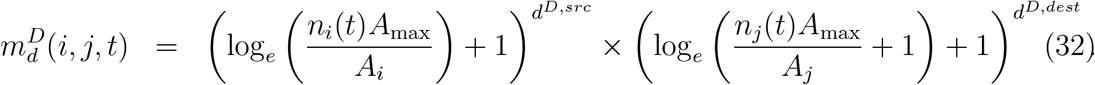

The between-region speciation rate, 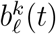, gives rise to two new species with ranges *k* ∈ *S* and *l* ∈ *S*, respectively, from an ancestral species with range *m* ∈ *S* at time *t*, where *S* is the set of possible species ranges. We define the absolute rate as follows:

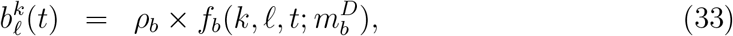

where

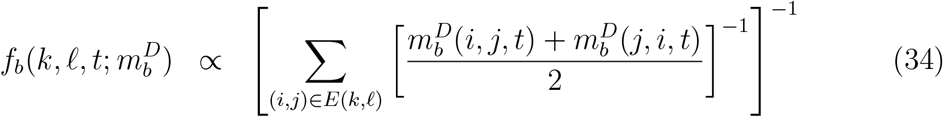

is the range split score function that depends on time *t*, species ranges *k* and *l*, and

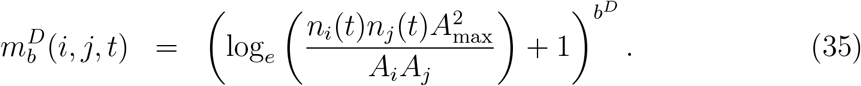

Here, *A*_*i*_ represents the area size for region *i*, and *A*_max_ denotes the maximum area across all regions. This generalization preserves the number of parameters, while allowing region-intrinsic feature (i.e., area size) to influence diversification and dispersal rates. We use relative area sizes, *A*_*i*_*/A*_max_, to ensure that the absolute rates remain well-defined.

Next, using the balance equations defined in Definition 1 together with the rate functions introduced above, we can solve the resulting system of balance equations to identify parameter values that correspond to a specified asymmetric equilibrium diversity across regions. Then, we can use the stochastic simulator to simulate trees under these parameter values, as shown in Supplemental Figure S3.

**Figure S3:**
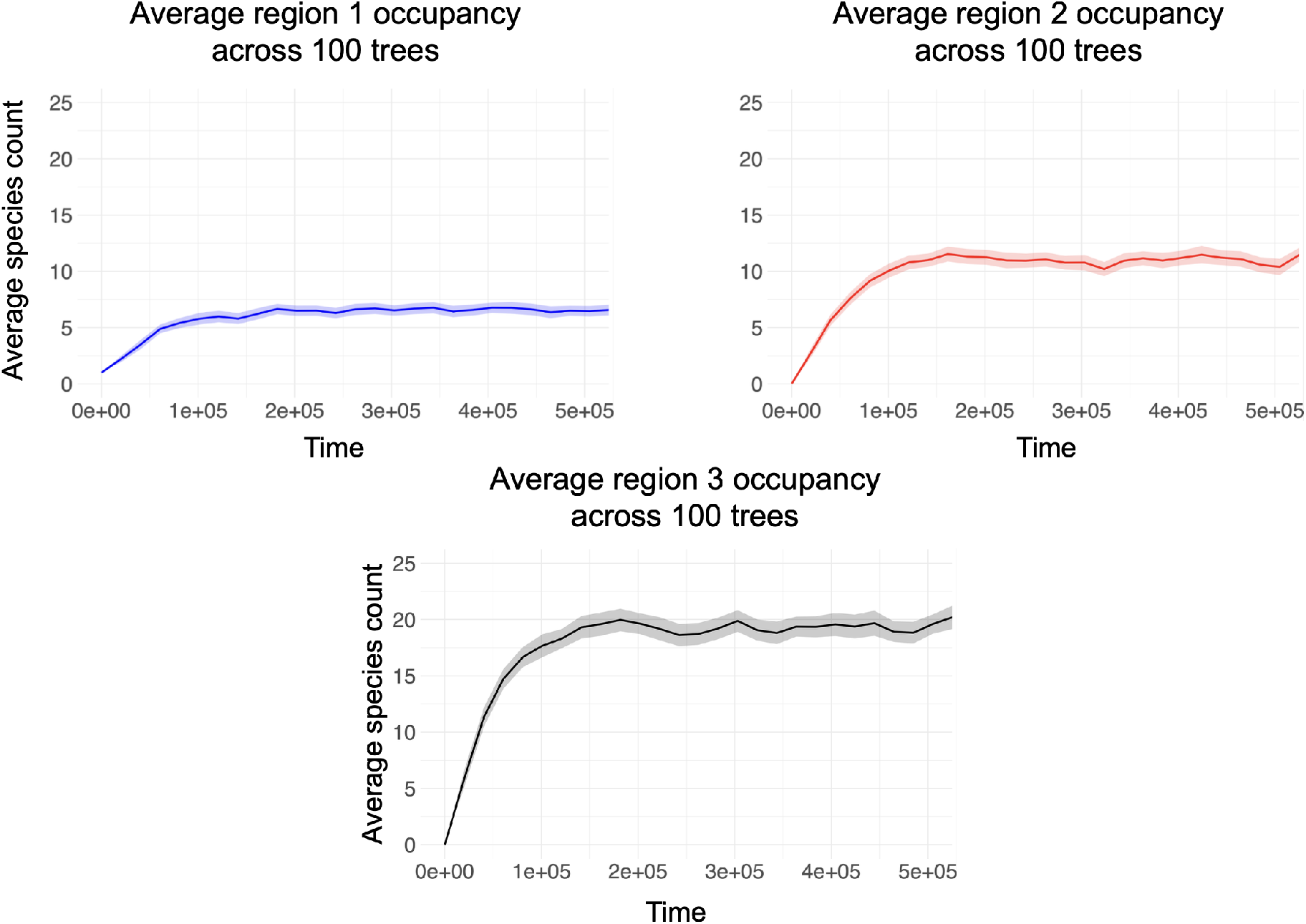
Mean trajectories of *n*_*i*_(*t*), for each of three regions and averaged across 100 replicates. Trajectories were simulated under the *Log-DDG* model variant with area size with parameter values equal to *ρ*_*w*_ = 0.0016, *ρ*_*e*_ = 4.43 *×* 10^−5^, *ρ*_*d*_ = 0.0010, *ρ*_*b*_ = 0, *w*^*D*^ = −2.69, *b*^*D*^ = 0, *d*^*D,dest*^ = −0.73, *d*^*D,src*^ = −1.75, *e*^*D*^ = 0.04, and *A*_*i*_*/A*_*max*_ = {0.01, 0.05, 1}. For each tree simulation, we initialize a tree with root species only in region 1. The shaded band around each line represents the 95% confidence interval of the mean across simulations, computed using the standard normal approximation.

## Tree statistics near the extreme case

**Figure S4:**
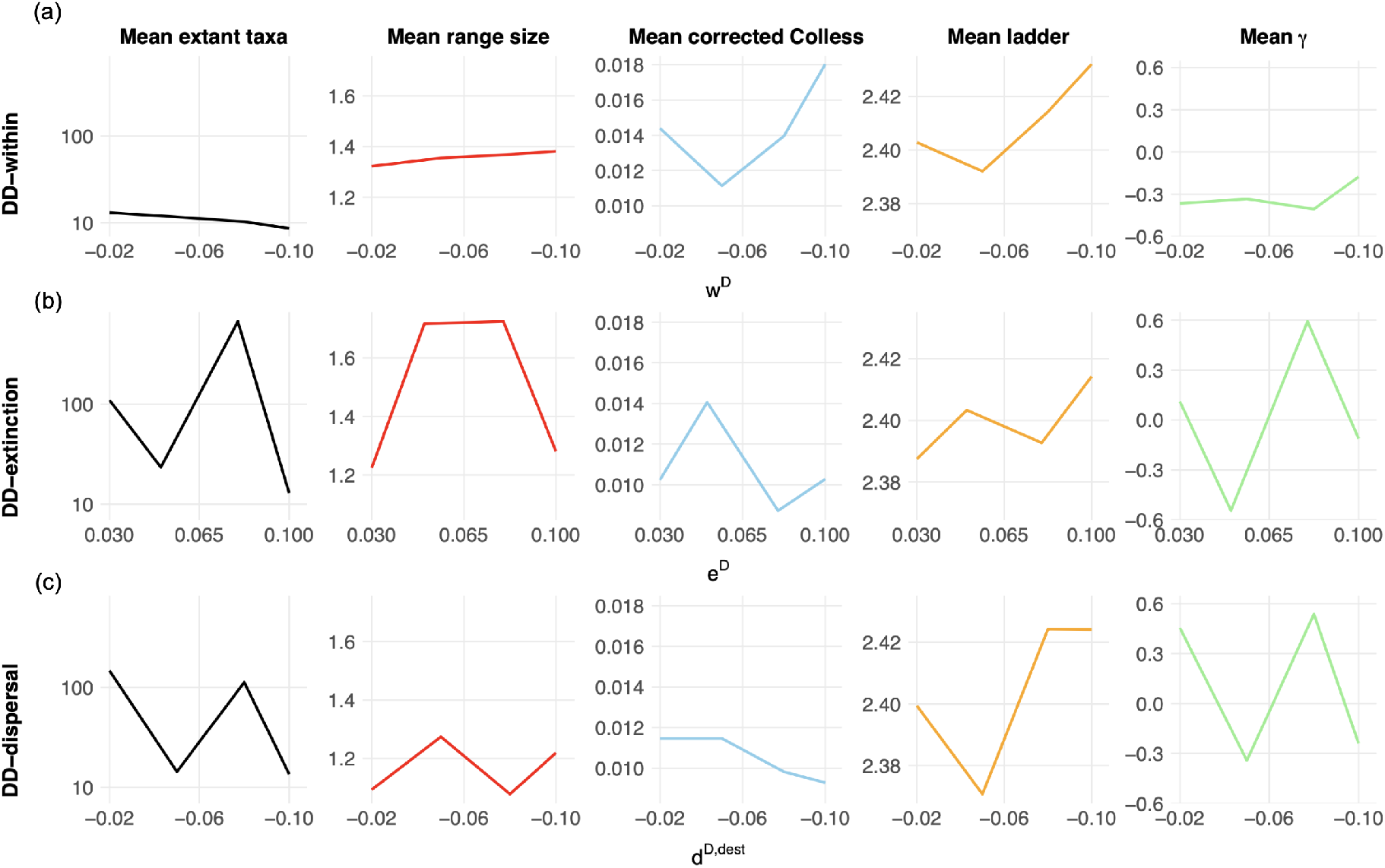
Summary statistics for simulations with diversity-dependent effects on (a) within-region speciation (“DD-within”), (b) extinction (“DD-extinction”), and (c) dispersal (“DD-dispersal”) under the extreme case scenarios for the effect of diversity-dependence on rates. The *x*-axis represents the values of each diversity-dependent effect (top: *w*^*D*^; middle: *e*^*D*^; bottom: *d*^*D,dest*^) and the *y*-axis represent the mean values of the summary statistics. A value of zero is not included on the *x*-axis because these simulations represent submodels in which the corresponding diversity-dependent effect is present; zero instead represents the absence of that effect, as represented by the corresponding alternative submodels in Table 2. The summary statistics include the mean number of extant taxa (log-scale), the mean range size (the average number of regions occupied per species), the mean corrected Colless index (tree topology; larger values indicate more imbalanced trees and smaller values indicate more balanced trees at global phylogenetic scales), the mean ladder index (tree topology; larger values indicate longer average ladder structures and smaller values indicate shorter average ladder structures at local phylogenetic scales), and the mean *γ* statistic (distribution of branching times; positive *γ* values indicate branching divergence times clustered near the present and negative *γ* values indicate branching divergence times clustered near the root).

Here, we consider the case where our Log-DDG model closely resembles the standard GeoSSE model. This can be achieved by simulating trees under the condition where the diversity-dependent effects have their values drawn from intervals near 0 *w*^*D*^, *e*^*D*^, *d*^*D,dest*^ ≈ 0.

As seen from Figure S4(a), some statistics, such as mean range size (Fig. S4(a), second column), and mean number of extant taxa (Fig. S4, first column), have similar trend to those from Figure 4(a). However, as expected, the variability in their values is less noticeable compared to those from Figure 4(a) due to *w*^*D*^ having values near 0.

As for Figure S4(b), it is noticeably harder to identify the trend of changes in these statistics values, possibly due to all *e*^*D*^ having values near 0. Similar to Figure S4(b), it is also noticeably harder to identify trend for change in values on these statistics for the model with only diversity-dependent effect on incoming dispersal (Fig. S4(c)), possibly due to *d*^*D,dest*^ has all its values drawn near 0. Overall, in all cases, the simulated extant-only trees appear relatively balanced at global scale, as shown by Colless indices having values close to 0.

## Tree shape under diversity-dependent within-region speciation and extinction

Here, we study tree shape statistics for trees simulated under our model with diversity-dependent effect on both within-region speciation and extinction. In summary, we observe consistent trends in the tree statistics when compared those using trees simulated under a single diversity-dependent effect on either processes. That is, the average range size is at the highest when *w*^*D*^ ≪ 0 and *e*^*D*^ ≈ 0 (Fig. S5, bottom left panel), and the average number of extant taxa is at the maximum when both *w*^*D*^, *e*^*D*^ ≈ 0 (Fig. S5, bottom right panel). Furthermore, consistent trends are also observed for both corrected Colless and *γ* statistics, namely the largest deviation from 0 in the Colless index occurs when *w*^*D*^ ≪ 0 and *e*^*D*^ ≫ 0 (Fig. S5, top left panel), and *γ* values tend to be larger when both *w*^*D*^, *e*^*D*^ ≈ 0 (Fig. S5, top right panel).

**Figure S5:**
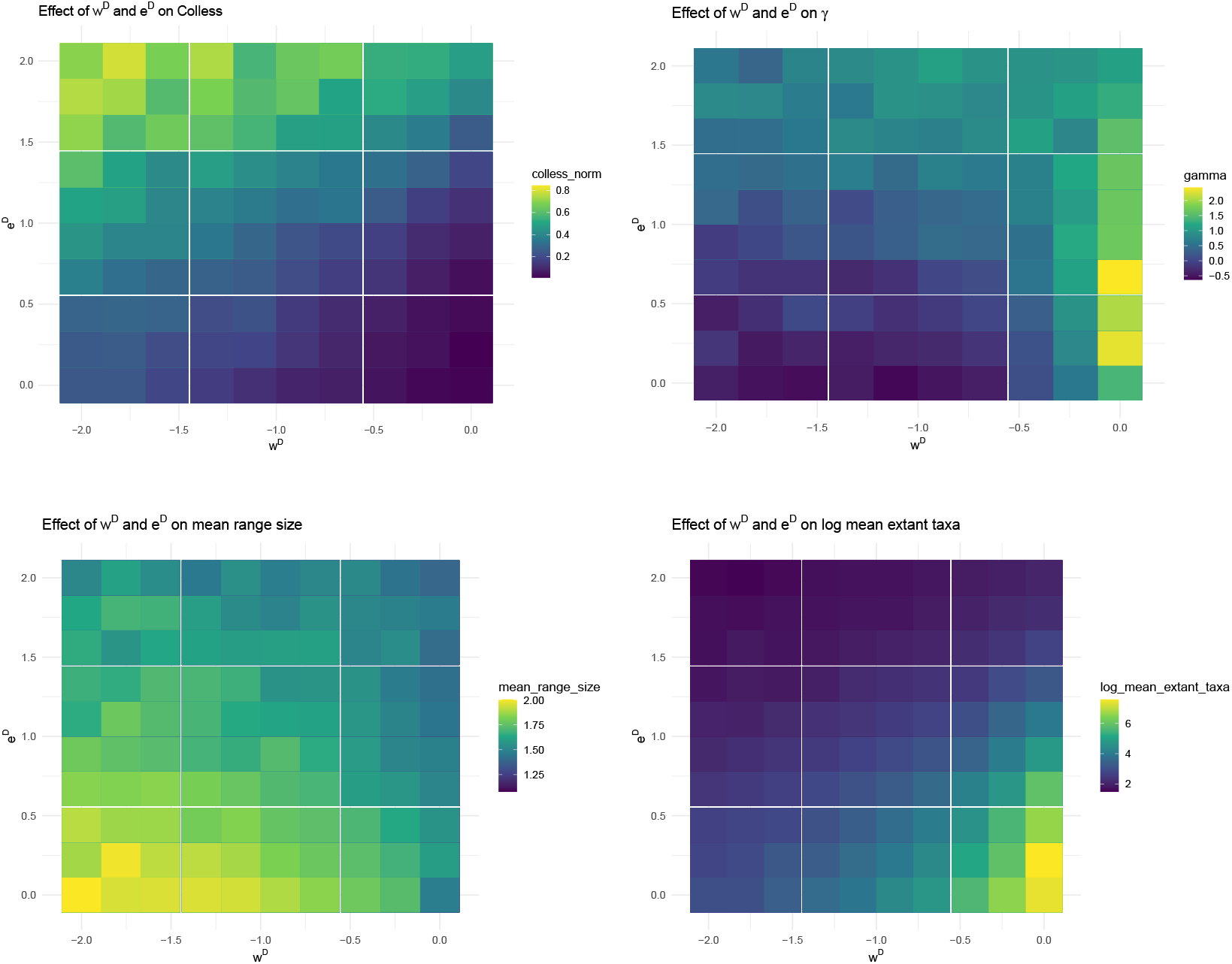
Various heatmaps showing changes in various tree statistics values due to varying degree of diversity-dependent effects on both within-region speciation and extinction.

## Tree shape under diversity-dependent incoming dispersal and extinction

**Figure S6:**
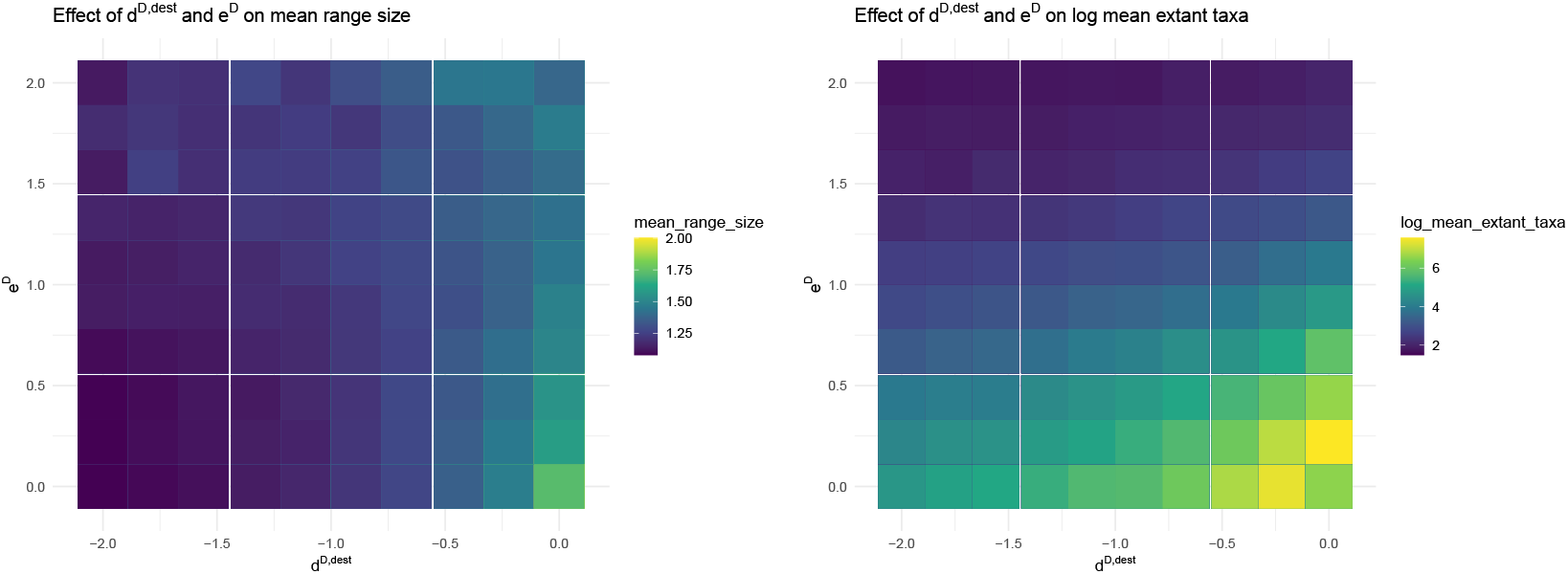
Various heatmaps showing changes in mean range size and mean number of extant taxa (in logarithmic scale) due to varying degree of diversity-dependent effects on both incoming dispersal and extinction.

Here, we study tree shape statistics for trees simulated under our model with diversity-dependent effect on both incoming dispersal and extinction. In summary, we observe the similar trend on both the average range size and average number of extant taxa when combining diversity-dependent effect on both extinction and incoming dispersal. That is, we observe higher range size and number of taxa in the region where the effect is weaker on both events (Fig. S6).

## Summary statistics for auxiliary data tensors

**Table S1:**
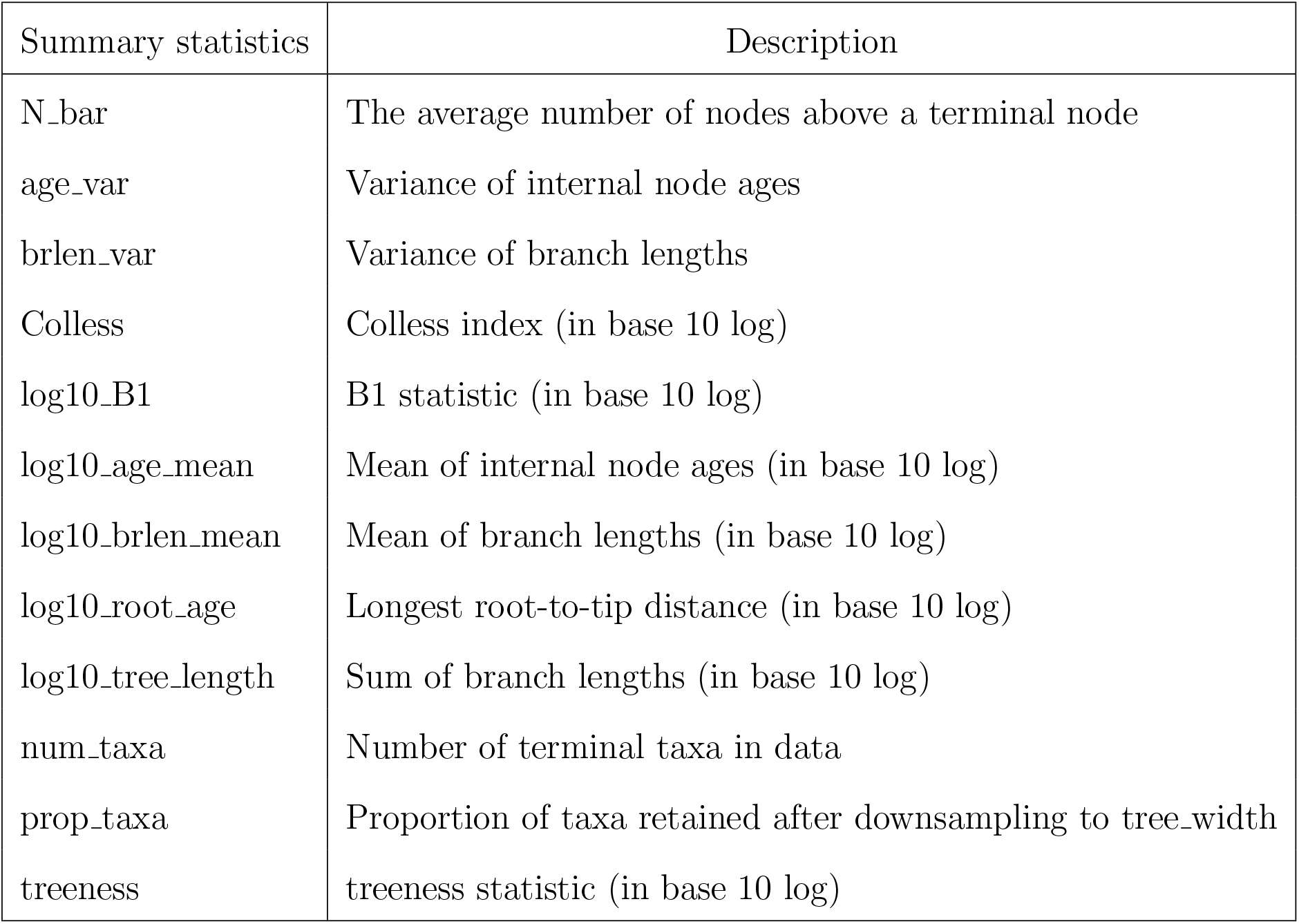
A list of summary statistics from data, encoded as auxiliary data tensors. Colless, B1, treeness, and N bar statistics are computed using Dendropy library (Sukumaran and Holder, 2010). Here, the Colless imbalance index computes the sum of differences of numbers of children in left and right subtrees over all internal nodes and the B1 statistic computes the reciprocal of the sum of the maximum number of nodes between each interior node and tip over all internal nodes excluding root.

| Summary statistics | Description |
| --- | --- |
| N_bar | The average number of nodes above a terminal node |
| age_var | Variance of internal node ages |
| brlen_var | Variance of branch lengths |
| Colless | Colless index (in base 10 log) |
| log10_B1 | B1 statistic (in base 10 log) |
| log10_age_mean | Mean of internal node ages (in base 10 log) |
| log10_brlen_mean | Mean of branch lengths (in base 10 log) |
| log10_root_age | Longest root-to-tip distance (in base 10 log) |
| log10_tree_length | Sum of branch lengths (in base 10 log) |
| num_taxa | Number of terminal taxa in data |
| prop_taxa | Proportion of taxa retained after downsampling to tree_width |
| treeness | treeness statistic (in base 10 log) |

## Performance of parameter estimation using Submodel 0

**Figure S7:**
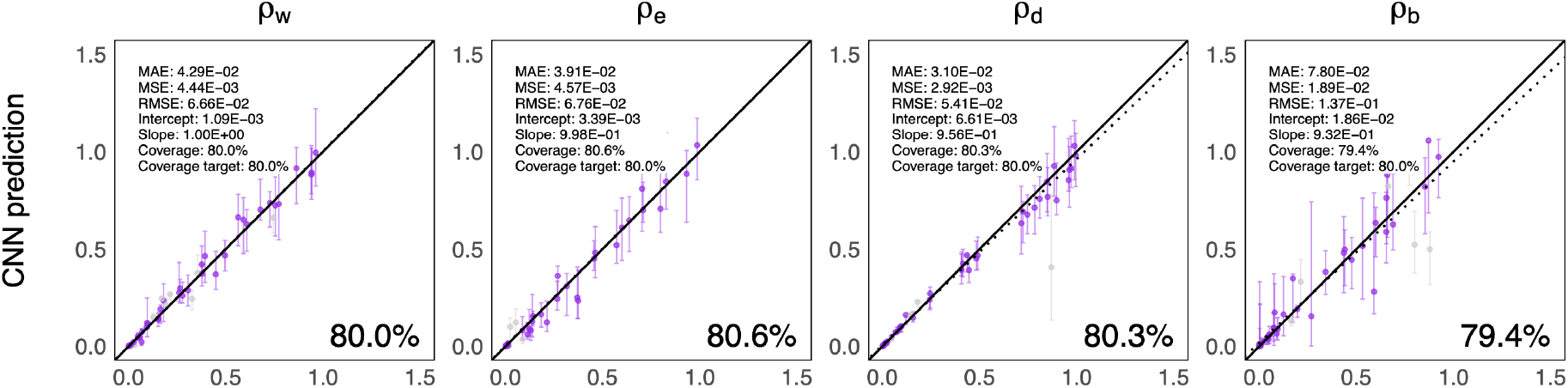
Plots showing 80% CPIs for each parameter on the test dataset simulated under submodel 0. The *x*-axis is showing the true parameter values, and the *y*-axis is showing the estimated values from CNN. All test data were used for the regression. Of these, point estimates (markers) and 80% CPIs (bars) are shown for 50 examples.

## Performance of parameter estimation using Submodel 1

**Figure S8:**
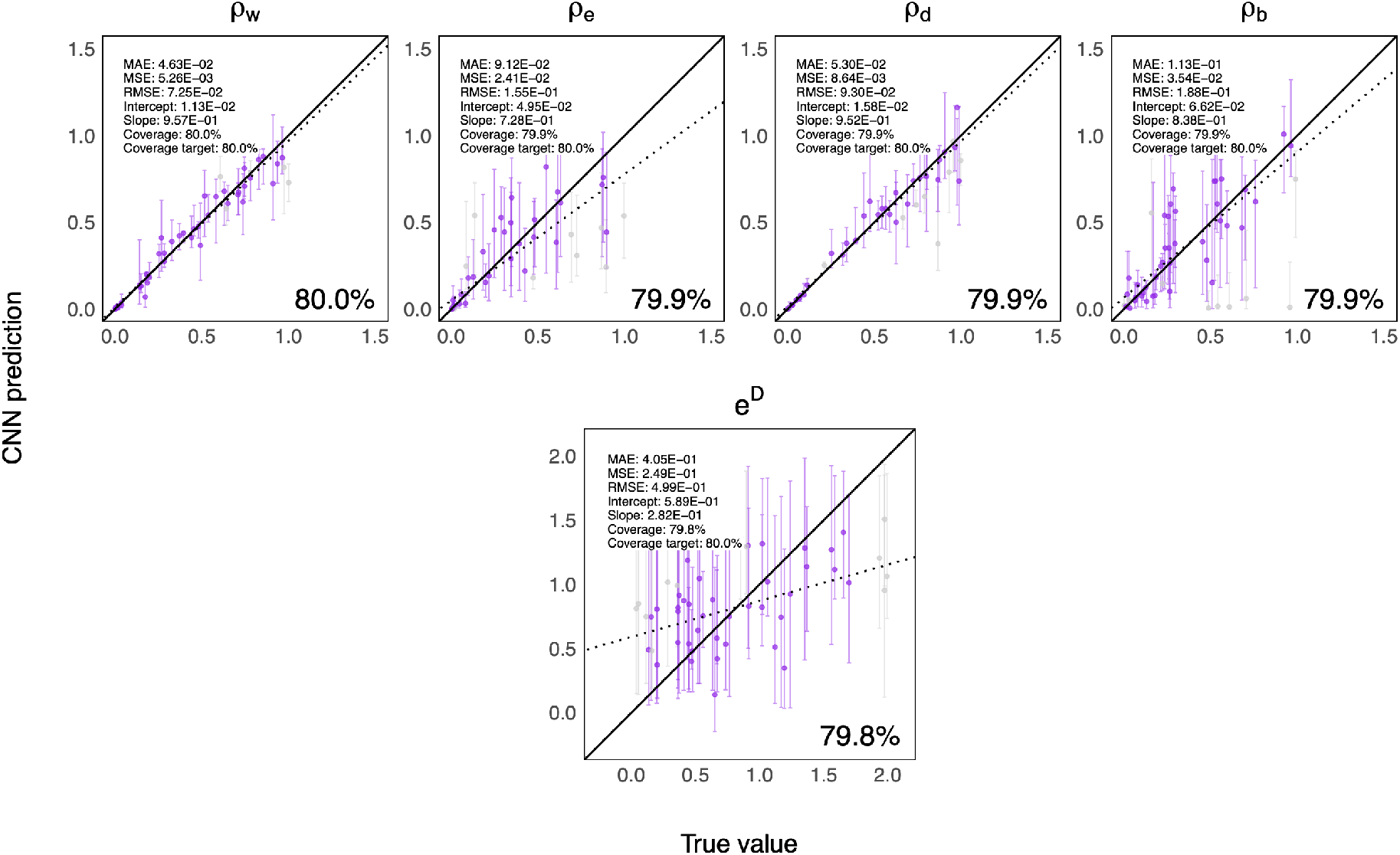
Plots showing 80% CPIs for each parameter on the test dataset simulated under submodel 1. The *x*-axis is showing the true parameter values, and the *y*-axis is showing the estimated values from CNN. All test data were used for the regression. Of these, point estimates (markers) and 80% CPIs (bars) are shown for 50 examples. The number shown on the bottom right of each panel indicates the percentage of test cases where the true parameter value falls inside the 80% CPI.

## Performance of parameter estimation using Submodel 2

**Figure S9:**
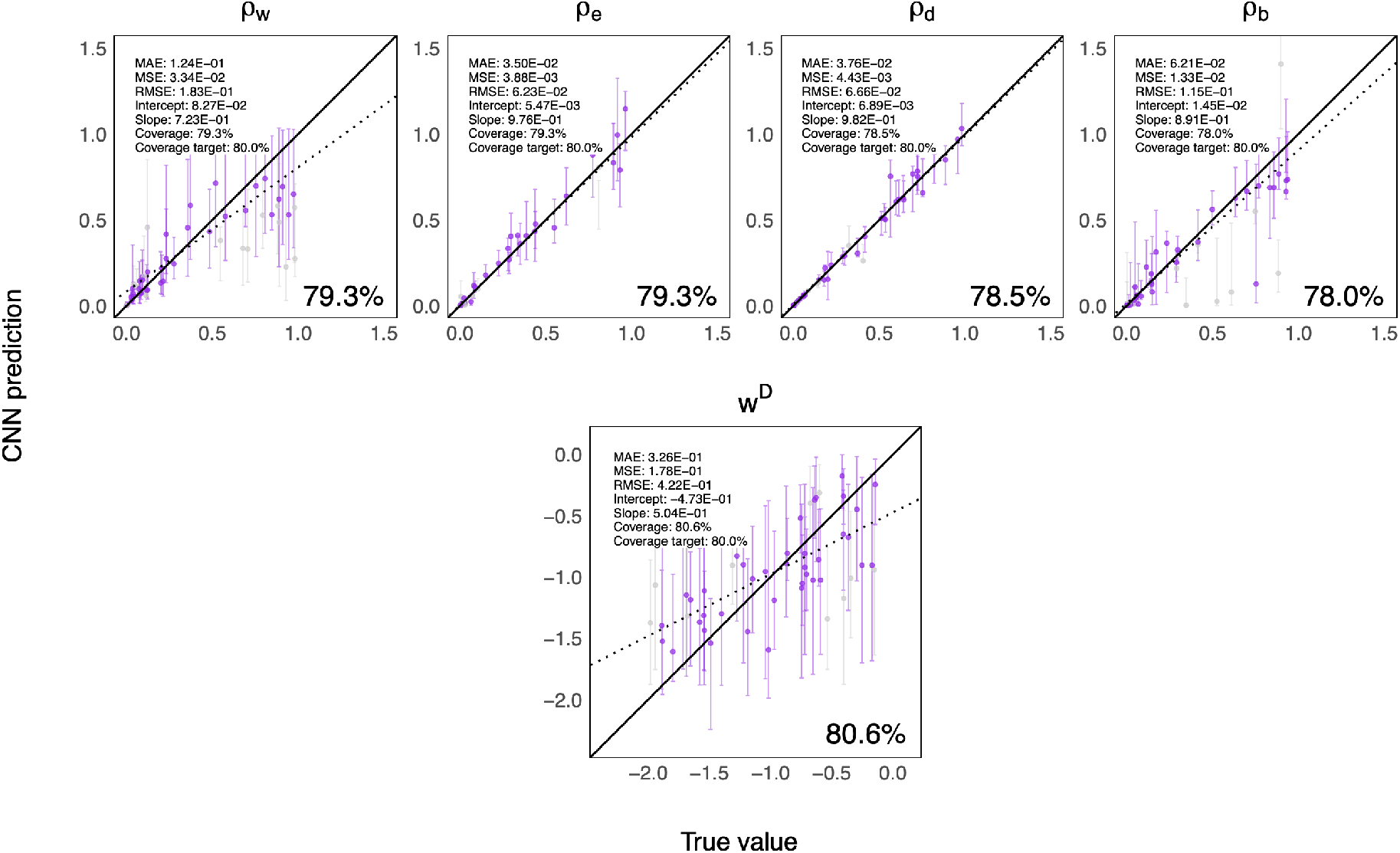
Plots showing 80% CPIs for each parameter on the test dataset simulated under submodel 2. The *x*-axis is showing the true parameter values, and the *y*-axis is showing the estimated values from CNN. All test data were used for the regression. Of these, point estimates (markers) and 80% CPIs (bars) are shown for 50 examples. The number shown on the bottom right of each panel indicates the percentage of test cases where the true parameter value falls inside the 80% CPI.

## Performance of parameter estimation using Submodel 3

**Figure S10:**
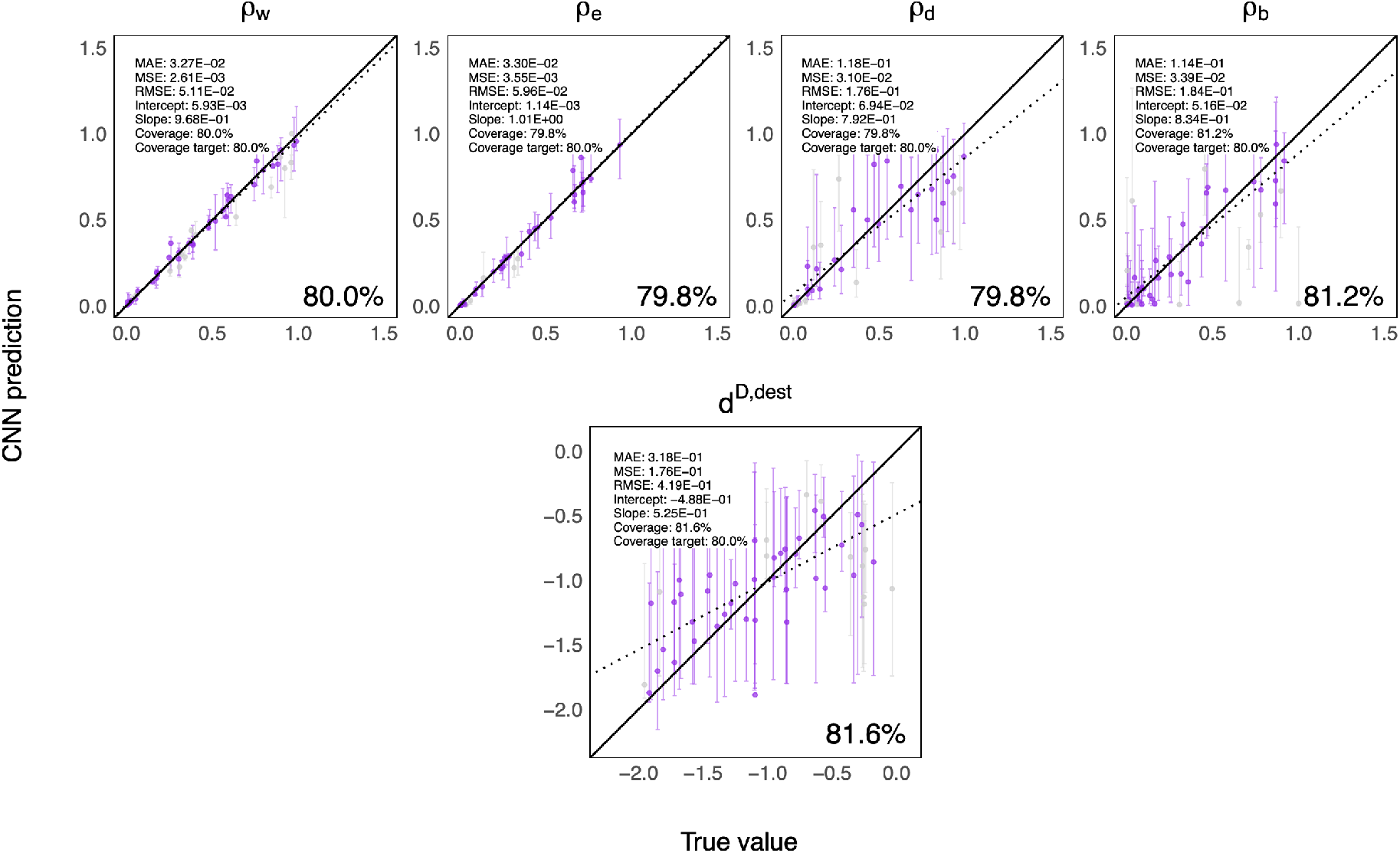
Plots showing 80% CPIs for each parameter on the test dataset simulated under submodel 3. The *x*-axis is showing the true parameter values, and the *y*-axis is showing the estimated values from CNN. All test data were used for the regression. Of these, point estimates (markers) and 80% CPIs (bars) are shown for 50 examples. The number shown on the bottom right of each panel indicates the percentage of test cases where the true parameter value falls inside the 80% CPI.

## Performance of parameter estimation using Submodel 4

**Figure S11:**
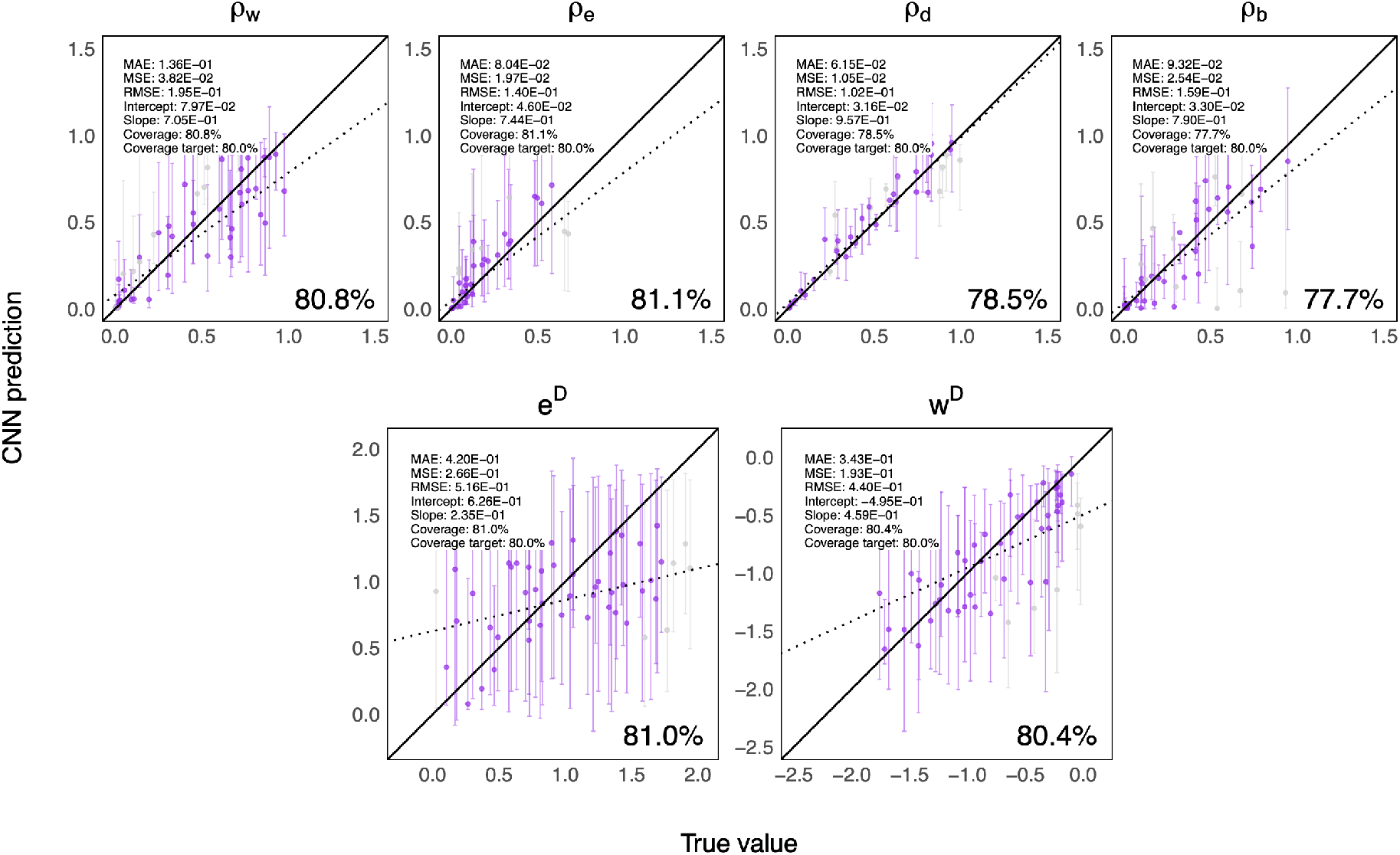
Plots showing 80% CPIs for each parameter on the test dataset simulated under submodel 4. The *x*-axis is showing the true parameter values, and the *y*-axis is showing the estimated values from CNN. All test data were used for the regression. Of these, point estimates (markers) and 80% CPIs (bars) are shown for 50 examples. The number shown on the bottom right of each panel indicates the percentage of test cases where the true parameter value falls inside the 80% CPI.

## Performance of parameter estimation using Submodel 5

**Figure S12:**
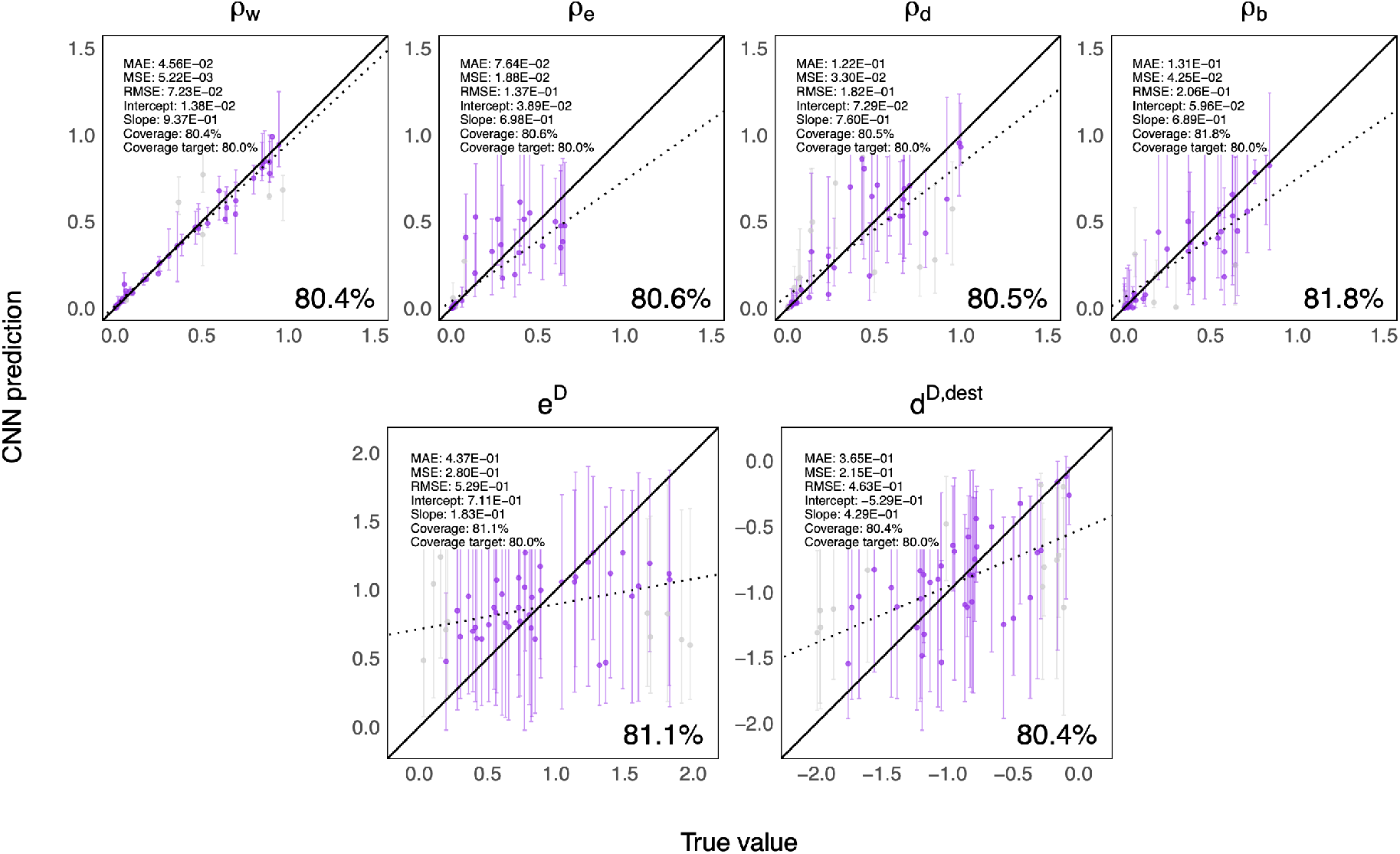
Plots showing 80% CPIs for each parameter on the test dataset simulated under submodel 5. The *x*-axis is showing the true parameter values, and the *y*-axis is showing the estimated values from CNN. All test data were used for the regression. Of these, point estimates (markers) and 80% CPIs (bars) are shown for 50 examples. The number shown on the bottom right of each panel indicates the percentage of test cases where the true parameter value falls inside the 80% CPI.

## Performance of parameter estimation using Submodel 6

**Figure S13:**
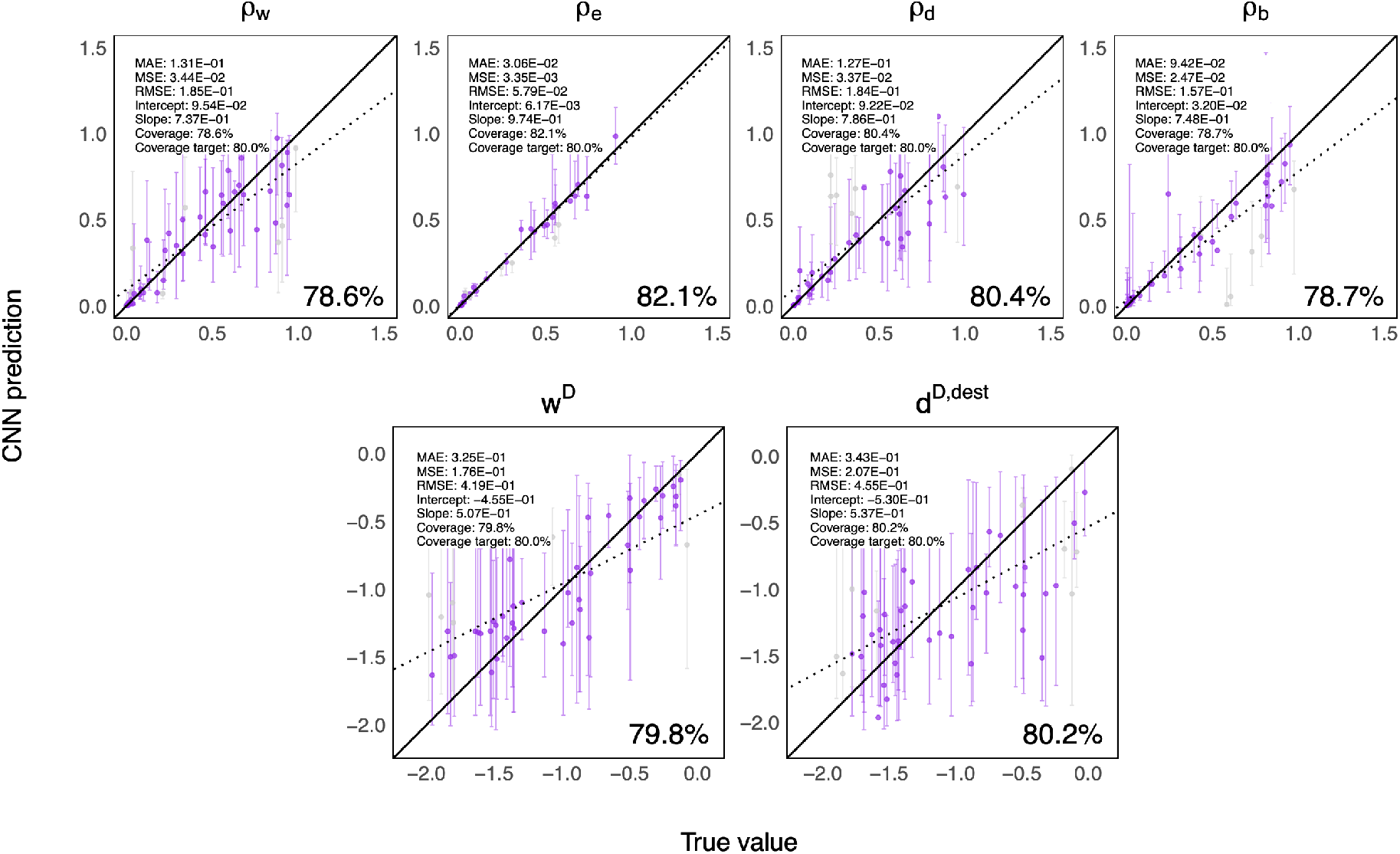
Plots showing 80% CPIs for each parameter on the test dataset simulated under submodel 6. The *x*-axis is showing the true parameter values, and the *y*-axis is showing the estimated values from CNN. All test data were used for the regression. Of these, point estimates (markers) and 80% CPIs (bars) are shown for 50 examples. The number shown on the bottom right of each panel indicates the percentage of test cases where the true parameter value falls inside the 80% CPI.

## Deep learning accuracy for empirical estimation

**Table S2:** DDGeoSSE estimates for Caribbean *Anolis* lizards and Neotropical *Oreinotinus* plants using phyddle. Rows 1-7 show point estimates and 80% CPIs (in brackets) for the best-fitting model. Rows 1-4 show base rates for within-region speciation (*ρ*_*w*_), extinction (*ρ*_*e*_), dispersal (*ρ*_*d*_), and between-region speciation (*ρ*_*b*_). Rows 5-7 show diversity-dependent effect parameters for within-region speciation *w*^*D*^, extinction *e*^*D*^, and incoming dispersal *d*^*D,dest*^. Note that the base rates for *Viburnum* have been converted to correspond to the original phylogeny (branch lengths are measured in unit time). The last column shows the loss score calculated from each model’s corresponding validation dataset. The empirical estimates and loss score from each model are comparable to the results shown by another set of independent networks trained on the same datasets on Table 4.

|  | Estimate | *Anolis* |  | *Viburnum* |  |
| --- | --- | --- | --- | --- | --- |
| Parameter estimation for best-fitting model | $\rho_w$ | 0.2724 | [0.1049, 0.6722] | 1.6401 | [0.5831, 2.4766] |
| | $\rho_e$ | 0.0003 | [0.0001, 0.0016] | 0.0021 | [0.00063, 0.0161] |
| | $\rho_d$ | 0.0009 | [0.0002, 0.0029] | 0.0021 | [0.00056, 0.0686] |
| | $\rho_b$ | 0.0842 | [0.0005, 0.3391] | 0.0266 | [0.00063, 2.8511] |
| | $w^D$ | -1.55 | [-1.83, -0.79] | -1.90 | [-2.18, -0.77] |
| | $e^D$ | 0.80 | [0.21, 1.76] | — | — |
| | $d^{D,dest}$ | -1.25 | [-1.87, -0.36] | -1.30 | [-2.00, -0.37] |
| **Validation loss score** |  | 0.609 |  | 0.487 |  |

## Empirical rates under best-fitting models trained networks without weight_decay

### Empirical rates under best-fitting models from trained networks with weight_decay

**Figure S14:**
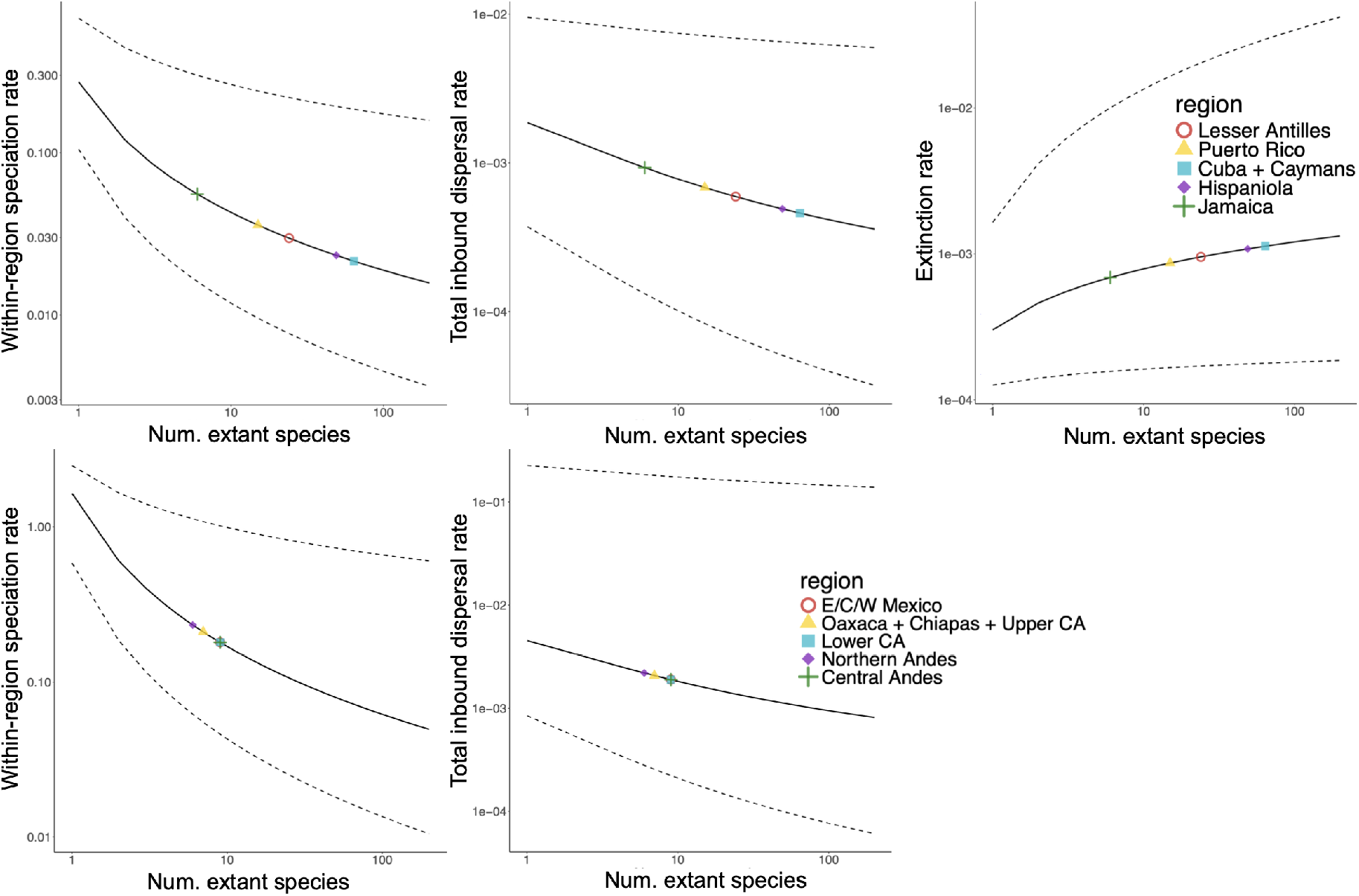
Empirical rates for *Anolis* (top panel) and *Oreinotinus* (bottom panel) using their best-fitting models, submodel 7 for *Anolis* and submodel 6 for *Oreinotinus*, with the networks trained with weight decay enabled. The *x*-axis shows the number of extant species and the *y*-shows the overall rates for each event: within-region speciation rates (left panel), total inbound dispersal rates (middle panel), and extinction rates (right panel). The total inbound dispersal rate for region *i* is computed using the formula: ∑_*j*_ *d*_*ji*_. The colored symbols show the region-dependent empirical rates for each process for the two empirical datasets. Their overall rates are computed using values shown in Table S2. The extinction rate for *Oreinotinus* is not plotted since it is equal to its base rate for all regions under submodel 6.

## Deep learning accuracy for model selection using simulated dataset – I

**Figure S15:**
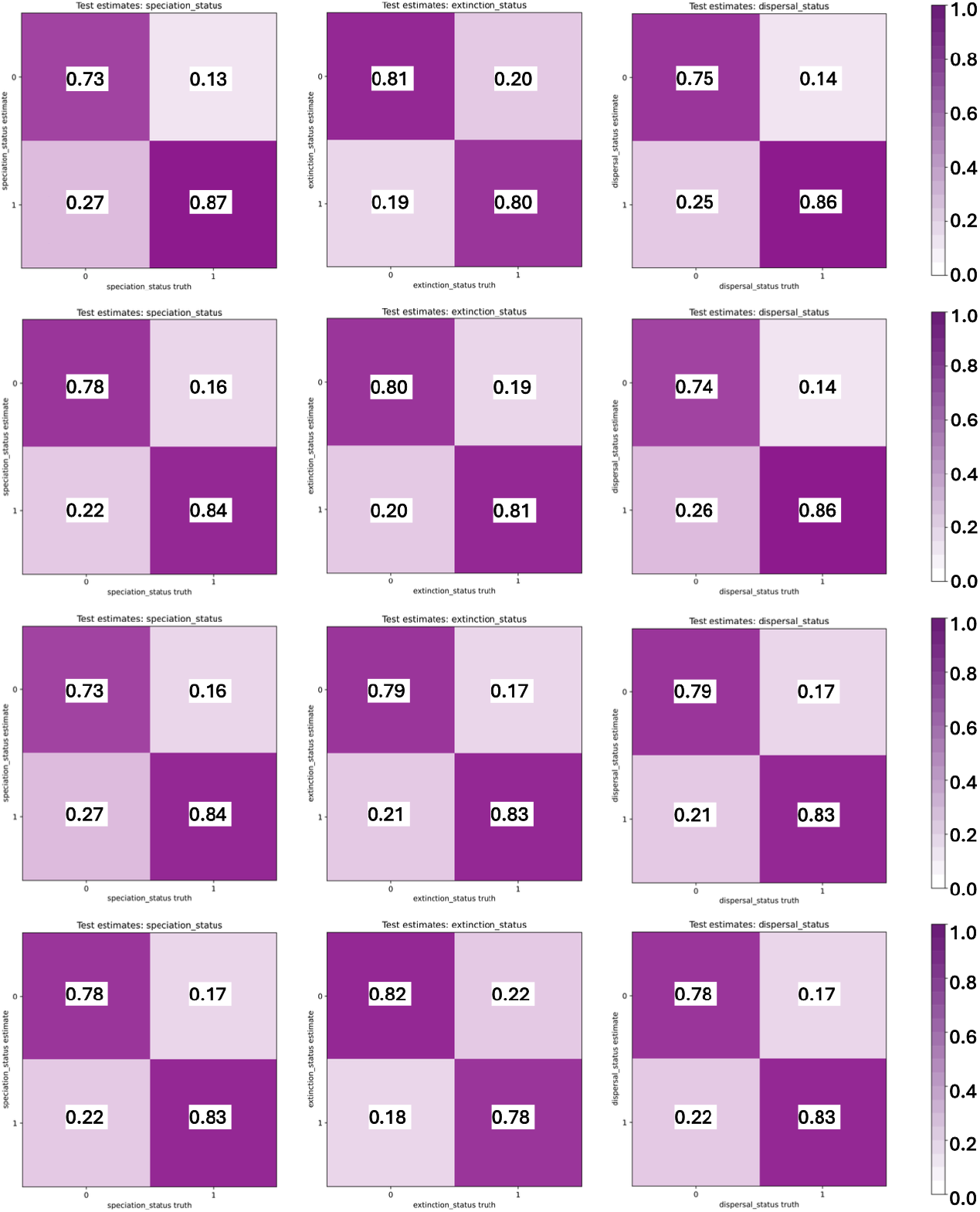
Performance on the test dataset using four independently trained networks for each training target on 400,000 simulations, generated by 8 submodels as described in Table 2 (50,000 each), for detecting the presence (1) or absence (0) of diversity dependence in within-region speciation (left panel), extinction (middle panel), and dispersal (right panel) separately. In total, we have 24 independently trained networks. The *x*-axis shows the true scenario, and the *y*-axis shows the predicted scenario.

## Deep learning accuracy for model selection using simulated dataset – II

**Figure S16:**
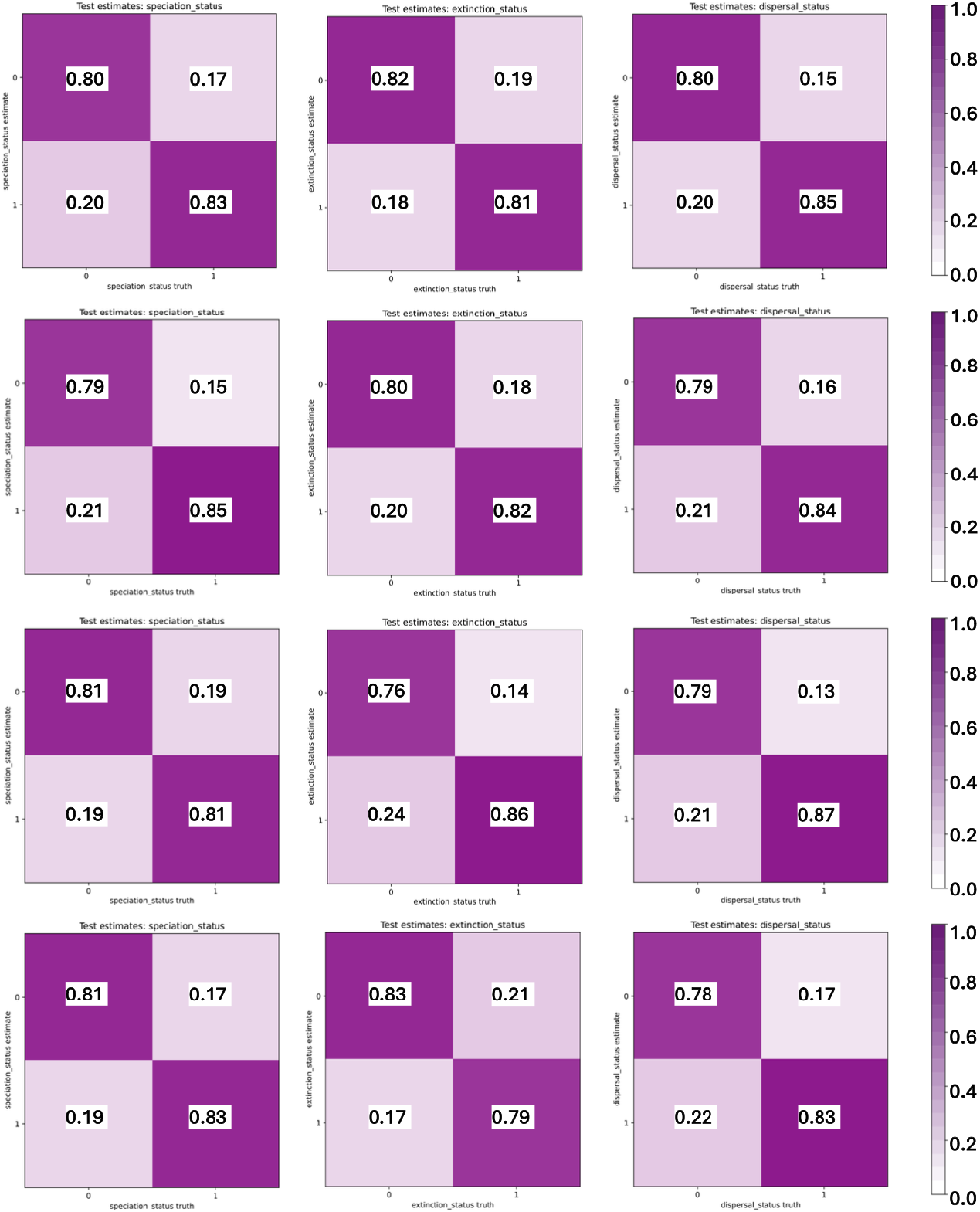
Performance on the test dataset using another set of four independently trained networks for each training target on 400,000 simulations, generated by 8 submodels as described in Table 2 (50,000 each), for detecting the presence (1) or absence (0) of diversity dependence in within-region speciation (left panel), extinction (middle panel), and dispersal (right panel) separately. In total, we have 24 independently trained networks. The *x*-axis shows the true scenario, and the *y*-axis shows the predicted scenario.

**Figure S17:**
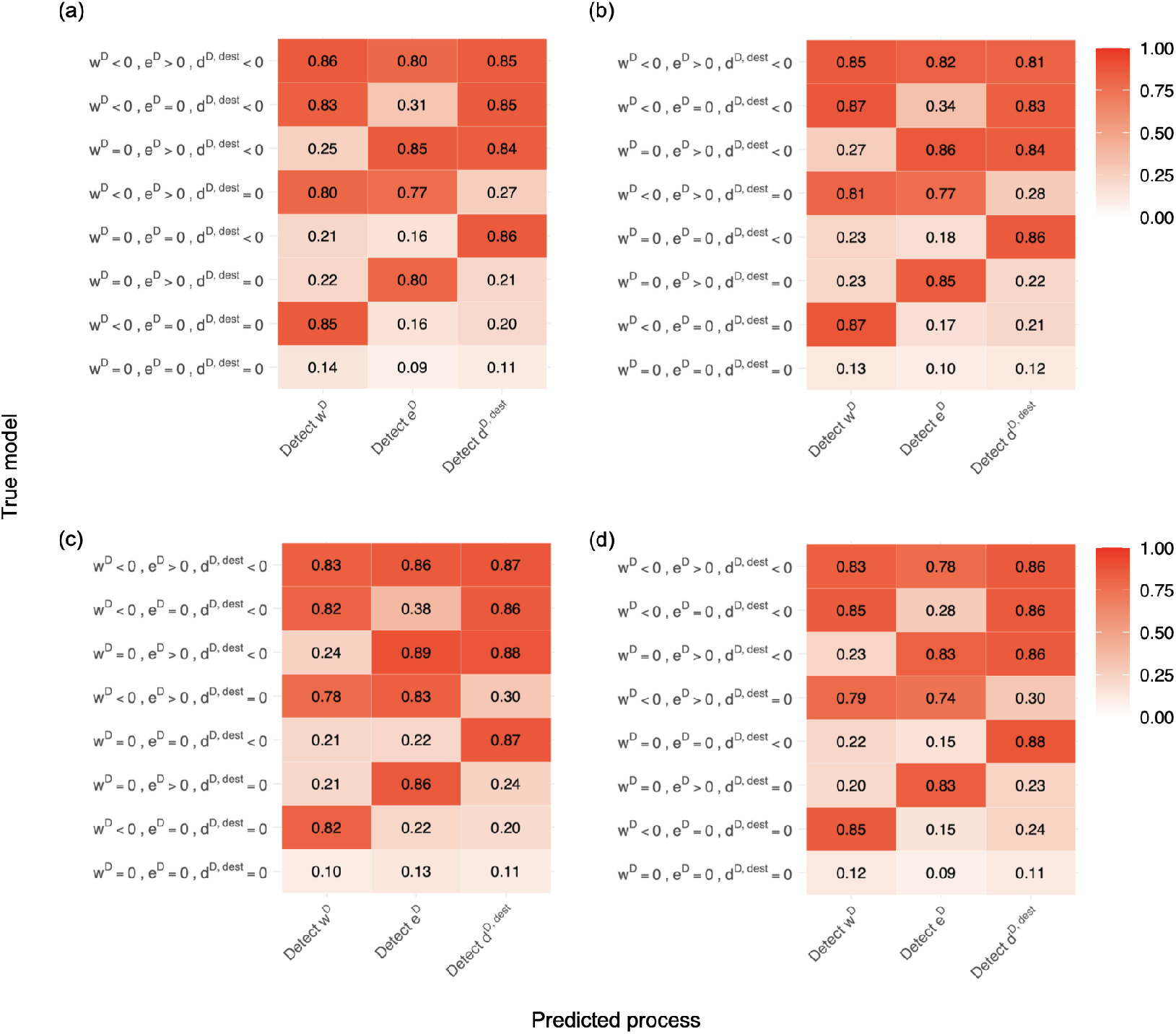
Process-level classification performance across test datasets under different submodels, which are categorized based on their properties of presence or absence of diversity-dependence on within-region speciation (*w*^*D*^), extinction (*e*^*D*^), and incoming dispersal (*d*^*D,dest*^). Rows correspond to the true generating submodel, while columns show the proportion of datasets for which the corresponding process-level classifier detects diversity dependence on within-region speciation, extinction, or incoming dispersal, using a predicted marginal probability threshold of 0.5. Panels (a)-(d) correspond to 4 sets of independent networks (networks 5-8) trained using weight decay option of 0.001 in phyddle. Higher values indicate stronger support for diversity dependence in the corresponding process. Overall, the classifiers generally show high support for processes that are truly diversity dependent and lower support for processes that are absent.

## Sensitivity analyses under model misspecification

We simulated 2,500 trees under a variant of Submodels 1–7 (Table 2) that incorporates relative area size, using the simulation settings described in Table 3 and the relative area-size vector {0.1261, 0.0802, 1, 0.6892, 0.0990}. We then estimated the model parameters using neural networks trained on datasets simulated under the original submodels in Table 2, which do not account for relative area size. As seen in Figures S18-S24, several samples from datasets simulated under the more complex models have their tree summary statistics (encoded as auxiliary data tensor; Table S1) outside the training distribution.

As observed in Table S3, under correct model specification, the achieved CPI levels are generally close to the 80% CPI target level, indicating accurate parameter estimation. However, under model misspecification, coverage deteriorates for several parameters, particularly for base rates the diversity-dependent effect is present and *d*^*D,dest*^, with achieved CPI levels as low as 64 − 70%. These values fall well below the 80% target and outside the expected binomial confidence interval. Nevertheless, coverage remains relatively close to the set level for several other parameters, namely *w*^*D*^ and *e*^*D*^.

Overall, this deterioration in CPI coverage is expected under model misspecification, because the neural networks were trained on simulations generated without relative area size, whereas the misspecified datasets were generated under a model that includes relative area size. The results are consistent with the observation that several samples from the more complex models are out-of-distribution, suggesting that the networks cannot be trusted to make accurate predictions for these samples.

**Table S3.** Parameter estimation accuracy under model misspecification and correct model specification. Each neural network was trained using data simulated under its corresponding submodel without incorporating relative area size, as described in Table 3. For the analysis without relative area size, we evaluated performance using a test dataset containing 2,500 simulations from each submodel and assessed the achieved CPI level. For the analysis with relative area size, performance was evaluated using a test dataset containing 2,500 simulations generated under a variant of each submodel that incorporates relative area size (see the section *Model Variant: Log-DDG Model with Relative Area Size*). Bold values indicate achieved CPI levels that fall outside the expected 95% confidence interval for a binomial experiment with 2,500 trials and a coverage probability of 0.80.

| Options | Submodel | Without relative area size |  |  |  |  |  |  | With relative area size |  |  |  |  |  |  |
| --- | --- | --- | --- | --- | --- | --- | --- | --- | --- | --- | --- | --- | --- | --- | --- |
| | | $\rho_w$ | $\rho_e$ | $\rho_d$ | $\rho_b$ | $w^D$ | $e^D$ | $d^{D,dest}$ | $\rho_w$ | $\rho_e$ | $\rho_d$ | $\rho_b$ | $w^D$ | $e^D$ | $d^{D,dest}$ |
| Without weight_decay | Submodel 1 | 80.0% | 79.9% | 79.9% | 79.9% | - | 79.8% | - | **77.8%** | **74%** | 80.2% | 79% | - | 79.2% | - |
|  | Submodel 2 | 79.3% | 79.3% | 78.5% | **78.0%** | 80.6% | - | - | **69.9%** | 79.1% | 79% | 81.3% | 79.7% | - | - |
|  | Submodel 3 | 80.0% | 79.8% | 79.8% | 81.2% | - | - | 81.6% | 84.2% | 84.4% | **64.5%** | 80.7% | - | - | **70.4%** |
|  | Submodel 4 | 80.8% | 81.1% | 78.5% | **77.7%** | 80.4% | 81.0% | - | **70.4%** | **73.3%** | 80.4% | 79.5% | 80.6% | 79.6% | - |
|  | Submodel 5 | 80.4% | 80.6% | 80.5% | **81.8%** | - | 81.1% | 80.4% | 80.9% | **73.6%** | **72.4%** | 78.9% | - | 79.8% | **84.5%** |
|  | Submodel 6 | 78.6% | **82.1%** | 80.4% | 78.7% | 79.7% | - | 80.2 | **70.8%** | 79.0% | **69.4%** | 80.2% | 80.2% | - | **84.4%** |
|  | Submodel 7 | 80.3% | 79.1% | 81.2% | 80.8% | 79.3% | 80.0% | 80.1% | **70.2%** | **71.8%** | **70.3%** | 80.0% | 80.7% | **78.4%** | **82.8%** |
| With weight_decay = 0.001 | Submodel 1 | 80.0% | 80.4% | 81.1% | 80.4% | - | 80.2% | - | 80.3% | **74.4%** | 79.4% | 79.5% | - | 80.6% | - |
|  | Submodel 2 | 79.9% | 80.7% | 79.6% | 79.4% | 81.0% | - | - | **67.6%** | 80.9% | **78.4%** | 78.6% | 81.3% | - | - |
|  | Submodel 3 | 80.9% | 79.9% | 78.8% | 80.2% | - | - | 80.4% | 80.4% | 78.6% | **64.6%** | **82.0%** | - | - | **66.0%** |
|  | Submodel 4 | 81.0% | 81.2% | 79.4% | 79.6% | 81.1% | 81.2% | - | **69.5%** | **75%** | 78.7% | 81.3% | **81.7%** | 80.8% | - |
|  | Submodel 5 | 80.0% | 80.4% | 79.6% | 81.4% | - | 80.4% | 80.7% | 80.1% | **73.4%** | **70.8%** | 80.5% | - | 78.5% | **83.8%** |
|  | Submodel 6 | 79.2% | 80.4% | 79.6% | 81.0% | 79.2% | - | 80.9% | **70.4%** | 79.7% | **70.2%** | 79.4% | 80% | - | **83.1%** |
|  | Submodel 7 | 78.9% | 80.2% | 81.0% | 81.0% | 79.6% | 81.1% | 78.6% | **71.9%** | **73.4%** | **72.0%** | 79.1% | **82.9%** | 80.4% | **82.6%** |

**Figure S18:**
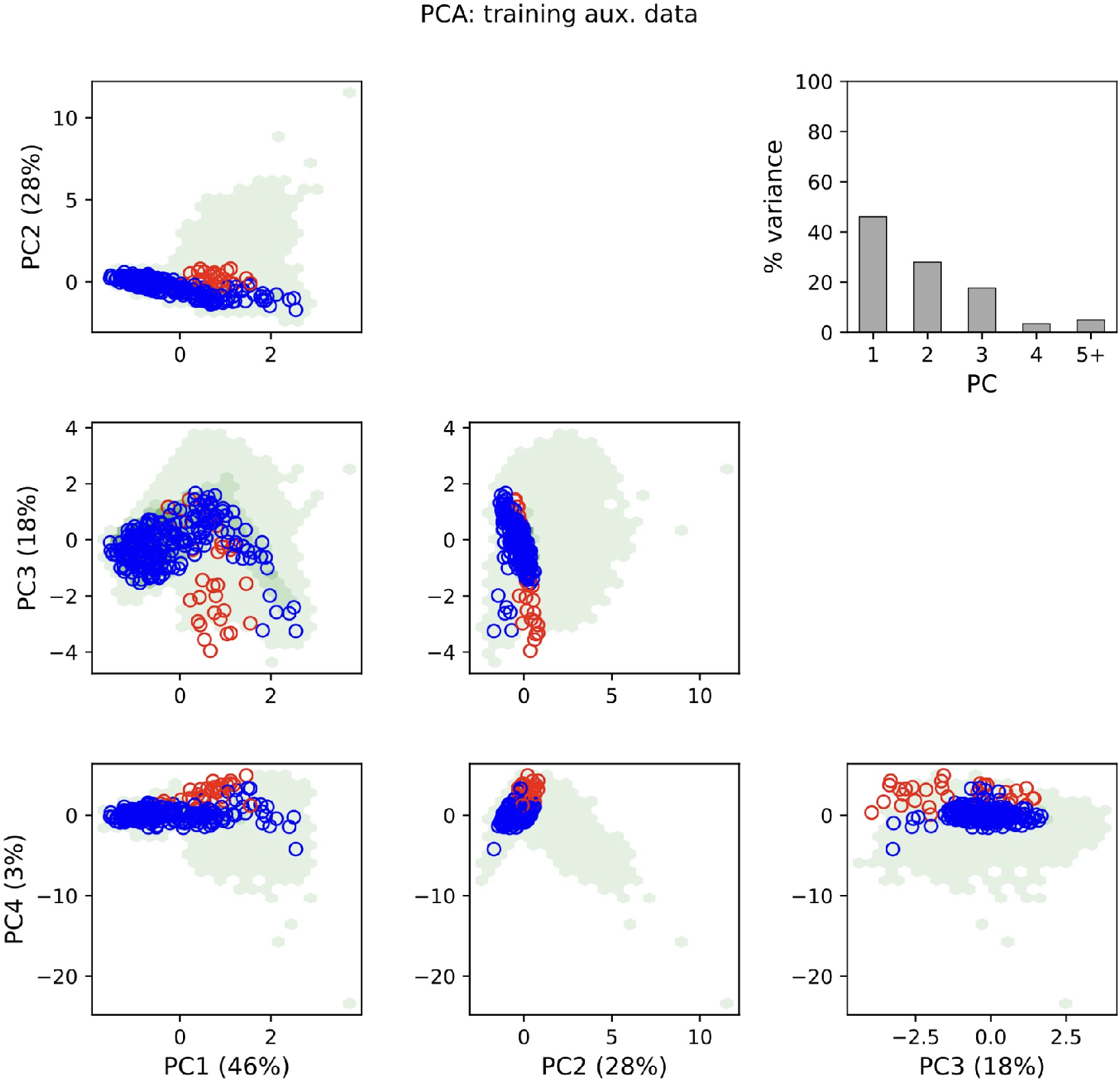
PCA plots for the auxiliary data tensor for data simulated with relative area size under Submodel 1. The blue and red circles correspond to a random subsample of size 250 of the summary statistics from the simulated dataset simulated under a variant of Submodel 1 that accounts for relative area size. The red circles represent simulations identified as outliers relative to the training dataset generated under Submodel 1 without relative area size, whereas the blue circles represent the remaining simulations. In total, 13.2% of the samples were categorized as outliers. These outliers indicate a shift in distribution between the two datasets and may contribute to reduced CPI coverage under model misspecification.

**Figure S19:**
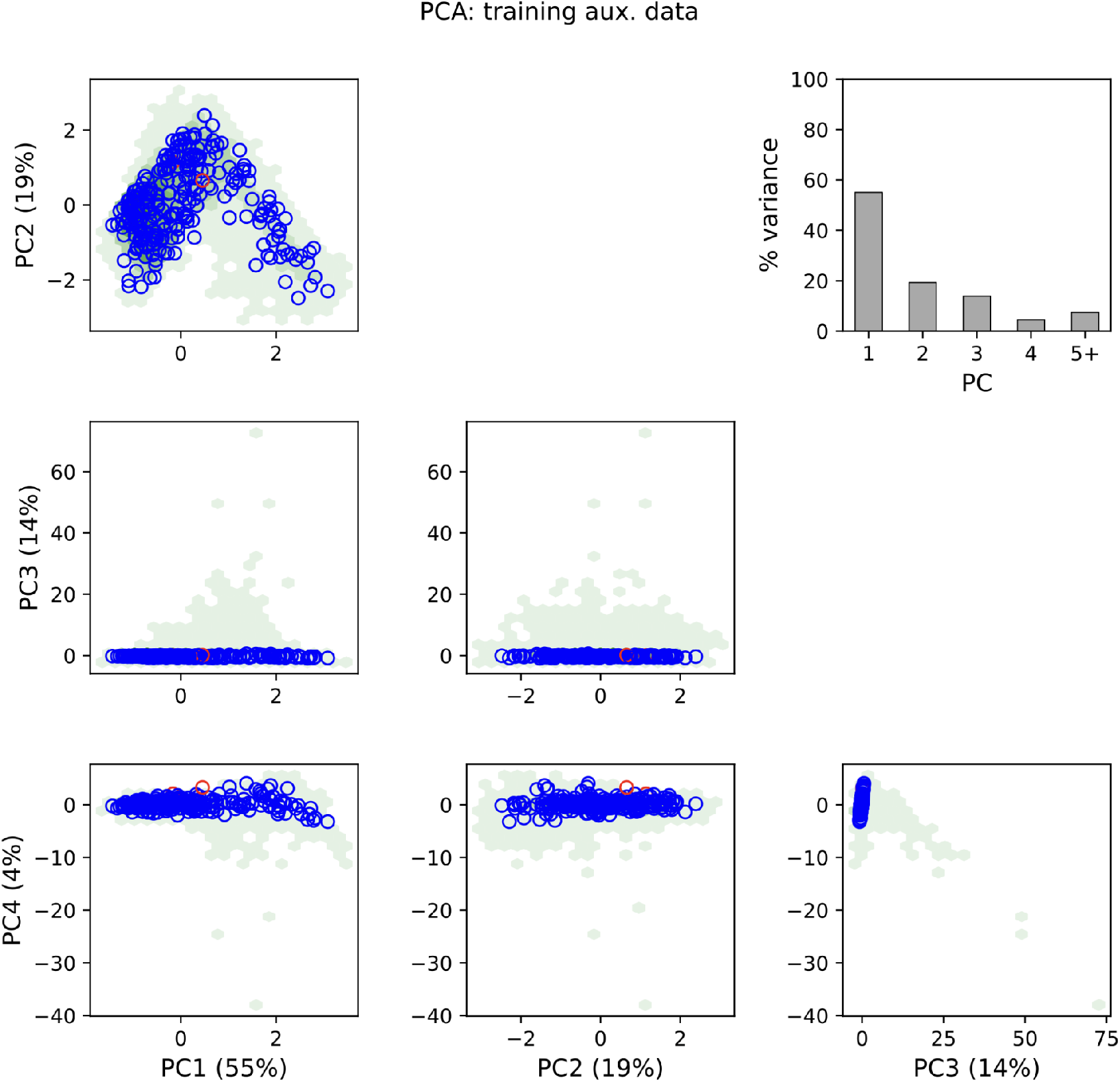
PCA plots for the auxiliary data tensor for data simulated with relative area size under Submodel 2. The blue and red circles correspond to a random subsample of size 250 of the summary statistics from the dataset simulated under a variant of Submodel 2 that accounts for relative area size. The red circles represent simulations identified as outliers relative to the training dataset generated under Submodel 2 without relative area size, whereas the blue circles represent the remaining simulations. In total, 2% of the samples were categorized as outliers. These outliers indicate a shift in distribution between the two datasets and may contribute to reduced CPI coverage under model misspecification.

**Figure S20:**
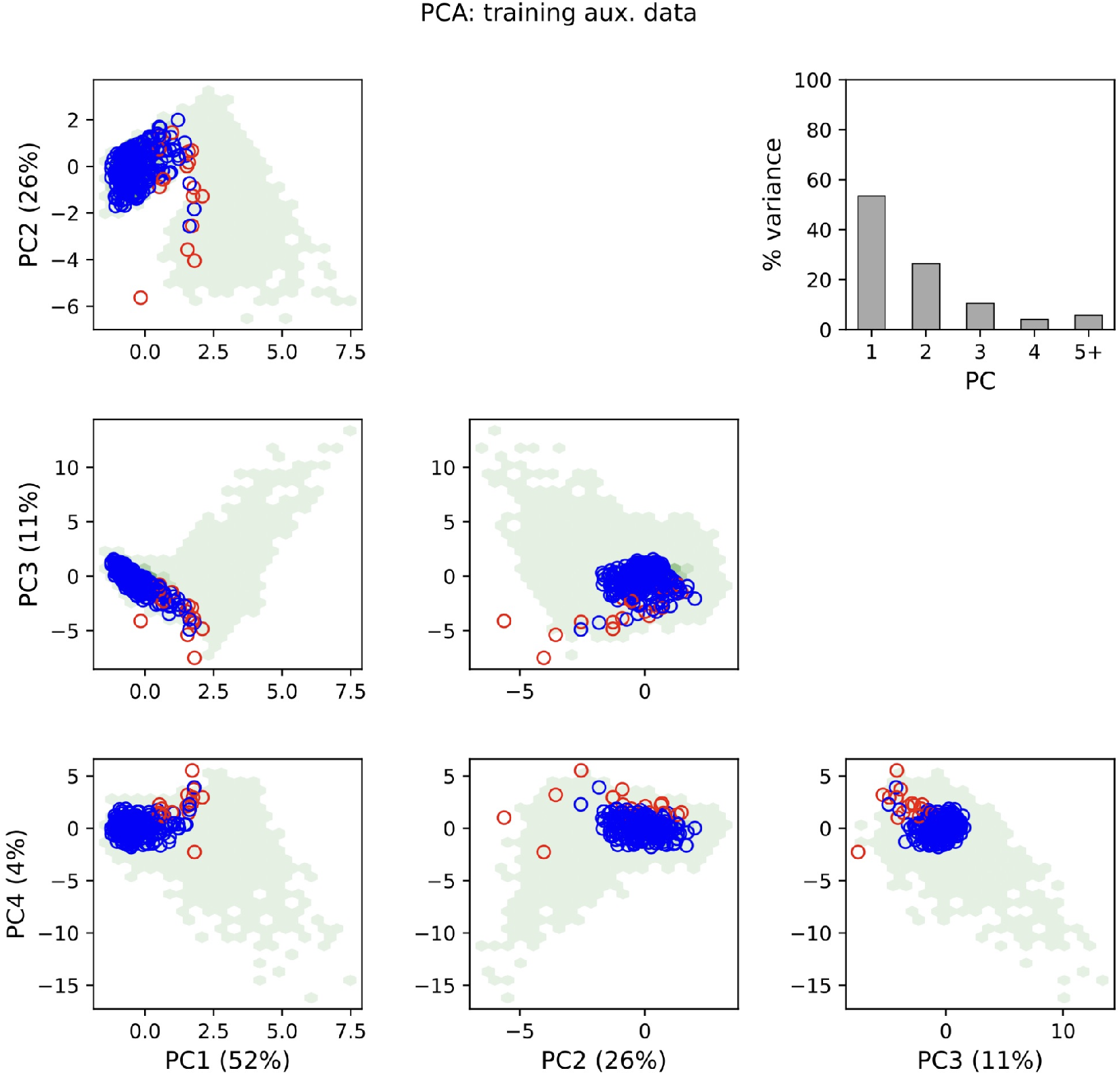
PCA plots for the auxiliary data tensor for data simulated with relative area size under Submodel 3. The blue and red circles correspond to a random subsample of size 250 of the summary statistics from the dataset simulated under a variant of Submodel 3 that accounts for relative area size. The red circles represent simulations identified as outliers relative to the training dataset generated under Submodel 3 without relative area size, whereas the blue circles represent the remaining simulations. In total, 7.2% of the samples were categorized as outliers. These outliers indicate a shift in distribution between the two datasets and may contribute to reduced CPI coverage under model misspecification.

**Figure S21:**
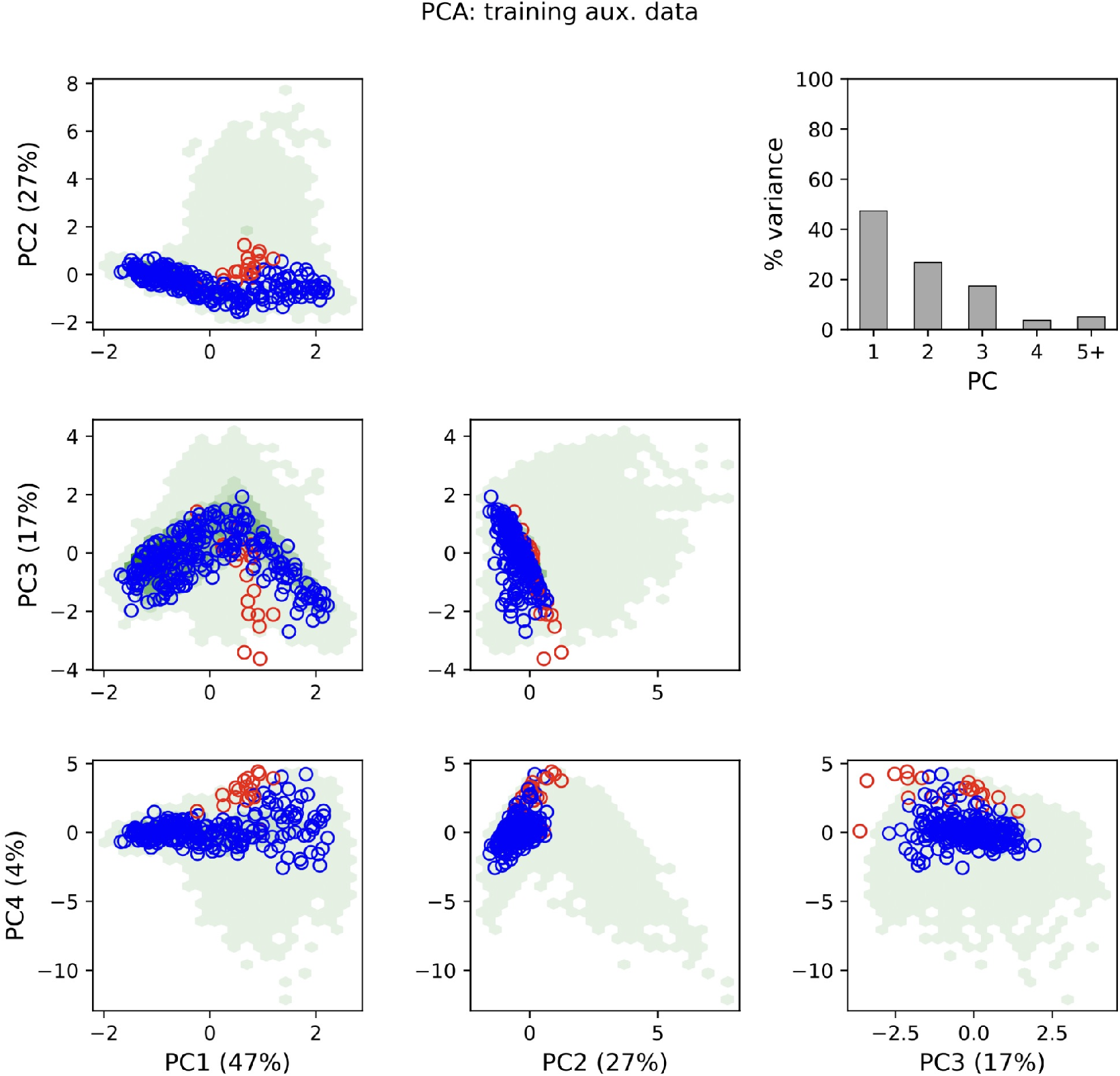
PCA plots for the auxiliary data tensor for data simulated with relative area size under Submodel 4. The blue and red circles correspond to a random subsample of size 250 of the summary statistics from the dataset simulated under a variant of Submodel 4 that accounts for relative area size. The red circles represent simulations identified as outliers relative to the training dataset generated under Submodel 4 without relative area size, whereas the blue circles represent the remaining simulations. In total, 8.4% of the samples were categorized as outliers. These outliers indicate a shift in distribution between the two datasets and may contribute to reduced CPI coverage under model misspecification.

**Figure S22:**
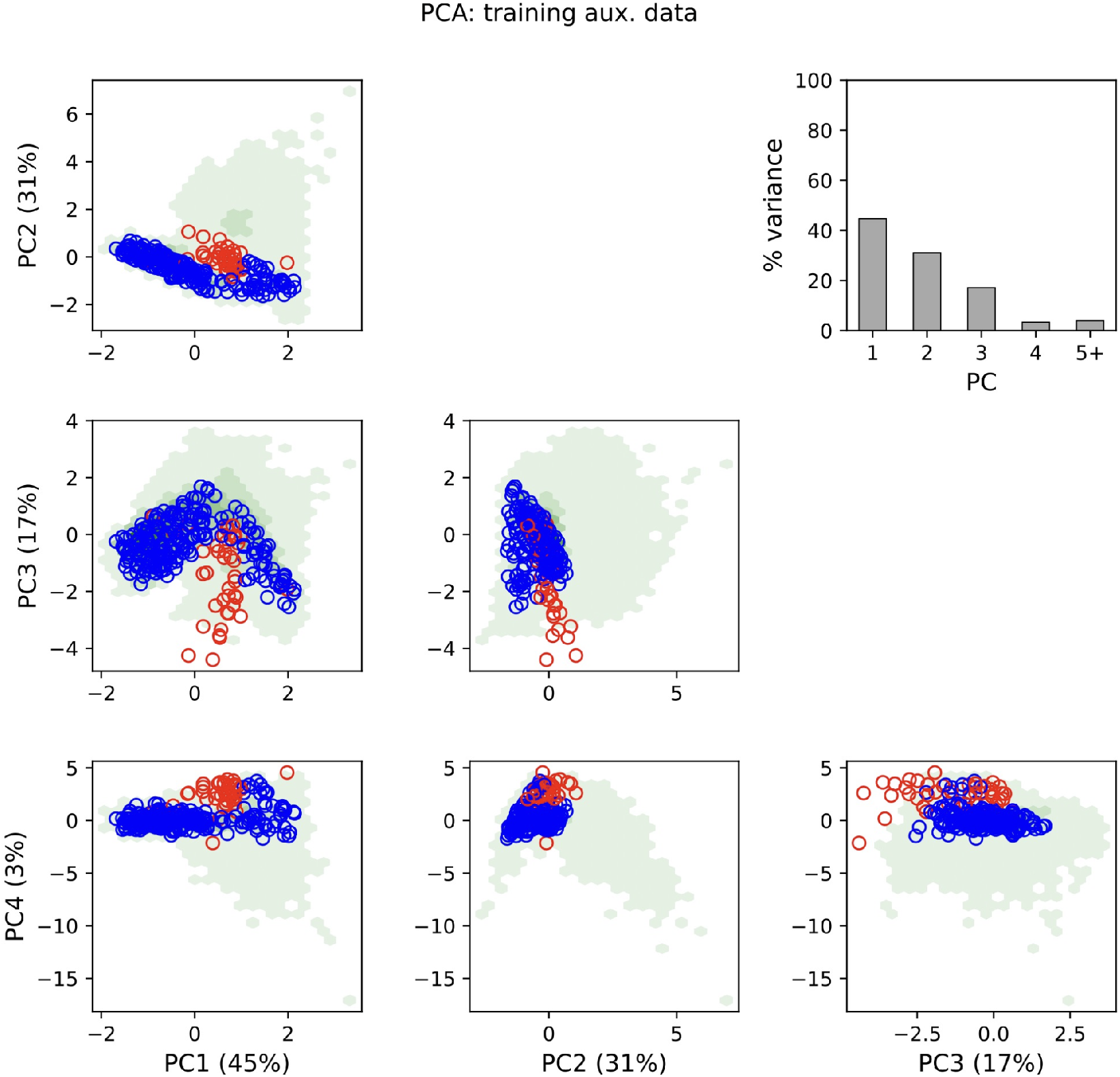
PCA plots for the auxiliary data tensor for data simulated with relative area size under Submodel 5. The blue and red circles correspond to a random subsample of size 250 of the summary statistics from the dataset simulated under a variant of Submodel 5 that accounts for relative area size. The red circles represent simulations identified as outliers relative to the training dataset generated under Submodel 5 without relative area size, whereas the blue circles represent the remaining simulations. In total, 16.8% of the samples were categorized as outliers. These outliers indicate a shift in distribution between the two datasets and may contribute to reduced CPI coverage under model misspecification.

**Figure S23:**
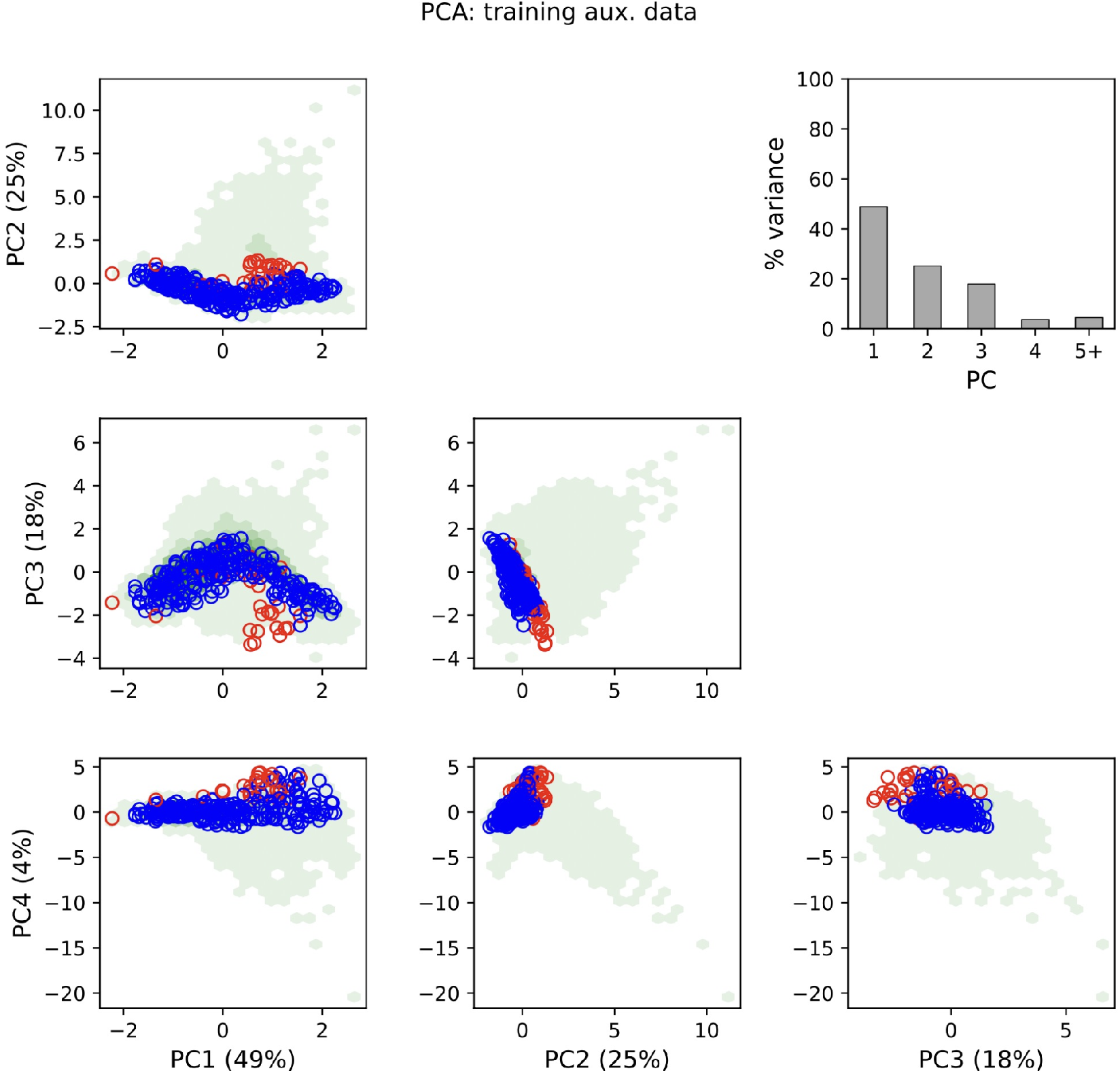
PCA plots for the auxiliary data tensor for data simulated with relative area size under Submodel 6. The blue and red circles correspond to a random subsample of size 250 of the summary statistics from the dataset simulated under a variant of Submodel 6 that accounts for relative area size. The red circles represent simulations identified as outliers relative to the training dataset generated under Submodel 6 without relative area size, whereas the blue circles represent the remaining simulations. In total, 12% of the samples were categorized as outliers. These outliers indicate a shift in distribution between the two datasets and may contribute to reduced CPI coverage under model misspecification.

**Figure S24:**
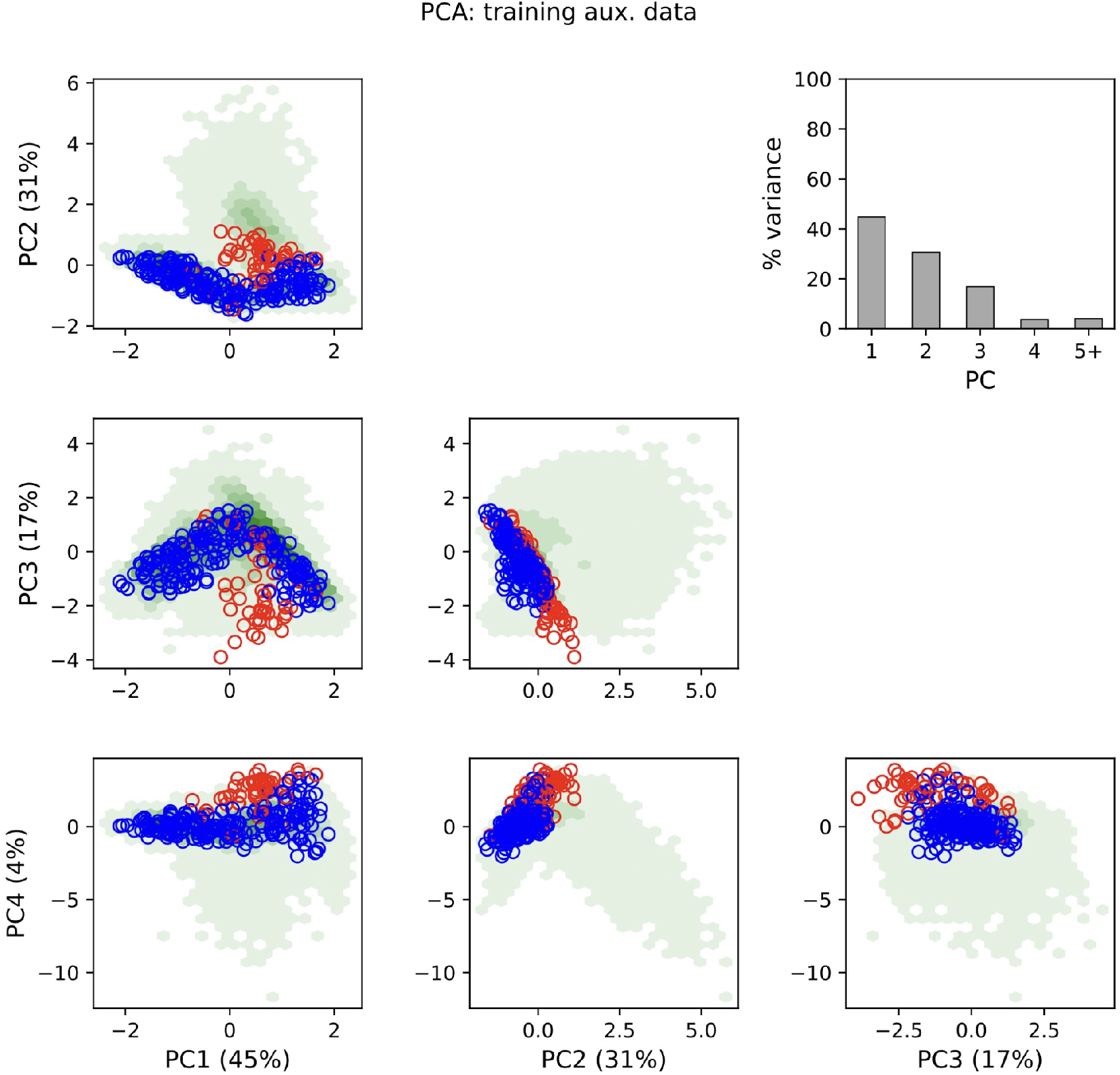
PCA plots for the auxiliary data tensor for data simulated with relative area size under Submodel 7. The blue and red circles correspond to a random subsample of size 250 of the summary statistics from the dataset simulated under a variant of Submodel 7 that accounts for relative area size. The red circles represent simulations identified as outliers relative to the training dataset generated under Submodel 7 without relative area size, whereas the blue circles represent the remaining simulations. In total, 20.4% of the samples were categorized as outliers. These outliers indicate a shift in distribution between the two datasets and may contribute to reduced CPI coverage under model misspecification.

## Phylogenies and biogeography of empirical systems

**Figure S25:**
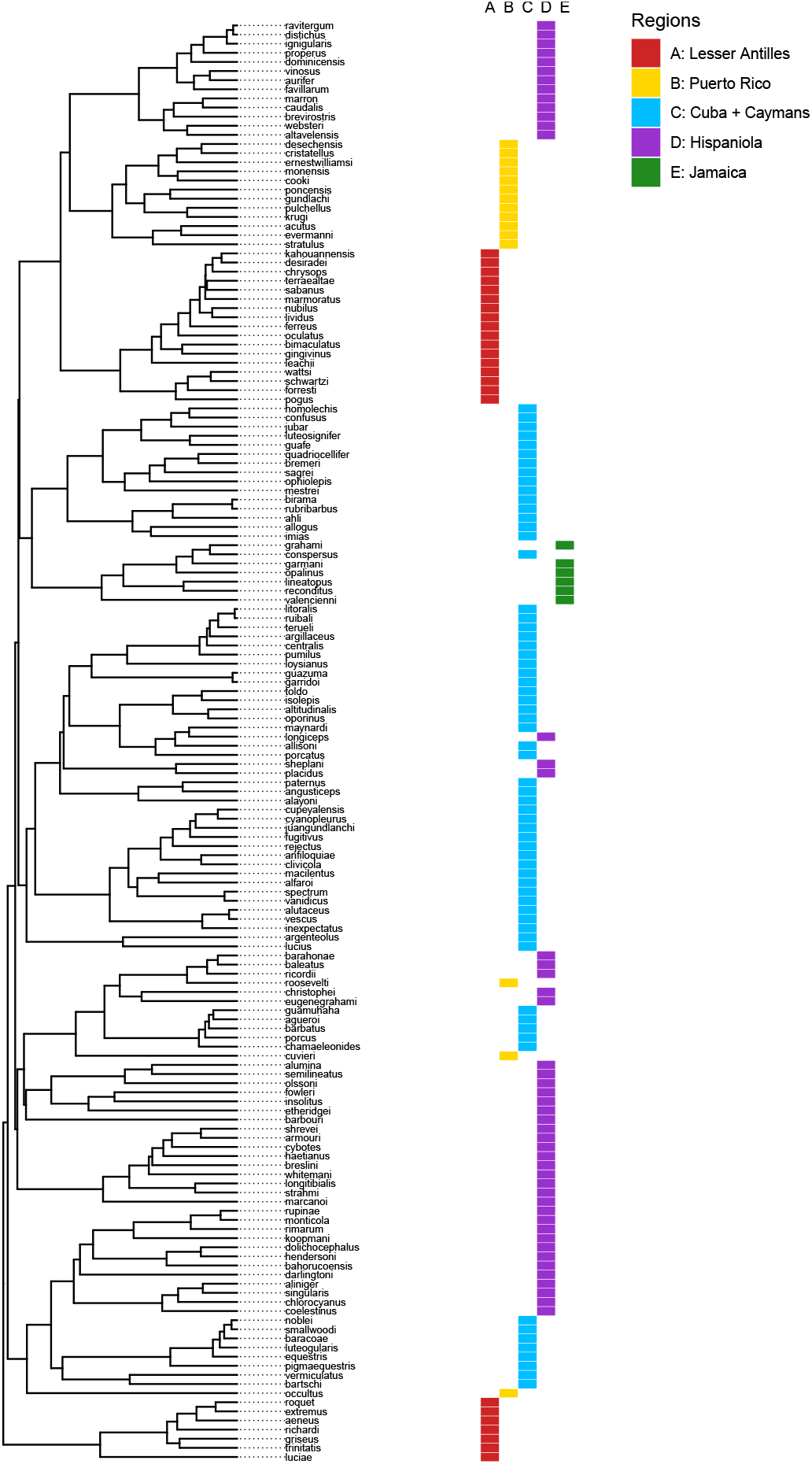
Phylogeny and species ranges for 158 *Anolis* lizard species inhabiting Caribbean islands. The phylogeny was subsampled from the tree produced by Poe et al. (2017). Ranges from Poe et al. (2017) were recoded for 5 island regions.

**Figure S26:**
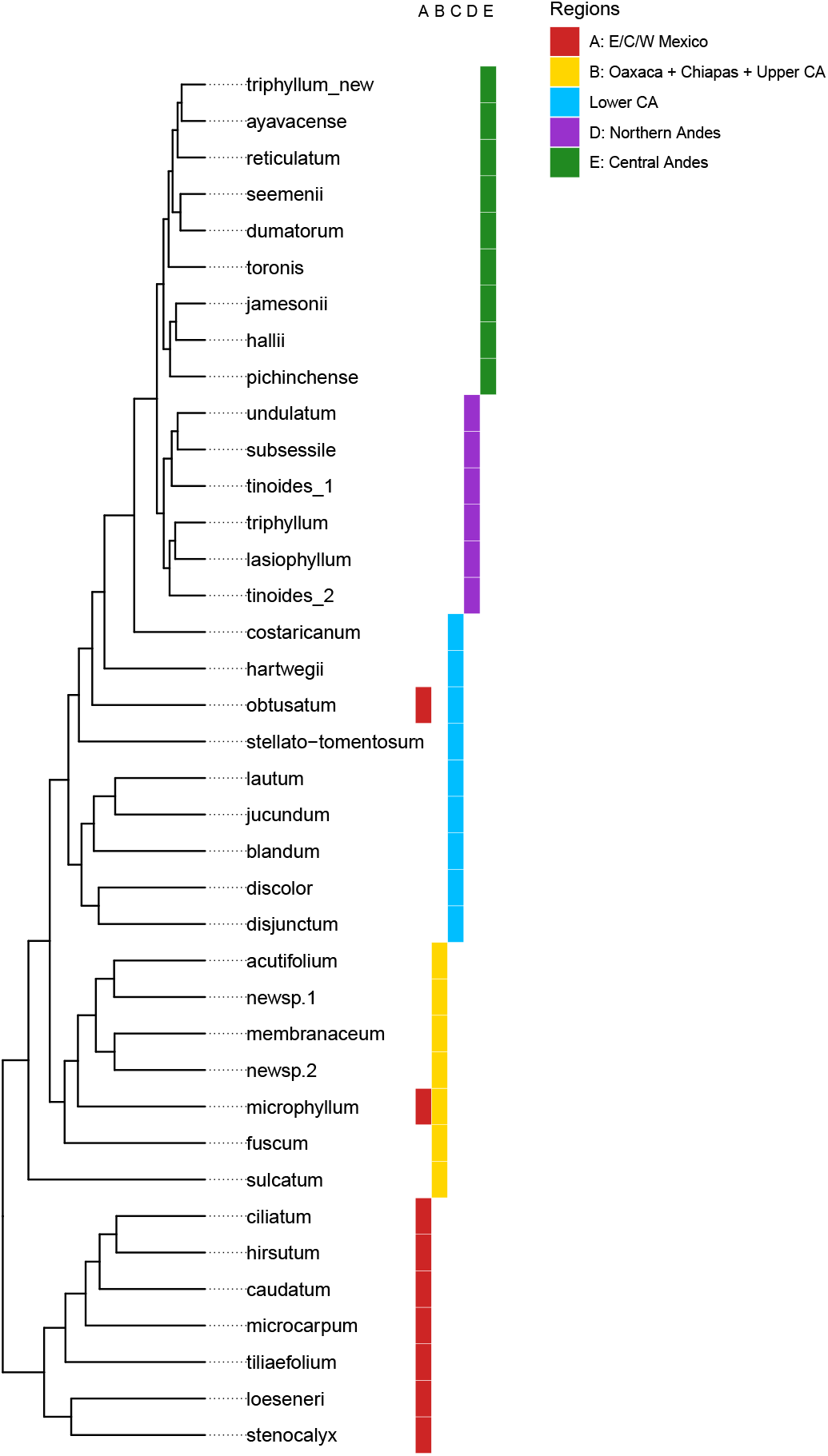
Phylogeny and species ranges for 38 *Oreinotinus* (a clade within *Viburnum*) plant species inhabiting neotropical cloud forests. The phylogeny was subsampled from the tree produced by Donoghue et al. (2022). Ranges from Donoghue et al. (2022) were recoded for 5 montane regions.

## Quality of trained networks for empirical parameter estimation

**Figure S27:**
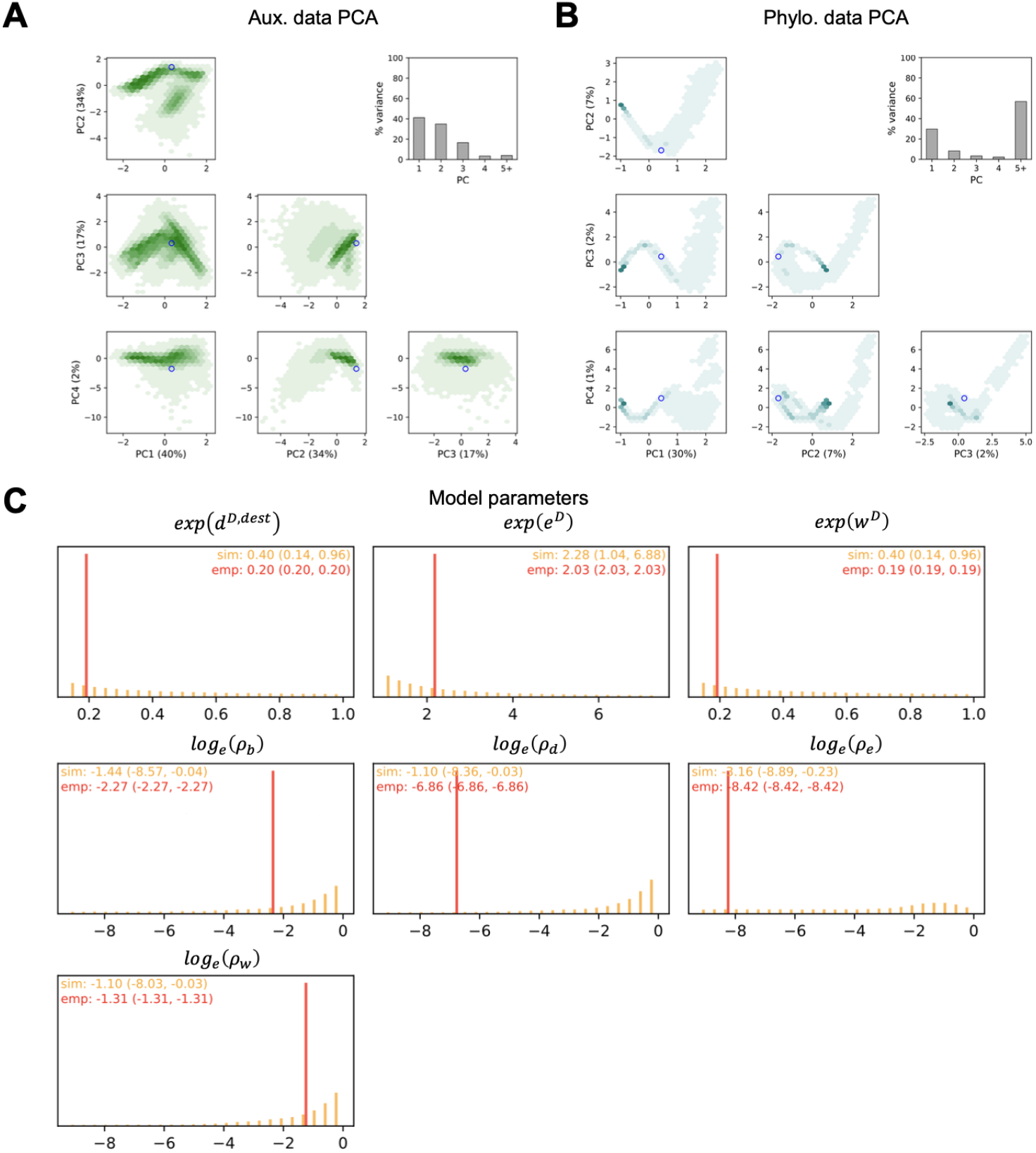
Network for *Anolis* analysis. PCA plots for the auxiliary data tensor (a) and phylogenetic data tensor (b) training sets. The empty circle represents the *Anolis* dataset in the PCA space. Histogram of Log-DDG model parameters in the training set (c). The red line represents the *Anolis* parameter estimates.

**Figure S28:**
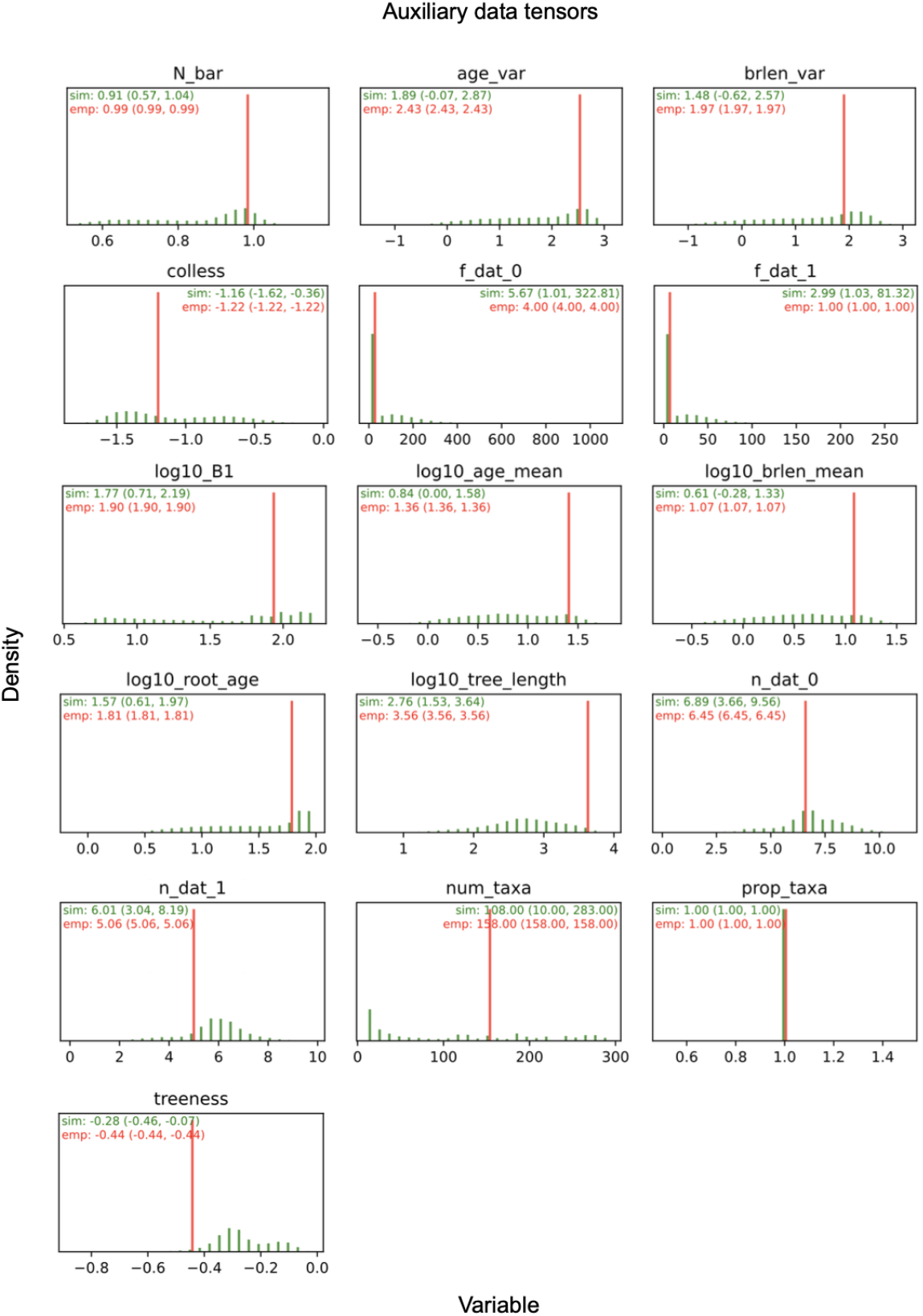
A panel of summary statistics (encoded as auxiliary data tensors) extracted from the inputted phylogeny and character data matrix for the *Anolis* tree (red bars) and the training dataset (green bars) from the best-fitted model for *Anolis* (Submodel 7; green bar). The numbers on the green intervals represent both 2.5% and 97.5% quantiles from the simulated (training) data. See Table S1 for the description of these summary statistics.

**Figure S29:**
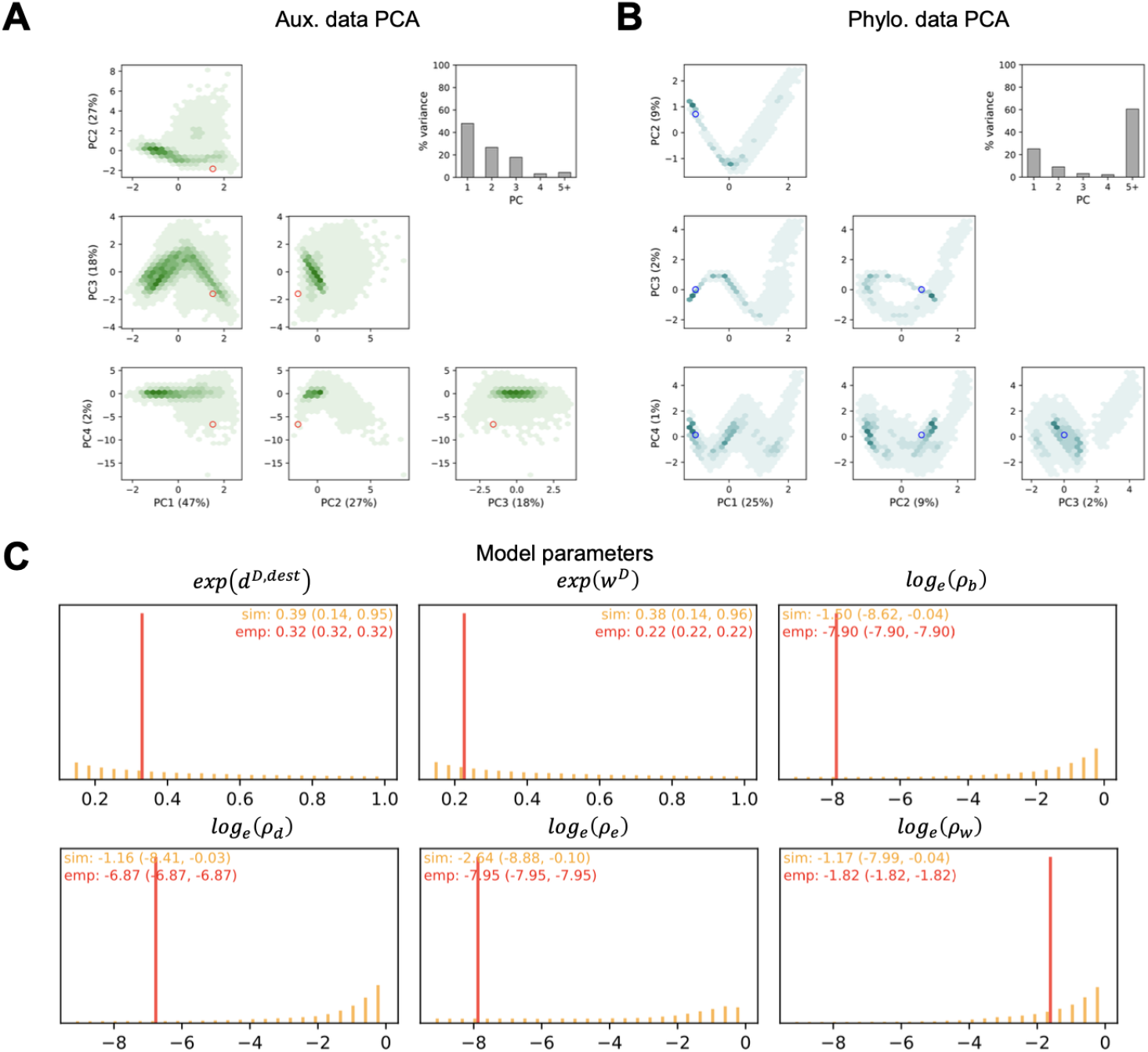
Network for *Viburnum* analysis. PCA plots for the auxiliary data tensor (a) and phylogenetic data tensor (b) training sets. The empty circle represents the *Viburnum* dataset in the PCA space. Histogram of Log-DDG model parameters in the training set (c). The red line represents the *Viburnum* parameter estimates.

**Figure S30:**
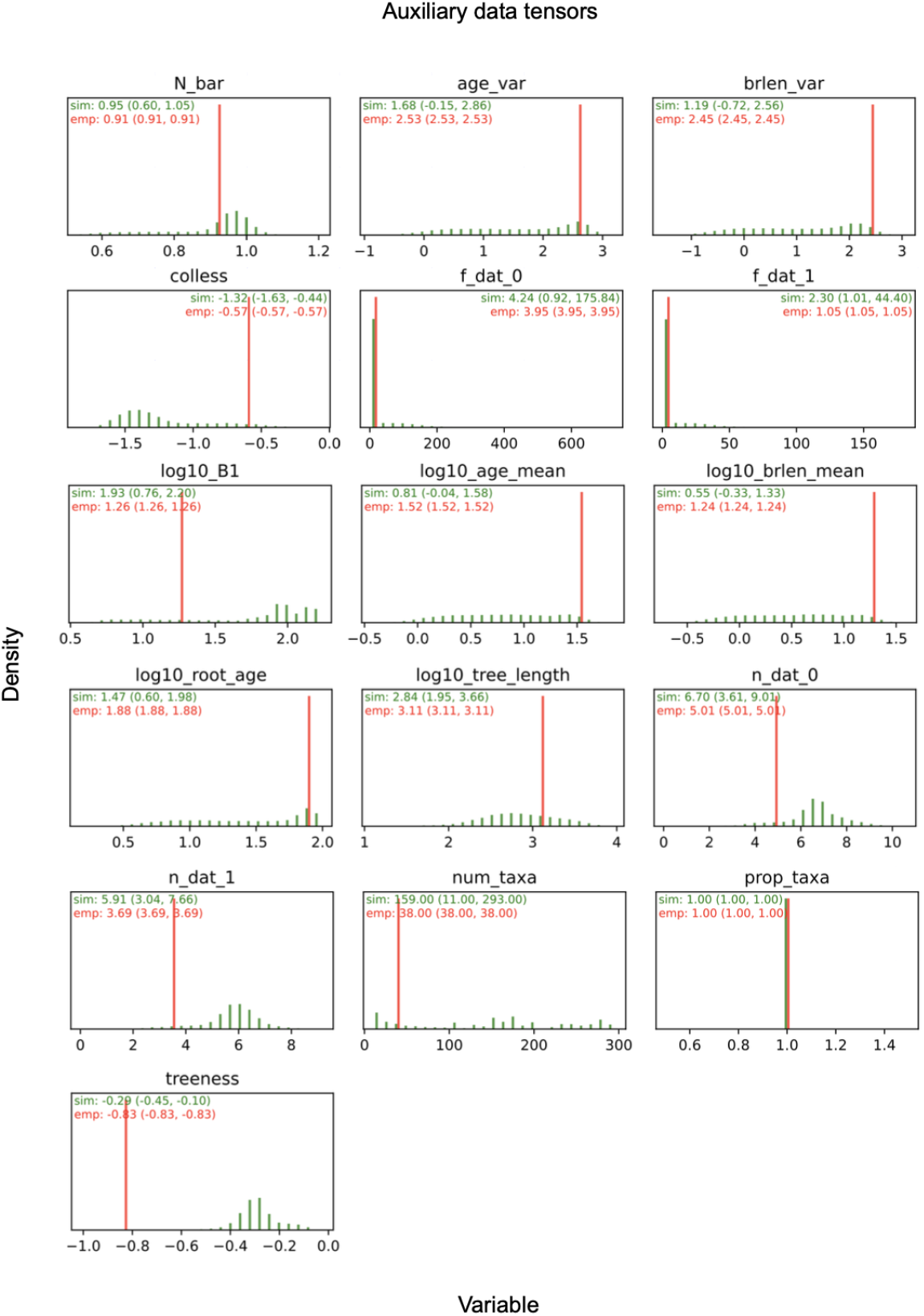
A panel of summary statistics (encoded as auxiliary data tensors) extracted from the inputted phylogeny and character data matrix for the *Viburnum* tree (red bars) and the training dataset (green bars) from the best-fitted model for *Viburnum* (Submodel 6; green bar). The numbers on the green intervals represent both 2.5% and 97.5% quantiles from the simulated (training) data. See Table S1 for the description of these summary statistics.

## Runtime benchmarking across different numbers of regions

**Figure S31:**
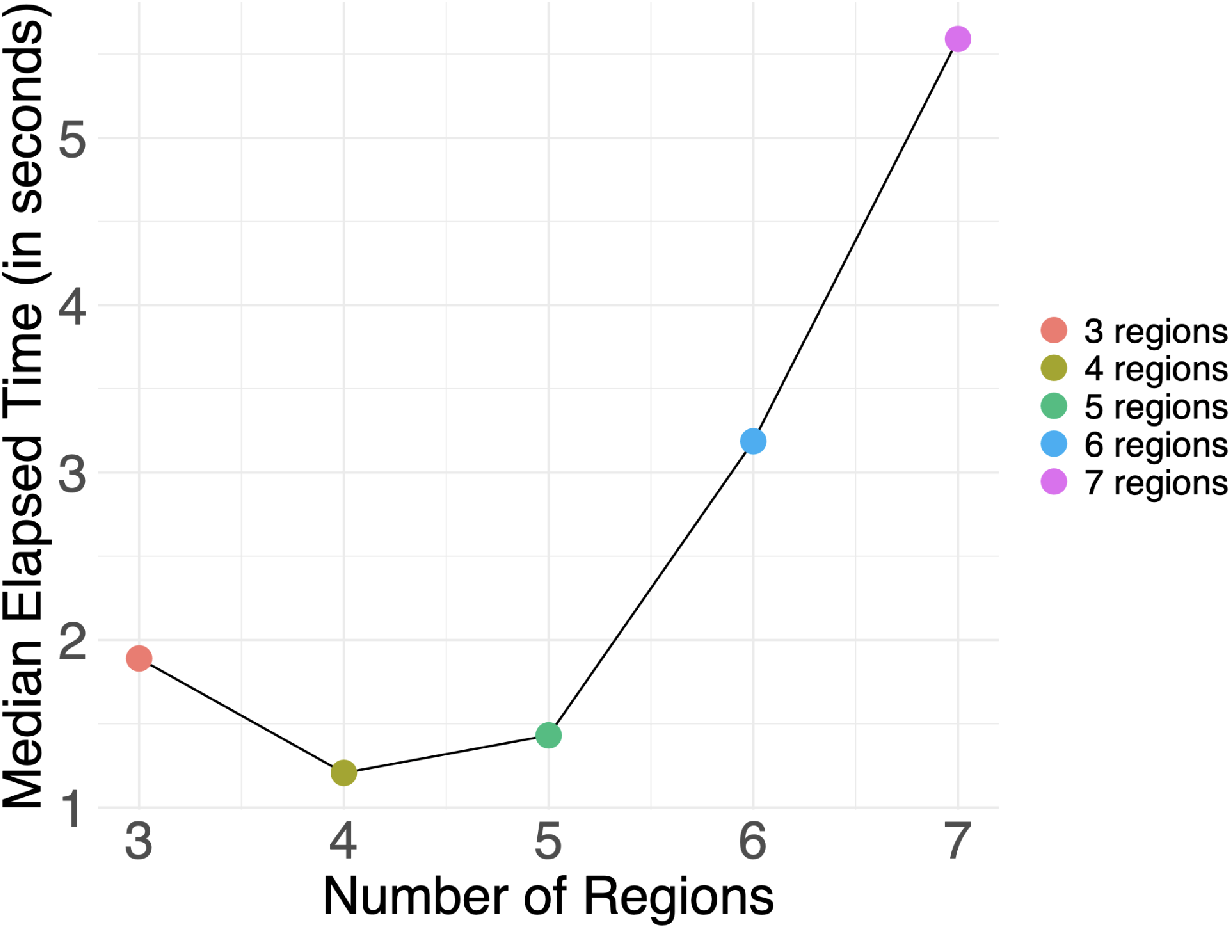
Median runtime (seconds) for datasets simulated under Submodel 7 with 3, 4, 5, 6, and 7 regions. For each experiment, we simulated 1,000 trees conditioned on a fixed number of extant taxa (*n* = 150). For each tree, we fixed the starting region of the root lineage and simulated under the parameter values *ρ*_*w*_ = 1.0, *ρ*_*d*_ = 0.1, *ρ*_*b*_ = 0.1, *ρ*_*e*_ = 0.2, *w*^*D*^ = −0.6, *e*^*D*^ = 0.5, *d*^*D,src*^ = 0, *d*^*D,dest*^ = −0.5, *b*^*D*^ = 0.

## Julia simulator validation using GeoSSE model

**Figure S32:**
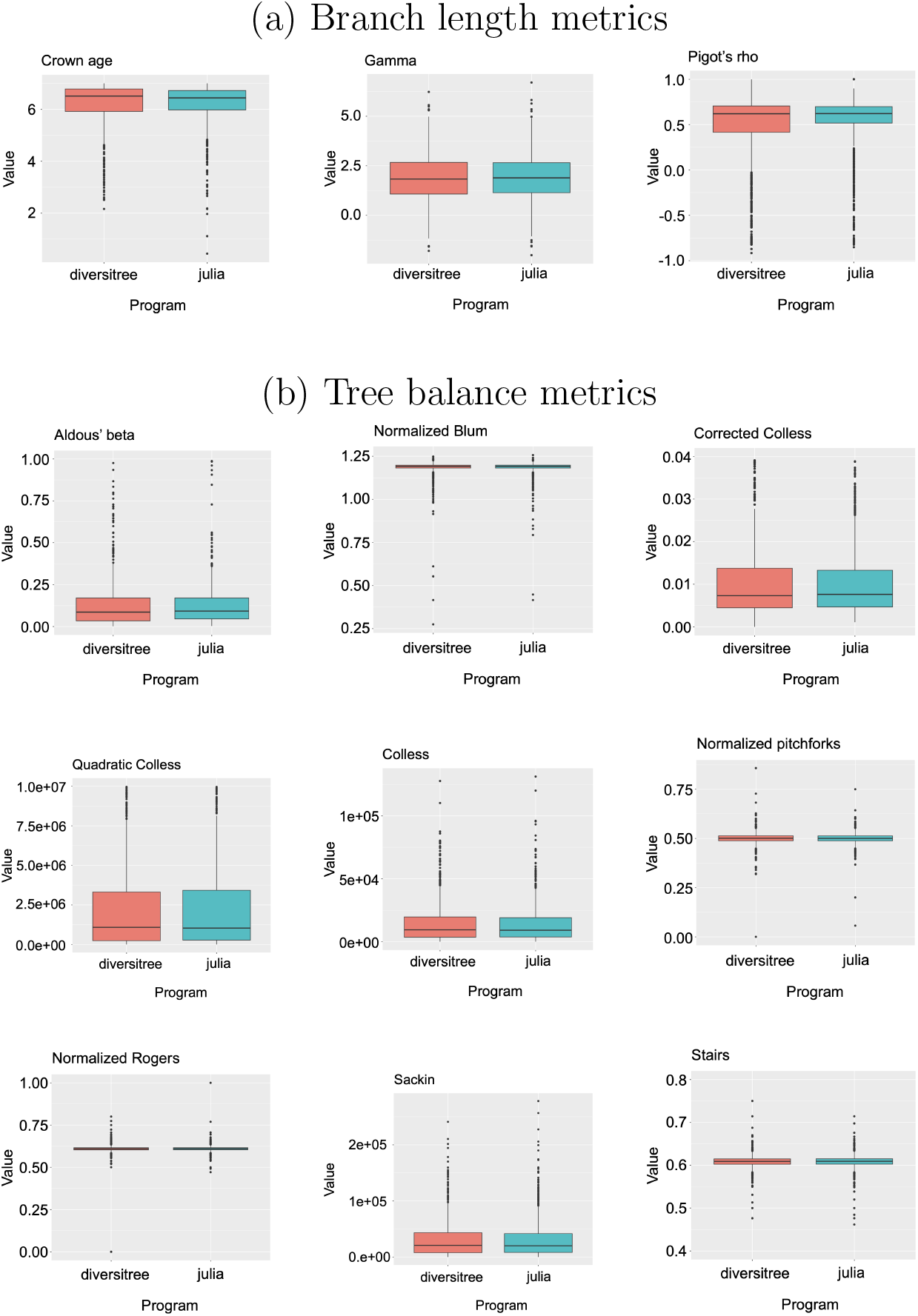
Comparison of tree balance and branch length metrics between our Julia script and diversitree for simulating GeoSSE trees without model misspecification. For each program, we simulated 1,000 trees using the following parameters: 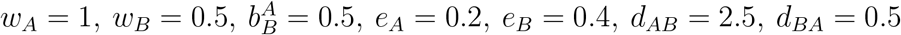. Each row shows related metrics as indicated by the bold row headings.

**Figure S33:**
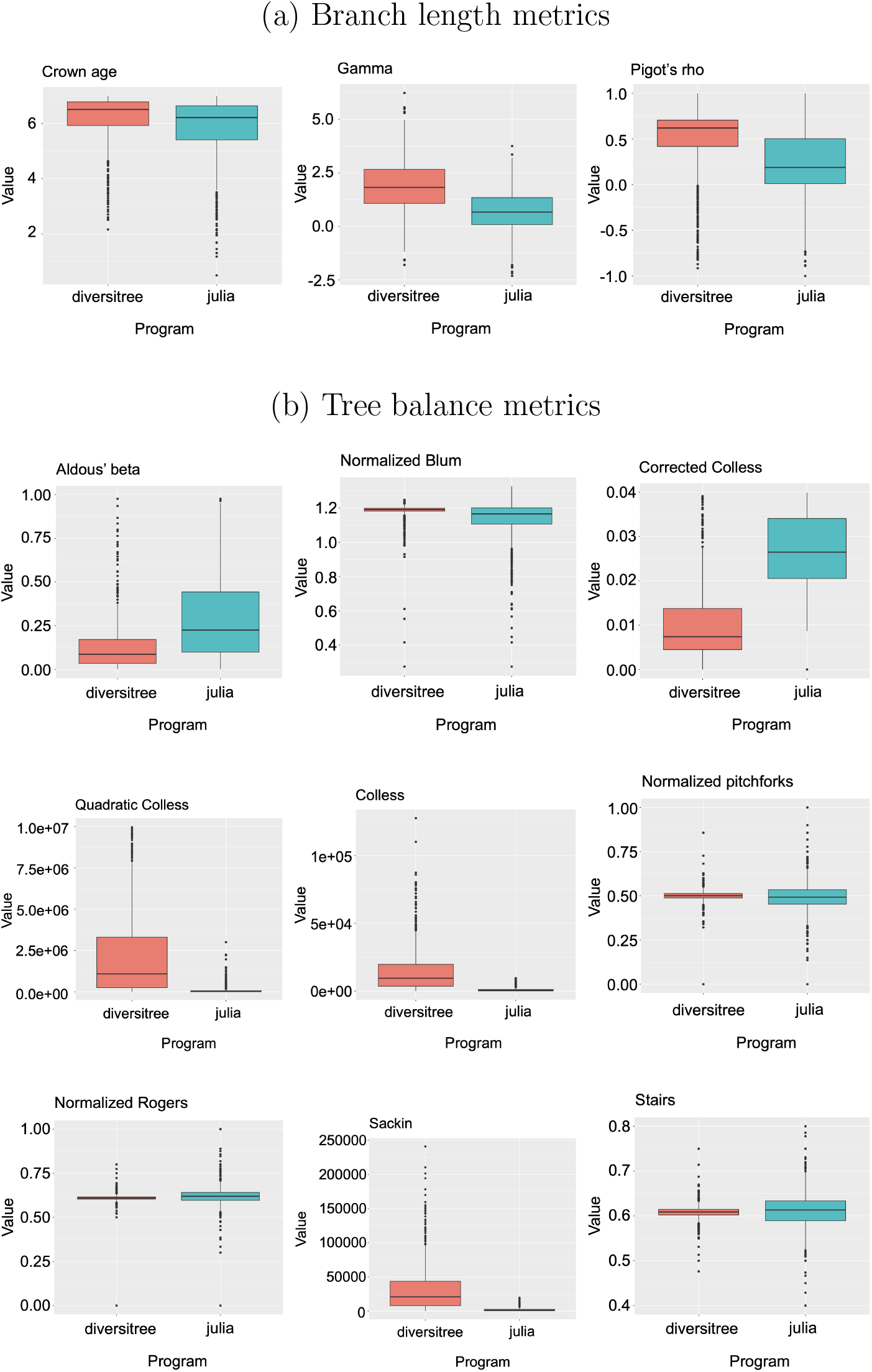
Comparison of tree balance and branch length metrics between our Julia script and diversitree for simulating GeoSSE trees with intentional model misspecification, where the speciation rates in both regions are doubled for Julia simulations to cause the results to differ. We simulated 1000 trees using the following parameters: 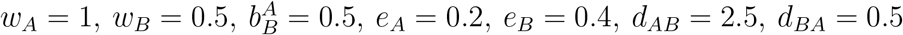.

## Neural network architecture

**Figure S34:**
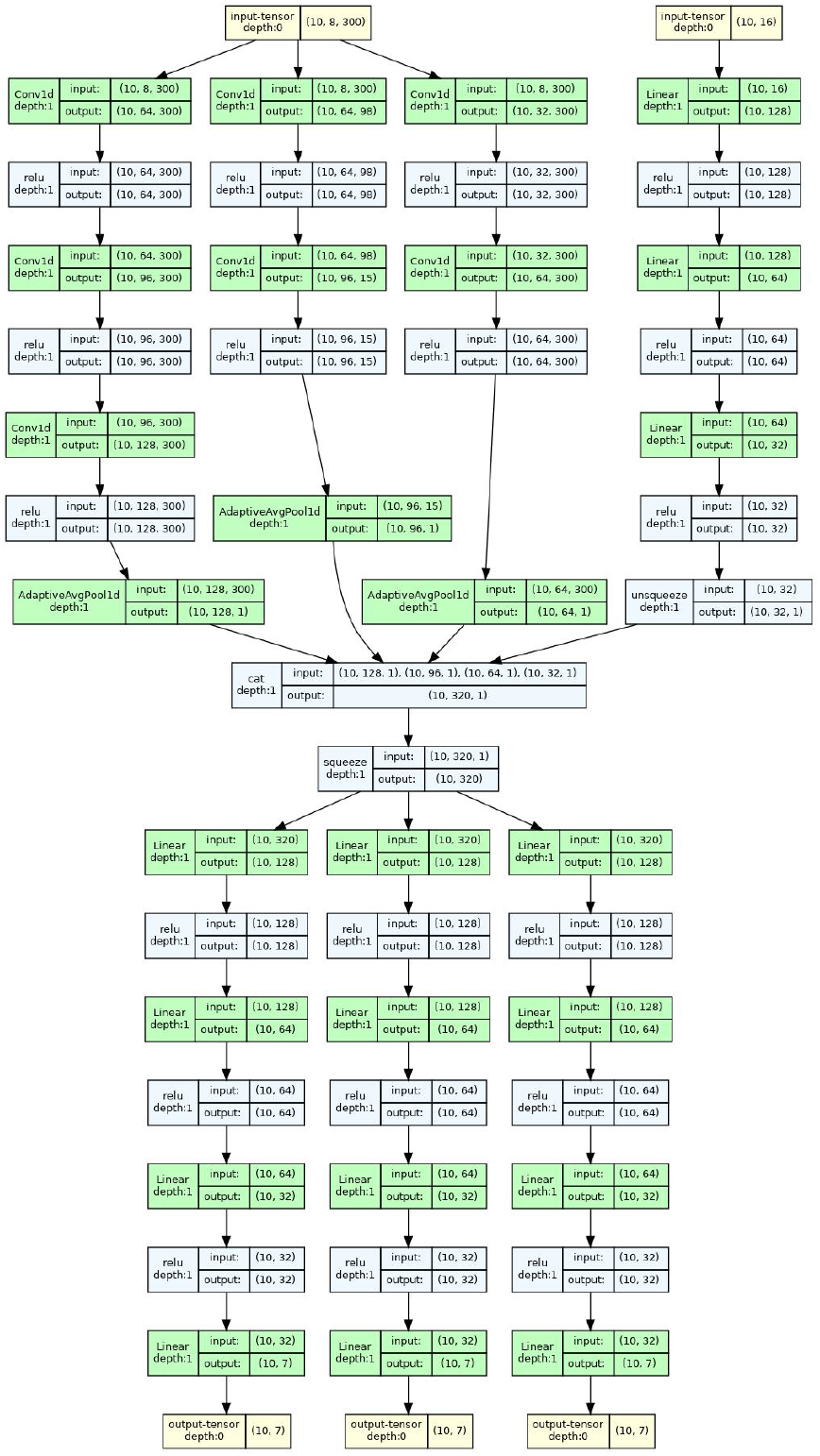
Neural network architecture used for parameter estimation tasks with phyddle. This architecture corresponds to Log-DDG submodel 7, with seven outputs corresponding to the four base rate parameters (*ρ*_*e*_, *ρ*_*w*_, *ρ*_*d*_, *ρ*_*b*_) plus three diversitydependent effect parameters (*e*^*D*^, *w*^*D*^, *d*^*D,dest*^). All other submodels used the same architecture, except with fewer outputs for those effect parameters that were fixed to 0 (no effect). Green boxes correspond to nodes, blue boxes correspond to activation functions, yellow boxes correspond to inputs and outputs. Arrows show how information is passed between layers, with numbers in parentheses describing tensor shapes.

**Figure S35:**
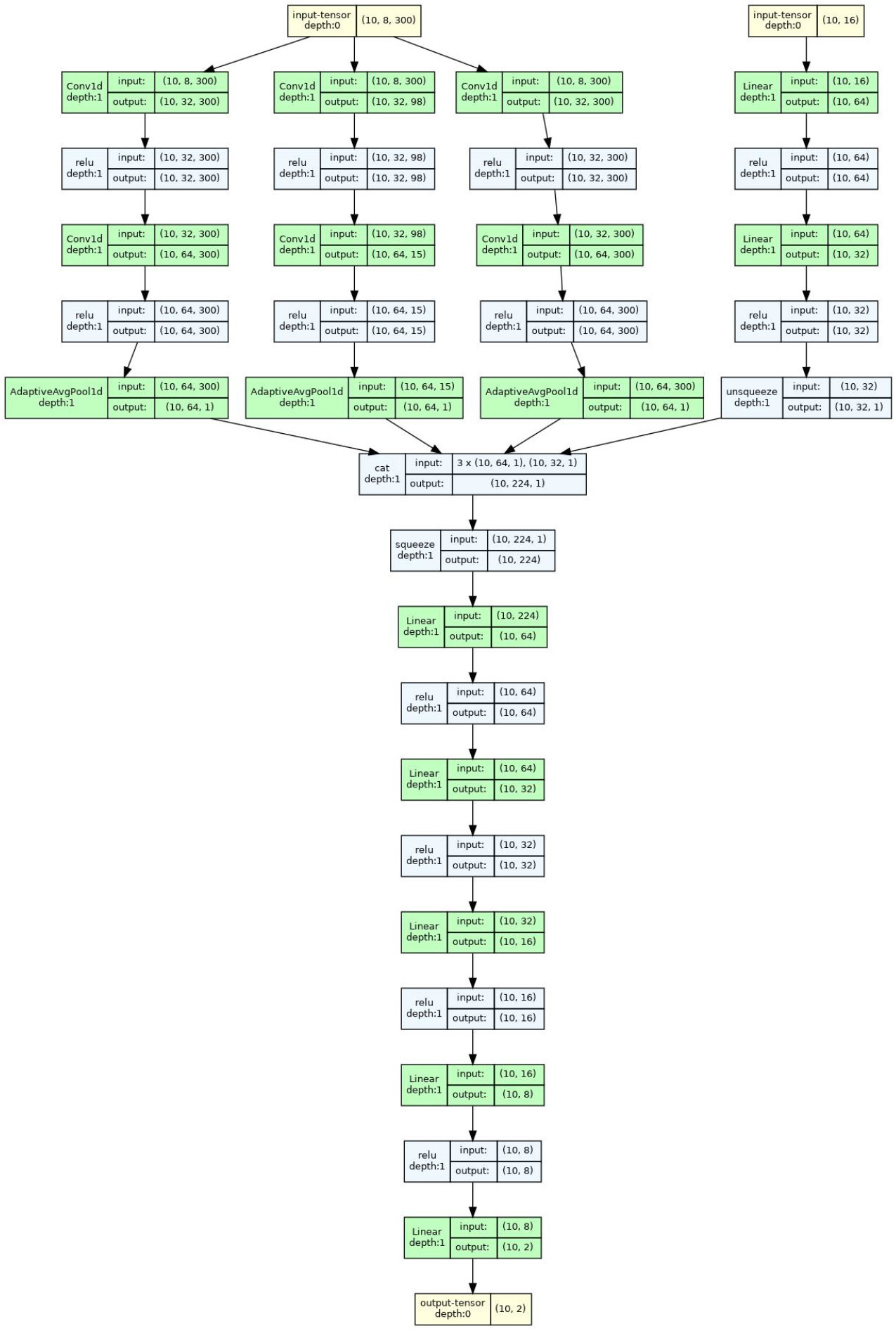
Neural network architecture used for model selection tasks with phyddle. While this particular example was used for model selection for diversity-dependent dispersal (to test whether *d*^*D,dest*^ = 0 or *d*^*D,dest*^ ≠ 0), the exact same architecture was used in separately trained networks for model selection for diversity-dependent within-region speciation (*w*^*D*^) and extinction (*e*^*D*^). Green boxes correspond to nodes, blue boxes correspond to activation functions, yellow boxes correspond to inputs and outputs. Arrows show how information is passed between layers, with numbers in parentheses describing tensor shapes.

